# The two-component system ChvGI maintains cell envelope homeostasis in *Caulobacter crescentus*

**DOI:** 10.1101/2022.01.18.476748

**Authors:** Alex Quintero-Yanes, Aurélie Mayard, Régis Hallez

## Abstract

Two-component systems (TCS) are often used by bacteria to rapidly assess and respond to environmental changes. The ChvG/ChvI (ChvGI) TCS conserved in α-proteobacteria is known for regulating expression of genes related to exopolysaccharide production, virulence and growth. The sensor kinase ChvG autophosphorylates upon yet unknown signals and phosphorylates the response regulator ChvI to regulate transcription. Recent studies in *Caulobacter crescentus* showed that *chv* mutants are sensitive to vancomycin treatment and fail to grow in synthetic minimal media. In this work, we identified the osmotic imbalance as the main cause of growth impairment in synthetic minimal media. We also determined the ChvI regulon and found that ChvI regulates cell envelope architecture by controlling outer membrane, peptidoglycan assembly/recycling and inner membrane proteins. In addition, we found that ChvI phosphorylation is also activated upon antibiotic treatment with vancomycin. We also challenged *chv* mutants with other cell envelope related stress and found that treatment with antibiotics targeting transpeptidation of peptidoglycan during cell elongation impairs growth of the mutant. Moreover, these antibiotics activate expression of the *chvIG-hprK* operon. Finally, we observed that the sensor kinase ChvG relocates from a patchy-spotty distribution to distinctive foci after transition from complex to synthetic minimal media. Interestingly, this pattern of (re)location has been described for proteins involved in cell growth control and peptidoglycan synthesis upon osmotic shock. Overall, our data support that the ChvGI TCS is mainly used to monitor and respond to osmotic imbalances and damages in the peptidoglycan layer to maintain cell envelope homeostasis.

**Importance:** The cell envelope is the first barrier protecting cells from harsh environmental conditions, such as temperature, pH, oxidative and osmotic imbalances. It is also an obstacle for the intake of antibiotics targeting essential cellular processes. Therefore, molecular components and systems responding to cell envelope stress and maintaining cell envelope homeostasis are important targets for drug therapy. Here we show that the two-component system ChvGI, highly conserved in free-living and pathogenic *α*-proteobacteria, is activated upon osmotic upshift and treatment with antibiotics targeting peptidoglycan synthesis to activate the transcription of multiple genes involved in cell envelope homeostasis in *Caulobacter crescentus*. We also show that the kinase sensor ChvG displays a dynamic localisation pattern that changes depending on osmotic imbalance. To our knowledge this is the first two-component system reported to change his cellular localisation upon environmental stress.

## Introduction

Two-component systems (TCS) equip cells with a rapid sensing and response mechanism to optimise survival in changing and stressful environments. The first component of canonical TCS, a sensor histidine kinase (HK), autophosphorylates upon input signal detection on a conserved histidine residue using the gamma-phosphoryl group of ATP. Thereafter, the phosphoryl group is transferred from the HK on a conserved aspartate residue of the second TCS component, a cognate response regulator (RR). In most cases, the RR harbours an output domain that binds to DNA upon phosphorylation to either activate or repress transcription of target genes (Stock *et* al., 2000; Goulian, 2010; Capra & Laub, 2012). On the other hand, HKs can also exert specific phosphatase activities on their RRs to turn off the response in absence of the input signal (Gao and Stock, 2009).

The Chv (**ch**romosomal **v**irulence factor) TCS conserved in *α*-proteobacteria and composed of the HK ChvG and the RR ChvI, was first reported as a pathogenic regulator in *Agrobacterium tumefaciens* responding to acid stress (Mantis & Winans, 1993; Li *et al.,* 2002). A study in *Sinorhizobium meliloti* showed that the ChvG homologue ExoS is negatively regulated by the periplasmic protein ExoR (Chen *et al.,* 2008). Further research in *A. tumefaciens* showed that low pH triggers ExoR proteolysis to derepress ChvGI activity (Wu *et al.,* 2012). While unlike other *α*-proteobacteria, ExoR orthologs are absent in the aquatic free-living bacterium *Caulobacter crescentus*, the ChvGI-dependent response to acidic conditions seems nonetheless conserved in *C. crescentus* (Stein *et al.,* 2021). Similarly, the ChvI homologue BvrR of the intracellular pathogen *Brucella abortus* is phosphorylated in combination of low pH and nutrient depletion conditions, which mimics post-infection conditions (Altamirano-Silva *et al.,* 2018).

In *C. crescentus*, ChvGI activates the expression of the small non-coding RNA (sRNA) ChvR when exposed to acidic stress or DNA damage with mitomycin C, or when cultured in synthetic minimal media (Frölich *et al.,* 2018). Once produced, ChvR subsequently inhibits translation of the TonB-dependent receptor (TBDR) ChvT (Frölich *et al.,* 2018). Interestingly, inactivating of *chvR* or *chvIG* sensitises *C. crescentus* cells to vancomycin treatment whereas a *chvT* mutant makes them resistant to this antibiotic, which suggests that vancomycin passes through the outer membrane via the TBDR ChvT to reach the periplasm (Vallet *et al.,* 2020). In addition, a *chvIG* knock-out (*ΔchvIG*) mutant could not propagate in synthetic minimal media when cells were inoculated at low density, while inactivating *chvT* in a *ΔchvIG* background partially restored growth (Stein *et al.,* 2021). Although *chvIG* mutants are sensitive to acid stress when grown in minimal media, the primary cause for the growth defect has not been determined.

ChvGI is also known in *C. crescentus* to coordinate its regulation with another TCS, NtrYX, which is under control of the periplasmic protein NtrZ (Stein *et al.,* 2021). NtrZ has been only described in *C. crescentus*, while the HK NtrY and RR NtrX are known for regulating multiple processes in *α*-proteobacteria such as nitrogen metabolism, motility, virulence and cell envelope integrity (Pawlowski *et al.,* 1991; Carrica *et al.,* 2012; Wang, D *et al.,* 2013; Lemmer *et al.,* 2020). Stein *et al*. (2021) showed that both ChvI and NtrX networks significantly overlap and these RRs have opposite regulatory functions in minimal media. For instance, while ChvI acts as a positive regulator of growth, NtrX inhibits growth. Interestingly, repression of growth in synthetic media is caused by unphosphorylated NtrX, so that inactivating *ntrX* in a *ΔchvI* background also partially restored growth. Actually, NtrY acts as a phosphatase over the phosphorylated NtrX (NtrX∼P), while NtrZ inhibits NtrY to presumably maintain high NtrX∼P levels.

Here we show that *chvIG* mutants are greatly impacted due to osmotic imbalances in minimal media, but also in complex media supplemented with osmolytes. The Although the general stress response (GSR) sigma factor SigT was shown to be activated upon osmotic imbalance (Alvarez-Martinez *et al*., 2007), we showed that *chvIG* mutants are much more sensitive to lower hypertonic conditions than *sigT* mutant. In agreement with previous data (Stein *et al.,* 2021), deletion of *chvT* and *ntrX* restored growth in hypertonic conditions. We provide a ChvI regulon, using ChIP-seq with polyclonal anti-ChvI antibodies and RNA-seq, which unveiled new targets related to peptidoglycan (PG) synthesis and recycling. We also showed that (i) ChvG-dependent phosphorylation of ChvI and (ii) expression of the *chvIG-hprK* operon is induced upon osmotic shock or treatment with antibiotics targeting the PG. Finally, we showed that upon these stressful conditions, the HK ChvG relocates from a patchy-spotty pattern typically displayed by PG-related proteins to midcell. Overall, our results support a critical role for the ChvGI TCS in maintaining cell morphology and cell envelope homeostasis upon stress.

## Methods

### Bacterial strains and growth conditions

Strains, plasmids and oligonucleotides used in this study are listed in supplementary tables (Tables S 1, 2 and 3). Plasmid construction details are presented in supplementary methods. *E. coli* strains were grown aerobically in either LB (broth) sigma or LB + 1.5% agar at 37 °C. All *C. crescentus* strains in this study are derived of the NA1000 wild-type strain, and growth was achieved at 30 °C in aerated conditions using either complex medium Peptone Yeast Extract (PYE) or synthetic media supplemented with glucose (M2G or M5GG) as already described in (Ronneau et al., 2016). M2G and M5GG were prepared using M2 (12.25 mM Na_2_HPO_4_, 7.75 mM KH_2_PO_4_, 9.35 mM NH_4_Cl) and M5 (10 mM PIPES pH 7.2, 1 mM NaCl, 1 mM KCl, 0.37 mM Na_2_HPO_4_, 0.23 mM KH_2_PO_4_) salts, respectively, and both supplemented with (0.5 mM MgSO_4_, 0.5 mM CaCl_2_, 0.01 mM FeSO_4_, 0.2% glucose). 1 mM glutamate sodium was added to make M5GG. Modified M2G 0% (Na^+^, K^+^), M2G 25% (Na^+^, K^+^) and M2G 50% (Na^+^, K^+^) were prepared without Na_2_HPO_4_ and KH_2_PO_4_ or either 50% or 25% of the Na_2_HPO_4_ and KH_2_PO_4_ concentrations in M2G, respectively. *C. crescentus* was grown on plates using either PYE or M2G supplemented with 1.5% agar at 30 °C. Growth was monitored by measuring absorbance at OD_660_ in liquid cultures using an automated plate reader (Biotek, Epoch 2) with continuous shaking at 30 °C. Gene expression under the control of inducible promoter (P*_xylX_*) was induced either with 0.1 % D(+)-xylose (Sigma). Expression of enhanced green fluorescent protein (*egfp*) and monomeric cherry (*mCherry*) protein fusions was induced in fresh exponentially growing cultures (OD_660_ ∼ 0.1) for 1 h and 4 h. Generalized transduction was performed with phage ϕCr30 according to the procedure described in (Ely, 1991).

Antibiotics for *E. coli* were used with the following final concentrations (*μ*g ml^-1^, in liquid/ solid medium) ampicillin (50/100), kanamycin (30/50), chloramphenicol (30/20), and oxytetracycline (12.5/12.5) for *C. crescentus* A22 (2.5/5) cefsulodin (80/80), cephalexin (15/15), mecillinam (100/100), moenomycin (0.5/1), kanamycin (5/20), oxytetracycline (1/2.5), polymyxin B (5/12.5), vancomycin (10/10) where appropriate. Plasmid delivery into *C. crescentus* was achieved by either bi- or tri-parental mating using *E. coli* S17-1 and *E. coli* MT607 as helper strains, respectively. In-frame deletions were created by using the pNPTS138-derivative plasmids as follows. Integration of the plasmids in the *C. crescentus* genome after single homologous recombination were selected on PYE plates supplemented with kanamycin. Three independent recombinant clones were inoculated in PYE medium without kanamycin and incubated overnight at 30 °C. Then, dilutions were spread on PYE plates supplemented with 3% sucrose and incubated at 30 °C. Single colonies were picked and transferred onto PYE plates with and without kanamycin. Finally, to discriminate between mutated and wild-type loci, kanamycin-sensitive clones were tested by PCR on colony using locus-specific oligonucleotides.

### Spotting assays

Ten-fold serial dilutions (in PYE) were prepared in 96-well plates from 5 ml cultures in standard glass tubes grown overnight at 30 °C in the corresponding media. Cells were then spotted on plates, were incubated at 30 °C for two-to-three days and pictures were taken. Cells in assays including strains with mutations in *ΔntrX* were normalised to OD_660_ since these mutations lead to growth defects in PYE.

### β-galactosidase assays

*β*-galactosidase assays were performed as already described in Beaufay et al., 2015. Briefly, overnight saturated cultures of *Caulobacter* cells harbouring *lacZ* reporter plasmids were diluted ≥50X in fresh medium and incubated at 30 °C until OD_660_ of 0.3 to 0.5. 100 μl samples were collected in a 96 well plate and kept at -80 °C until measurement. Then, 50 μl of aliquots of the previously frozen samples were thawed and immediately treated with 50 μl Z buffer (60 mM Na_2_HPO_4_, 40 mM NaH_2_PO_4_, 10 mM KCl, 1 mM MgSO_4_, pH 7.0) supplemented with 0.1 g polymyxin B and 0,27 % (v/v) *β*-mercaptoethanol for 30 min at 28 °C. To this, 150 μl of Z buffer were added, followed by 50 μl of 4 mg/ml O-nitrophenyl-β-D-galactopyranoside (ONPG). Then, ONPG hydrolysis was measured at 30 °C for 30 min. The activity of the *β*-galactosidase expressed in miller units (MU) was calculated using the following equation: MU = (OD_420_ × 1,000) / [OD_660_ × t × v] where “t” is the time of the reaction (min), and “v” is the volume of cultures used in the assays (ml). Experimental values were the average of three independent experiments.

### Microscopy

Strains grown in PYE were imaged either in exponential or stationary phase using 1.5 % agar pads with the indicated medium. Cells in osmotic shock conditions were pelleted and washed twice with the indicated stress medium and imaged in 1.5 % agar pads maintaining the stress condition. Images were obtained using Axio Imager Z1 microscope (Zeiss), Orca-Flash 4.0 camera (Hamamatsu) and Zen 2.3 software (Zeiss). Temperature (30 °C) was maintained stable during microscopy analysis using the Tempcontrol 37-analog 1 channel equipment (HemoGenix®) coupled to the Axio Imager Z1 microscope. Images were processed with ImageJ. Demographs were obtained with MicrobeJ by segmenting each cell, integrating fluorescence, sorting cells by length and plotting fluorescence intensity and cellular widths to indicate the relative position of the protein fusions, in which 0 represents mid-cell and 2 or -2 the cell poles (Ducret *et al*., 2016). Confocal microscopy images were obtained using a LSM900 Ayriscan microscope. Samples were prepared for imaging Fresh using cells grown to OD_660_ 0.5. Cells were fixed immediately with a 12.5% paraformaldehyde and 150 mM NaPO_4_ solution. Cells were mixed gently and incubated for 15 min at room temperature, then at 4 °C for 40 min. Then, cells were washed four times with 1.5X volume of dH_2_O. Cells were immobilized in CELLlview slides (Greiner bio-one) by treating wells beforehand with 30 μl 0.1 % poly-L-lysine, then removing poly-L-lysine by aspiration and hydrating wells with a drop water. Thereafter, 10 μl of cells were added to the well and let to sit for 10 mins. Cells in suspension were aspirated and the remain coated cells let to dry for 1 min. Finally, a drop of SlowFade^TM^ Gold Antifade Mountant (Thermo Fisher) was added to wells containing immobilised cells.

### Protein purification and production of polyclonal antibodies

In order to immunize rabbits for production of ChvI polyclonal antibodies His6-ChvI was purified as follows. A BL21(DE3) strain harboring plasmid pET-28a-*chvI* was grown in LB medium supplemented with kanamycin until an OD_600_ of 0.7 was reached. IPTG (isopropyl-*β*-D-thiogalactopyranoside) (Thermo Fisher Scientific) was added at a final concentration of 1 mM, and the culture was incubated at 37 °C for 4 h. Then, cells were harvested by centrifugation for 20 min at 5,000 *g* and 4 °C. The pellet was resuspended in 20 ml BD buffer (20 mM Tris-HCl [pH 8.0], 500 mM NaCl, 10% glycerol, 10 mM MgCl2, 12.5 mM imidazole) supplemented with complete EDTA-free protease cocktail inhibitor (Roche), 400 mg lysozyme (Sigma), and 10 mg DNase I (Roche) and incubated for 30 min on ice. Cells were then lysed by sonication and the lysate cleared by centrifugation (12,000 rpm for 30 min at 4 °C) was loaded on a Ni-nitrilotriacetic acid (Ni-NTA) column and incubated for 1 h at 4 °C with end-over-end shaking. The column was then washed with 5 ml BD buffer, 3 ml Wash1 buffer (BD buffer with 25 mM imidazole), 3 ml Wash2 buffer (BD buffer with 50 mM imidazole), and 3 ml Wash3 buffer (BD buffer with 75 mM imidazole). Proteins bound to the column were eluted with 3 ml elution buffer (BD buffer with 100 mM imidazole) and aliquoted in 300 μl fractions. All the fractions containing the protein of interest (checked by Coomassie blue staining) were pooled and dialyzed in dialysis buffer (50 mM Tris [pH 7.4], 12.5 mM MgCl2). Purified ChvI was used to immunize rabbits (CER Groupe, Marloie, Belgium).

### Immunoblot analysis

Proteins crude extracts were prepared by harvesting cells from exponential growth phase (OD_660_ ∼ 0.3). The pellets were then resuspended in SDS-PAGE loading buffer by normalizing to the OD_660_ before lysing cells by incubating them for 10 min at 95 °C. The equivalent of 0.5 ml of cult (OD_660_ ∼ 0.3) was loaded and proteins were subjected to electrophoresis in a 12% SDS-polyacrylamide gel, transferred onto a nitrocellulose membrane then blocked overnight in 5% (wt/vol) nonfat dry milk in phosphate buffer saline (PBS) with 0.05% Tween 20. Membrane was immunoblotted for ≥ 3 h with primary monoclonal anti-GFP (1:5,000) antibodies (JL8, Clontech-Takara), then followed by immunoblotting for ≤ 1 h with secondary antibodies: 1:5,000 anti-mouse linked to peroxidase (Dako Agilent), and vizualized thanks to Clarity*™* Western ECL substrate chemiluminescence reagent (BioRad) and Amersham Imager 600 (GE Healthcare).

### Chromatin immunoprecipitation followed by deep sequencing (ChIP-Seq) assay

A ChIP-Seq protocol was followed as described by Coppine *et al*. (2020). Briefly, 80 ml of mid-log-phase cells (OD_660_ of 0.6) were cross-linked in 1% formaldehyde and 10 mM sodium phosphate (pH 7.6) at room temperature (RT) for 10 min and then for 30 min on ice. Cross-linking was stopped by addition of 125 mM glycine and incubated for 5 min on ice. Cells were washed twice in phosphate buffer solution (PBS; 137 mM NaCl, 2.7 mM KCl, 10 mM Na_2_HPO_4_, 1.8 mM KH_2_PO_4_, pH 7.4) resuspended in 450 μl TES buffer (10 mM Tris-HCl [pH 7.5], 1 mM EDTA, and 100 mM NaCl), and lysed with 2 μl of Ready-lyse lysozyme solution for 5 min at RT. Protease inhibitors (Roche) were added, and the mixture was incubated for 10 min. Then, 550 μl of ChIP buffer (1.1% Triton X-100, 1.2 mM EDTA, 16.7 mM Tris-HCl [pH 8.1], and 167 mM NaCl, plus protease inhibitors) were added to the lysate and incubated at 37 °C for 10 min before sonication (2 x 8 bursts of 30 sec on ice using a Diagenode Bioruptor) to shear DNA fragments to a length of 300 to 500 bp. Lysate was cleared by centrifugation for 10 min at 12,500 rpm at 4 °C, and protein content was assessed by measuring the OD_280_. Then, 7.5 mg of proteins was diluted in ChIP buffer supplemented with 0.01% SDS and precleared for 1 h at 4 °C with 50 μl of SureBeads Protein A Magnetic Beads (BioRad) and 100 μg bovine serum albumin (BSA). One microliter of polyclonal anti-ChvI antibodies was added to the supernatant before overnight incubation at 4 °C under gentle agitation. Next, 80 μl of BSA presaturated protein A-agarose beads were added to the solution and incubated for 2 h at 4 °C with rotation, washed once with low-salt buffer (0.1% SDS, 1% Triton X-100, 2 mM EDTA, 20 mM Tris-HCl [pH 8.1], 150 mM NaCl), once with high-salt buffer (0.1% SDS, 1% Triton X-100, 2 mM EDTA, 20 mM Tris-HCl [pH 8.1], 500 mM NaCl), once with LiCl buffer (0.25 M LiCl, 1% NP-40, 1% deoxycholate, 1 mM EDTA, 10 mM Tris-HCl [pH 8.1]), and once with TE buffer (10 mM Tris-HCl [pH 8.1] 1 mM EDTA) at 4 °C, followed by and a second wash with TE buffer at RT. The DNA-protein complexes were eluted twice in 250 μl freshly prepared elution buffer (0.1 M NaHCO3, 1% SDS). NaCl was added at a concentration of 300 mM to the combined eluates (500 μl) before overnight incubation at 65 °C to reverse the cross-link. The samples were treated with 20 μg of proteinase K in 40mM EDTA and 40 mM Tris-HCl (pH 6.5) for 2 h at 45 °C. DNA was extracted using a Nucleospin PCR cleanup kit (Macherey-Nagel) and resuspended in 50 μl elution buffer (5 mM Tris-HCl [pH 8.5]). DNA sequencing was performed using an Illumina NextSeq 550 (paired-end 2x75) instrument (BIO.be). NGS data were analysed as described in Coppine et al., 2020.

### RNA-seq

WT and *ΔchvI* cells were grown overnight to OD_660_ ∼ 0.3 and then exposed to 6% sucrose for 4 h. Thereafter, total RNA was extracted with RNeasy® Protect Bacteria Kit from Qiagen and following manufacturer’s instructions. The quantity and quality (A260/A280 ration) of RNA was determined with a Thermo Scientific ™ Nanodrop ™ One Microvolume UV-Vis Spectrophotometer. RNASeq TTRNA libraries were prepared according to the manufacturer’s instructions and sequenced with Illumina NovaSeq 6000 (paired-end 2x100) instrument (BIO.be). NGS data were analysed using Galaxy (https://usegalaxy.org) (Afgan *et al.,* 2018). Briefly, FastQC was used to evaluate the quality of the reads; HISAT2 was used to map the reads onto the NA1000 reference genome (NC_011916.1) and generate bam files; featureCounts was used to generate counts tables using bam files and DESeq2 was used to determine differentially expressed genes. The Volcano plot was generated using GraphPad Prism 9 software.

### *In vivo* ^32^P labelling

Cells were grown overnight in PYE, then washed twice and grown in M5G medium lacking phosphate and was grown overnight in M5G with 0.05 mM phosphate to OD_660_ ∼ 0.3. Then, one ml of culture was labelled for 4 min at 30 °C using 30 μCi *γ*-[^32^P]ATP (PerkinElmer). Samples were incubated with 6% sucrose, 10 μg/ml vancomycin or 100 μg/ml vancomycin for 7 minutes at 30 °C. Then, cells were pelleted for 2 min at 15,000 rpm and the supernatant removed completely without disturbing the pellets. Cell pellets were resuspended in 50 μl Lysis Buffer (50 mM Tris pH 7.0, 150 mM NaCl, 80 mM EDTA, 2% Triton X100; sterilised with 2 μm Acrodisc syringe filters) by pipetting. Lysed samples were incubated 3 min on ice, 450 μl of cold LBS were added and proteins were collected using centrifugation for 15 min at 13,000 and 4 °C. Then, 40 μl of SureBeads Protein A Magnetic Beads (BioRad) were washed 4 times in 1 ml PBS + 0.1% Tween (PBS-T) and finally resuspended in 65 μl. Thereafter, 3 μl of anti-ChvI antibody were added to the washed protein A-agarose beads and incubated at RT in a shaker at 1300 rpm. Protein A beads with anti-ChvI antibodies were washed 3 times with PBS-T and 1 time with cold (4 °C) Low Salt Buffer (LSB; 50 mM Tris pH 7.0, 100 mM NaCl, 50 mM EDTA, 2% Triton X100; sterilised with 2 μm Acrodisc syringe filters), immediately resuspended in 50 μl LSB and kept in ice. ChvI immunoprecipitation (IP) was performed by adding the Protein A beads with anti-ChvI antibodies previously prepared to the cell-free protein samples, and incubating at 4 °C with rotation for 90 minutes. IP samples were washed 1 time in 1 ml LSB and 3 times in 1 ml High Salt Buffer (50 mM Tris pH 7.0, 500 mM NaCl, 50 mM EDTA, 0.1% Triton X100; sterilised with 2 μμ Acrodisc syringe filters). Samples were eluted using 25 μl of 2.5X SDS-loading Buffer and incubation at 37 °C for 5 min. Samples with SDS loading Buffer were separated from magnetic beads, and 20 μl loaded on a SDS-Page gel and run at 200 V-50 mA-100 W. The gel was dried for 1h at 70 °C under vacuum in a Model 583 Gel Dryer (BioRad). Finally, the dried gel was exposed on phosphoscreen for 5 days and revealed using a Cyclone Plus Phosphor Imager (PerkinElmer).

### Statistical analyses

All the statistical analyses were performed using GraphPad Prism 9 software. A *P* value of <0.05 was considered as significant.

### Data availability

ChIP-Seq and RNA-Seq data have been deposited to the Gene Expression Omnibus (GEO) repositery with the accession number GSE200466.

## Results

### High osmolyte concentration in M2G impairs growth of *chvIG* mutants

It was shown previously that *C. crescentus chvIG* mutants failed to grow in minimal media with xylose as sole carbon source (M2X), while these mutants grew similarly to WT cells in complex media (PYE) (Stein *et al.,* 2021). Likewise, we observed that mutants inoculated in M2G (with glucose as sole carbon source) in both solid and liquid media were impaired for growth (**Fig. 1A-B**). To date, the cause for growth impairment in *chvIG* mutants in minimal media has not been determined. Considering that stressful conditions are present in synthetic minimal media (M2G or M2X) but missing in PYE, we tested osmolytes as causative agents. Indeed, the synthetic minimal media M2G contains a higher concentration of osmolytes than the complex media PYE (Hocking *et al.,* 2012). Therefore, we assessed the viability of *chvIG* mutants on plates lacking Na^+^ and K^+^ salts (**Fig. 1A**, M2G 0% Na^+^, K^+^). In these conditions the *chvI* mutant grew similarly to WT cells. We also reduced these osmolytes in liquid cultures, and observed that growth was restored in the *chvI* mutant when half (and below) of the regular concentration found in M2G was used (**Fig. 1B**, M2G 50% Na^+^, K^+^). Furthermore, we observed that adding osmolytes (NaCl, sucrose and those in M2 salts) to complex media impaired growth and decreased viability of a *chvI* mutant (**Fig. 1A**, PYEX + 40 mM NaCl, PYE + 6% Sucrose; **Fig. 1C**, PYE + M2 salts). Additionally, ectopic expression of *chvI* from the xylose-inducible promoter (P*_xylX_::chvI*) in a *chvI* mutant restored growth in PYEX plates supplemented with NaCl whereas expressing back *chvI* in a *chvIG* mutant did not (**Fig. 1A**). Finally, while a phospho-mimetic mutant of *chvI* (*chvI_D52E_*) grew similarly to WT, a phospho-ablative mutant of *chvI* (*chvI_D52A_*) failed to propagate in M2G (**Fig. 1D**). Together, our data show that a fully functional ChvGI TCS is required to survive and grow in hyperosmotic environments.

**Fig. 1.**
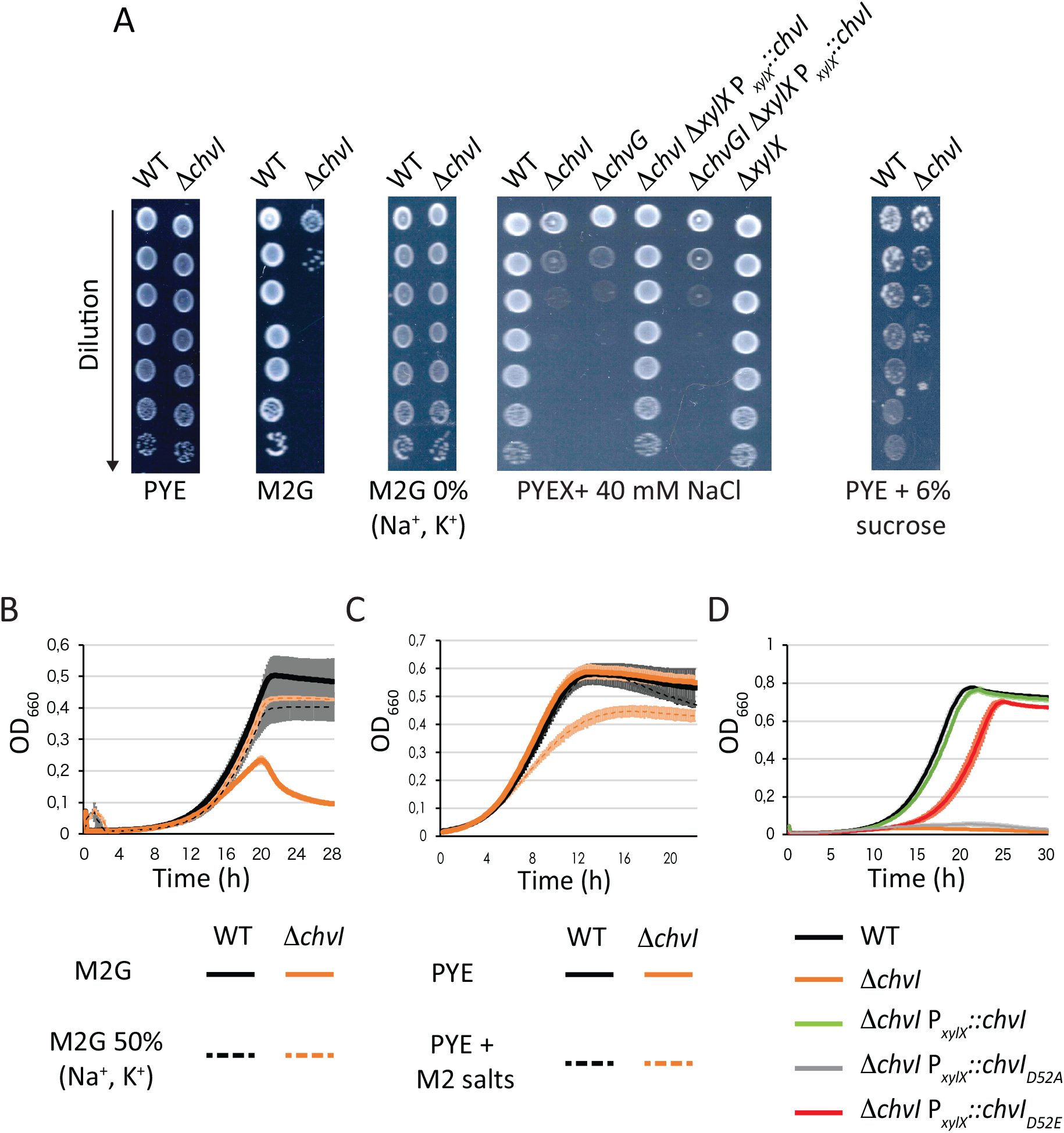
Viability and growth of *chvGI* mutants in hyperosmotic conditions. (A) Viability of WT and *ΔchvI* cells in complex (PYE) and synthetic minimal media (M2G) plates with varying osmolytes concentrations. For complementation assays, PYE was supplemented with 0.1% xylose (PYEX). (B) Growth in minimal media with 100% (solid lines) or 50% (dashed lines) Na_2_HPO_4_ and KH_2_PO_4_ concentrations referred to M2G. (C) Growth in PYE supplemented with 1X M2 salts. (D) Growth in M2G of mutants complemented with WT, phospho-ablative (*chvI_D52A_*) or phospho-mimetic (*chvI_D52E_*) mutants of *chvI* expressed from the xylose-inducible promoter P*_xylX_*. Data from (B) to (D) represent the average value of biological replicates (n=3, error bars show standard deviation). WT and *ΔchvI* mutant are represented in (B) to (D) in black and orange, respectively.

### Inactivating *chvT* or *ntrX* improves fitness of a *chvI* mutant under osmotic stress

Considering that mutations in *chvT* and *ntrX* partially alleviated the viability impairment of *chvI* mutants in minimal media (Stein *et al*., 2021), we tested whether *chvT* or *ntrX* inactivation could also partially protect *chvI* mutants from hypertonic conditions. We first confirmed that a *ΔchvI* mutant was impaired for growth under osmotic stress by using an excess of KCl in PYE (**Fig. 2**, PYEX + 50 mM KCl). In contrast, neither *ntrX* nor *chvT* inactivation did interfere with viablity in PYE + KCl conditions compared to WT (**Fig. 2**). The *ntrX* gene (previously annotated as CC1743) was suspected to be essential for growth in complex PYE medium but dispensable for growth in synthetic minimal medium (Skerker *et al.,* 2005). In agreement with that report, we found that the viability of *ntrX* mutants was reduced in PYE (**Fig. 2**). Notwithstanding this fitness cost, *ΔntrX ΔchvI* double mutants could be generated and grown on PYE, as shown independently in this study and by Stein *et al*. (2021). More importantly, we observed that inactivating *ntrX* or *chvT* in a *ΔchvI* background partially restored growth in hypertonic conditions. These data indicate that additional cell envelope defects in *ΔchvI* mutants are responsible for growth impairment on hypertonic conditions. Altogether, these data suggest that the homeostasis of the cell envelope is lost in *chvIG* mutants, leading to a hypersensitivity to osmolytes.

**Fig. 2.**
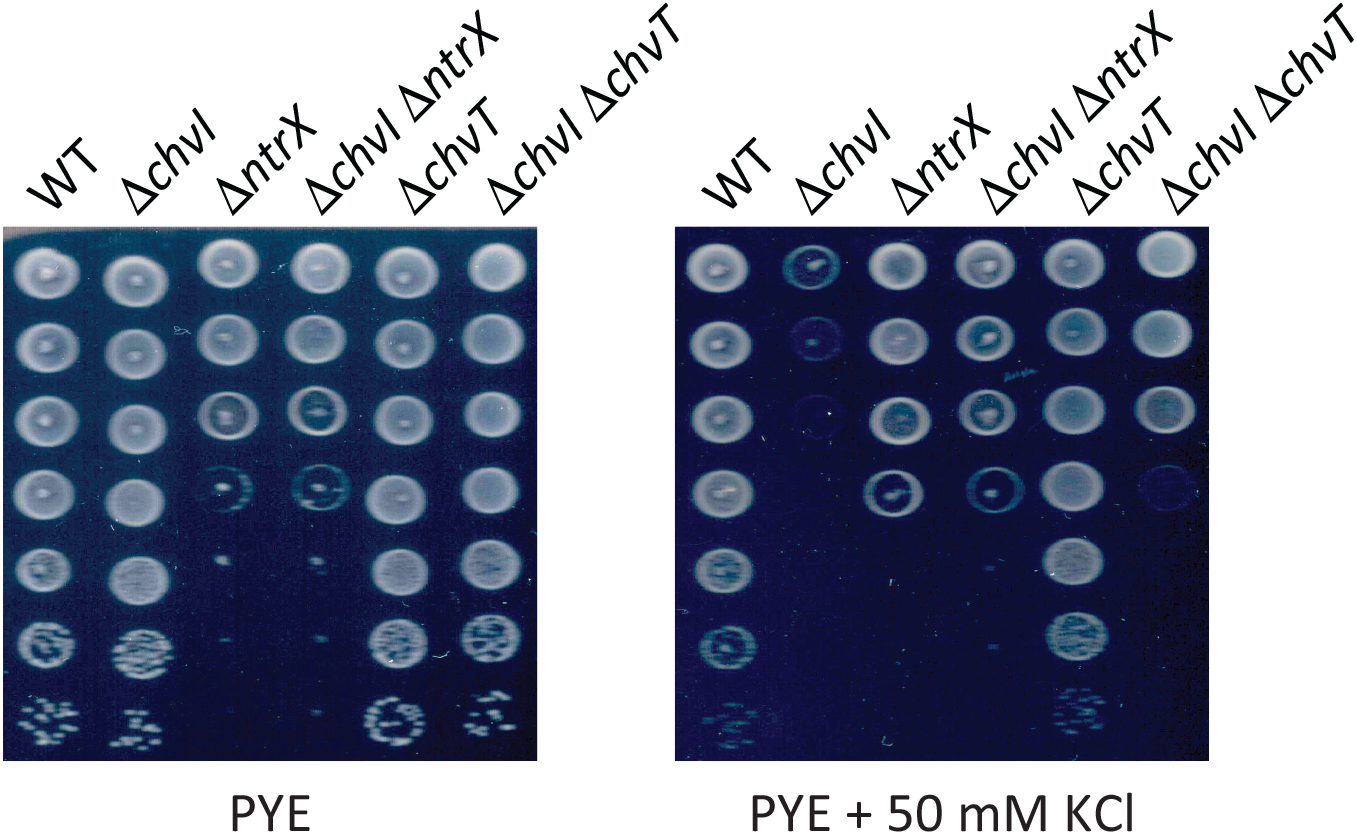
Growth and viability of *chvI*, *chvT* and *ntrX* mutants in hypertonic conditions. Viability of single (*ΔchvI*, *ΔchvT* and *ΔntrX*) and double (*ΔchvI ΔchvT* and *ΔchvI ΔntrX*) mutants on PYE agar with or without 50 mM KCl. Images represent three biological replicates.

### The ChvI regulon reveals osmotic stress and cell envelope-related target genes

Transcriptomic analyses were previously performed on a *ΔchvI* mutant overexpressing the phospho-mimetic mutant *chvI_D52E_* in PYE (Stein *et al.,* 2021). Nonetheless, cells expressing *chvI_D52E_* as the only copy, either from its own promoter at the endogenous *chvI* locus (P*_chvI_::chvI_D52E_*) or ectopically from the xylose-inducible promoter (P*_xylX_::chvI_D52E_*) led to a slight growth delay and cell filamentation (**Fig. 1D** and **Fig. S1**). This suggests either that ChvI_D52E_ does not perfectly mimic phosphorylated ChvI or that unphosphorylated ChvI is also required for optimal growth in synthetic minimal media. Hence, we determined the ChvI regulon of WT cells grown in M2G by using ChIP-seq with polyclonal antibodies targeting ChvI, which allowed to unveil the DNA regions directly bound by ChvI (**Fig. 3A**, **Table S4**).

**Fig. 3.**
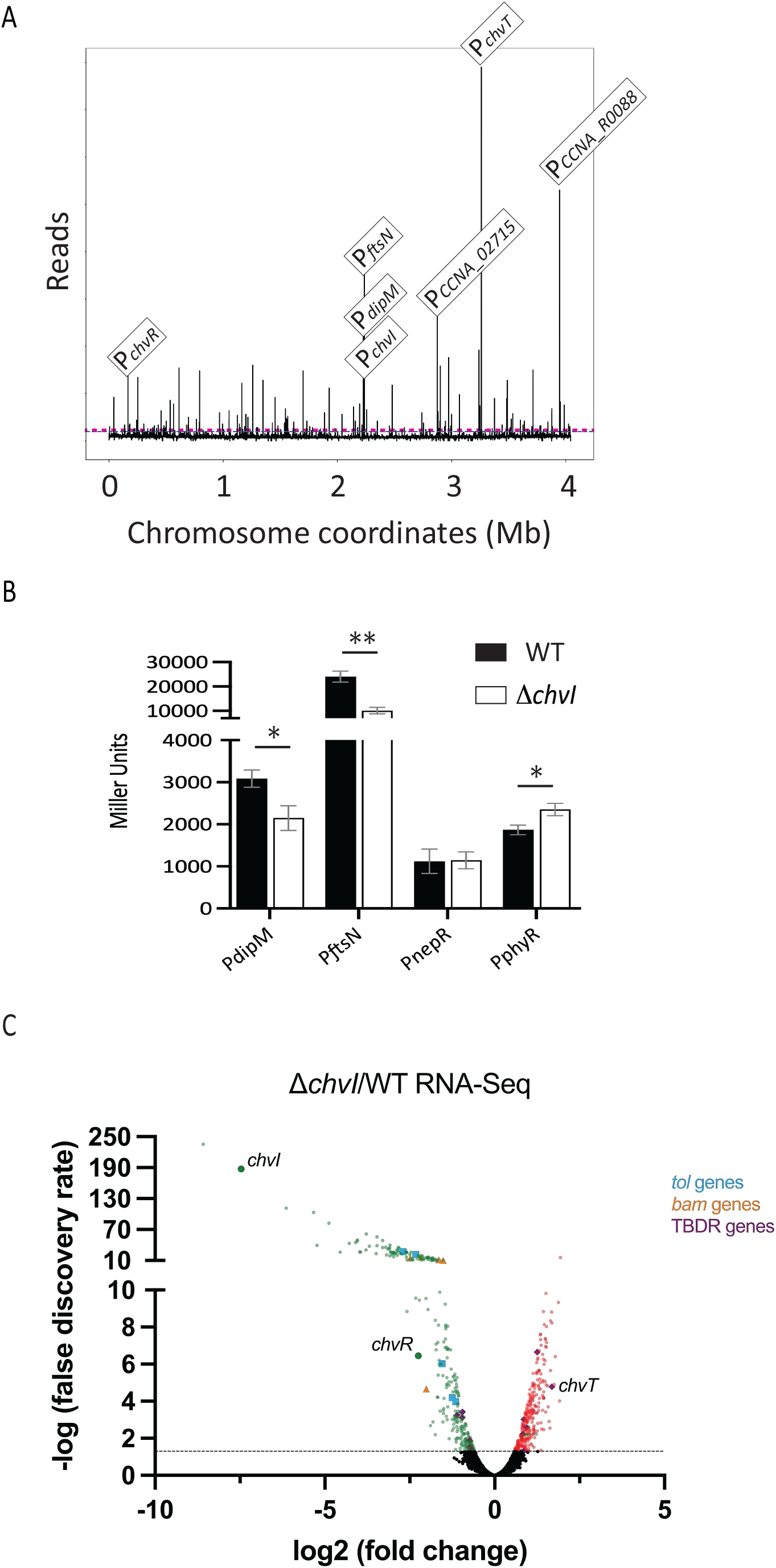
ChvI target genes determined by ChIP-seq and validated by β-galactosidase assays. (A) Genome-wide occupancy of ChvI on the chromosome of *C. crescentus* determined by ChIP-seq on WT strain grown in M2G. The x-axis represents the coordinates on the genome (Mb), the y-axis shows the normalized ChIP-Seq read abundance in reads. Some top hits, corresponding to the promoter regions of *chvT*, *CCNA_R0088*, *ftsN, CCNA_02715*, *chvR*, *dipM* and *chvI* operon, are highlighted. (B) Activity of the *dipM*, *ftsN*, *nepR* and *phyR* promoters (Miller Units) in WT (black bars) and *ΔchvI* (white bars) cells grown O.N in PYE, then washed and exposed for 4h in M2G. The data represent the average values of biological replicates (n=3, error bars show standard deviation). * = *p* < 0.05, ** = *p* < 0.01, single factor ANOVA analysis of *β*-galactosidase activity. C. Volcano plot representing the relation between the fold change and P values on gene expression between *ΔchvI* and WT strains exposed to osmotic upshift in PYE 6% sucrose by RNA-seq. Genes identified are presented as dots. Significant down- and up-regulated genes are presented as green and red dots, respectively, while genes with no significant alterations are presented as black dots. *chvI, chvR* and *chvT* genes as well as genes encoding Tol (blue) and Bam (orange) complexes or TonB-dependent receptors (purple) are highlighted.

From our experiment, 190 peaks (DNA binding sites) were identified for ChvI (**Fig. 3A**, **Table S4**). Interestingly, the top target is the promoter region of *chvT* itself, suggesting that ChvI regulates ChvT not only (i) indirectly at the post-transcriptional level via transcriptional activation of the sRNA *chvR* but also (ii) directly at the transcriptional level and (**Fig. 3A**). Besides *chvT* and *chvR*, we identified potential new targeted genes involved in (i) cell division, morphology and peptidoglycan (PG) synthesis, such as *mreB*, *ftsZ*, *ftsN* and *dipM*; and (ii) general stress response (GSR), such as *sigT* (sigma factor T), *nepR* (*sigT* antagonist) and *phyKR* (TCS regulating NepR activity negatively).

To further confirm some of the new targeted genes, we tested their expression by measuring their promoter activity upon osmotic shock. As shown in **Fig. 3B**, the activity of *dipM* and *ftsN* promoters was significantly lower in the *ΔchvI* mutant compared to WT, whereas *phyR* promoter activity was significantly higher in *ΔchvI* than in WT. In contrast, the *nepR* promoter had similar activity in both WT and *ΔchvI* strains. These results suggest that ChvI can work as an activator (e.g. *dipM* and *ftsN*) or a repressor (e.g. *phyR*) of gene expression.

Interestingly, *hprK* is part of the SigT regulon determined upon osmotic shock (Tien *et al*., 2018). Given that *hprK* is part of the same operon as *chvIG*, we tested whether the entire operon *chvIG-hprK* could be under the control of SigT. Consistently, we found that in cells grown in PYE, washed and exposed for four hours in M2G, the *chvI* promoter (P*_chvI_*) was significantly higher in the *ΔsigT* mutant compared to WT whereas the activity of P*_chvI_* was strongly reduced in the *ΔchvI* mutant (**Fig. S2A**). Thus, SigT is a negative regulator of ChvI while ChvI is subjected to a positive feedback loop. Considering that *phyR* promoter activity was higher in a *ΔchvI* background (**Fig. 3B**), this indicates that ChvI regulates negatively SigT by inhibiting PhyR. To better understand the regulatory mechanism between ChvI and SigT we tested viability of mutants for the genes in osmotic conditions used in this study, that is PYE supplemented 40 mM NaCl (**Fig. 1A**), but also in more stringent conditions used in a previous study, that is PYE supplemented 85 mM NaCl (Alvarez-Martinez *et al.,* 2007) (**Fig. S2B**). We observed that, contrary to *chvI*, *sigT* is not essential for viability in PYE + 40 mM NaCl. In contrast, at a higher osmotic regime (PYE + 85m M NaCl) the viability of *ΔsigT* declined. Moreover, inactivating *sigT* in a *ΔchvI* background did not aggravate the poor viability of *ΔchvI* mutant upon osmotic stress. Together, these data suggest that ChvGI responds to lower osmotic stress than SigT and that both SigT and ChvI antagonise each other at the transcriptional level.

We also performed RNA-seq on WT and *ΔchvI* cells grown in PYE and exposed to 6% sucrose for 4 hours. We found 301 down-regulated and 293 up-regulated genes with ≥1.5-fold change and a false-discovery rate [FDR] P*adj* ≤ 0.05 (**Fig. 3C**; **Tables S5** and **S6**). Among these genes, 126 down-regulated and 44 up-regulated genes were identified in our ChIP-seq experiment (**Tables S5** and **S6**). This further suggests that ChvI performs direct dual positive and negative transcriptional regulation, but mostly positive. Furthermore, we confirmed that upon osmotic shock, ChvI regulates the expression of multiple genes involved in cell envelope homeostasis such as for example, *tol* genes, *bam* genes and TBDR-encoding genes (**Fig. 3C**; **Tables S5** and **S6**).

### ChvI is phosphorylated under osmotic upshift

Since both single *ΔchvG* and *ΔchvI* mutants are sensitive to osmotic upshift (**Fig. 1A**), this suggests that ChvI might be phosphorylated and activated by ChvG in such stressful conditions. We tested this hypothesis by determining the *in vivo* phosphorylated levels of ChvI (ChvI∼P) with or without osmotic shock. In order to facilitate uptake of [γ-^32^P] ATP, *Caulobacter* cells were grown in a medium depleted for the phosphate salts Na_2_HPO_4_ and KH_2_HPO_4_ (M5GG). Hence, K^+^ and Na^+^ concentrations in M5GG are respectively 1.01 mM and 1.02 mM, which is much less that in M2G (7.75 mM of K^+^ and 12.25 mM of Na^+^). At such low concentrations of osmolytes, the *ΔchvI* strain grew, as expected, similarly to WT in M5GG (**Fig. 4A**). In contrast and in agreement with our previous findings, the growth of the *ΔchvI* mutant was severely impaired in M5GG supplemented with 6% sucrose compared to WT (**Fig. 4A**). Consistent with these data, we observed that ChvI was barely phosphorylated in WT grown in M5GG while no ChvI∼P was detected in *ΔchvG* (**Fig. 4B**). More importantly, ChvI was hyperphosphorylated in cells grown under osmotic shock with sucrose (**Fig. 4C-D**).

**Fig. 4.**
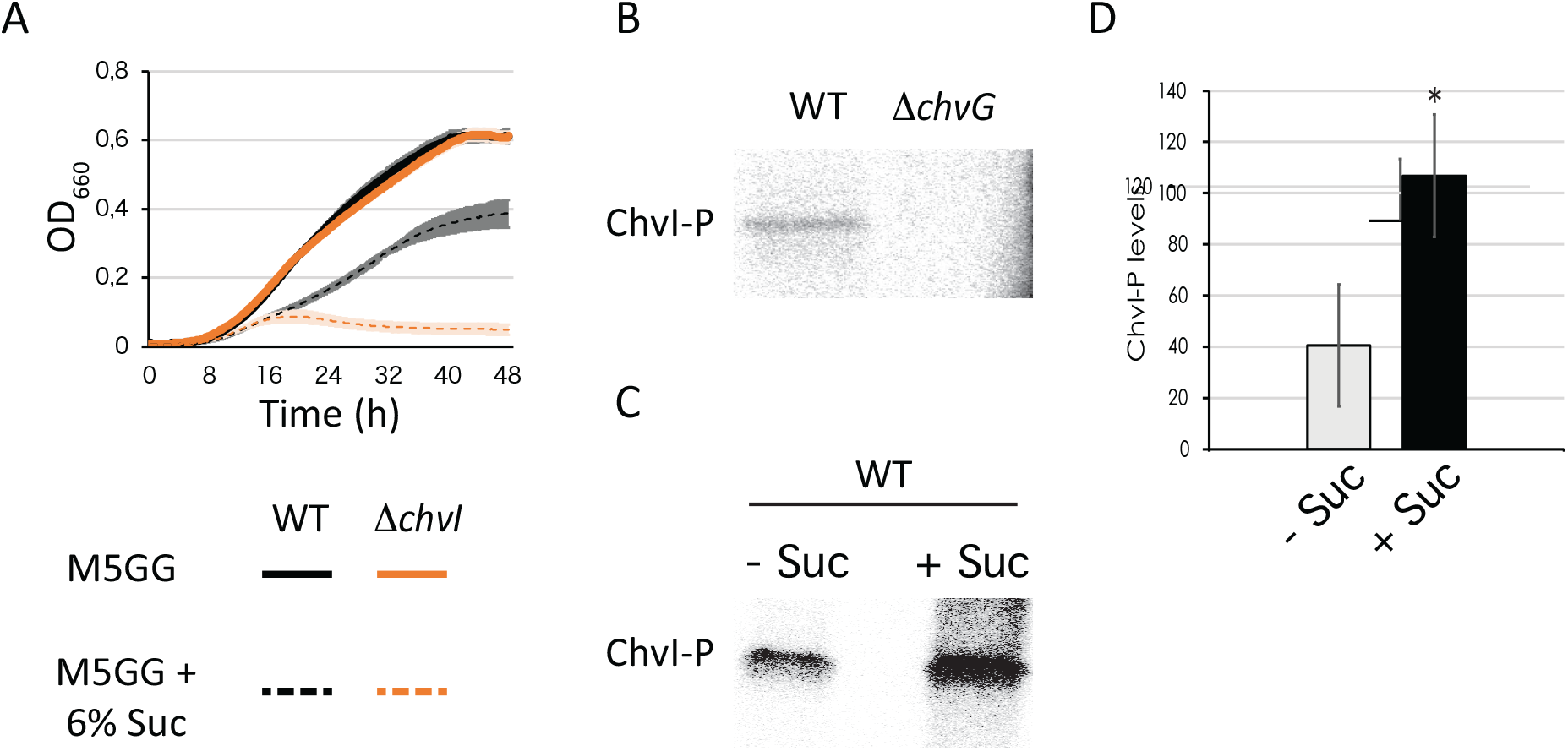
ChvG-dependent phosphorylation of ChvI is stimulated upon osmotic shock. (A) Growth of WT (black) and *ΔchvI* mutant (orange) in M5GG with (dashed lines) or without (solid lines) 6% sucrose (Suc). The data represent the average value of biological replicates (n=3, error bars show standard deviation). (B) *In vivo* phosphorylation levels of ChvI in WT and *Δchv*G mutant grown in M5GG. (C) *in vivo* phosphorylation levels of ChvI in WT grown in M5GG without (-) or with (+) sucrose (Suc) for 7 min. (D) Quantified *in vivo* phosphorylation levels of ChvI in the same growth conditions than in (C). The data represent the average values of biological replicates (n=3, error bars show standard deviation). * = *p* < 0.05, single factor ANOVA analysis of ChvI phosphorylation.

### ChvGI is sensitive to peptidoglycan synthesis inhibition

Stress response systems surveying cell envelope homeostasis in Gram-negative bacteria are sensitive to different stressful conditions including osmotic shock (Jubelin *et al.,* 2005; Laubacher & Ades, 2008; Cho *et al.,* 2014). Moreover, deletion of the *chvGI-hprK* operon have been shown to result in sensitivity to vancomycin (Vallet *et al.,* 2020), which targets the D-Ala-D-Ala moiety of the PG precursors thereby inhibiting PG crosslinking. First, we confirmed that vancomycin impairs growth and viability of the single *ΔchvI* mutant (**Fig 5.A** and **E**). We also tested growth and viability upon treatment with other cell envelope stressors, such as acidic pH, detergent and antibiotics targeting the outer membrane or the PG synthesis machinery. We did not observe significant impacts on fitness between WT and *ΔchvI* cells under treatment with acidic stress, A22 drug targeting the actin-like protein MreB, SDS and polymyxin B which target the outer membrane, and the antibiotic cephalexin, which inhibits the PG transpeptidase activity of the Penicillin Binding Protein 3 (Pbp3) associated to the cell division machinery (divisome) (**Fig. S3A-B**). In contrast, we found that exposure to other antibiotics targeting PBPs associated to cell elongation machinery (elongasome) impaired growth of the *ΔchvI* mutant. For instance, treatment with mecillinam, which inhibits PG crosslinking during elongation by binding to Pbp2 significantly impaired growth and viability of *ΔchvI* cells compared to WT (**Fig. 5B** and **E**). Also, treatment with antibiotics targeting the bifunctional glycosyltranferase/transpeptidase Pbp1a, such as Cefsulodin and Moenomycin also diminished the growth and viability of the *ΔchvI* mutant (**Fig. 5B-C** and **E**). Altogether, these results indicated that ChvGI is a TCS responding to osmotic upshift but also to PG crosslinking damage.

**Fig. 5.**
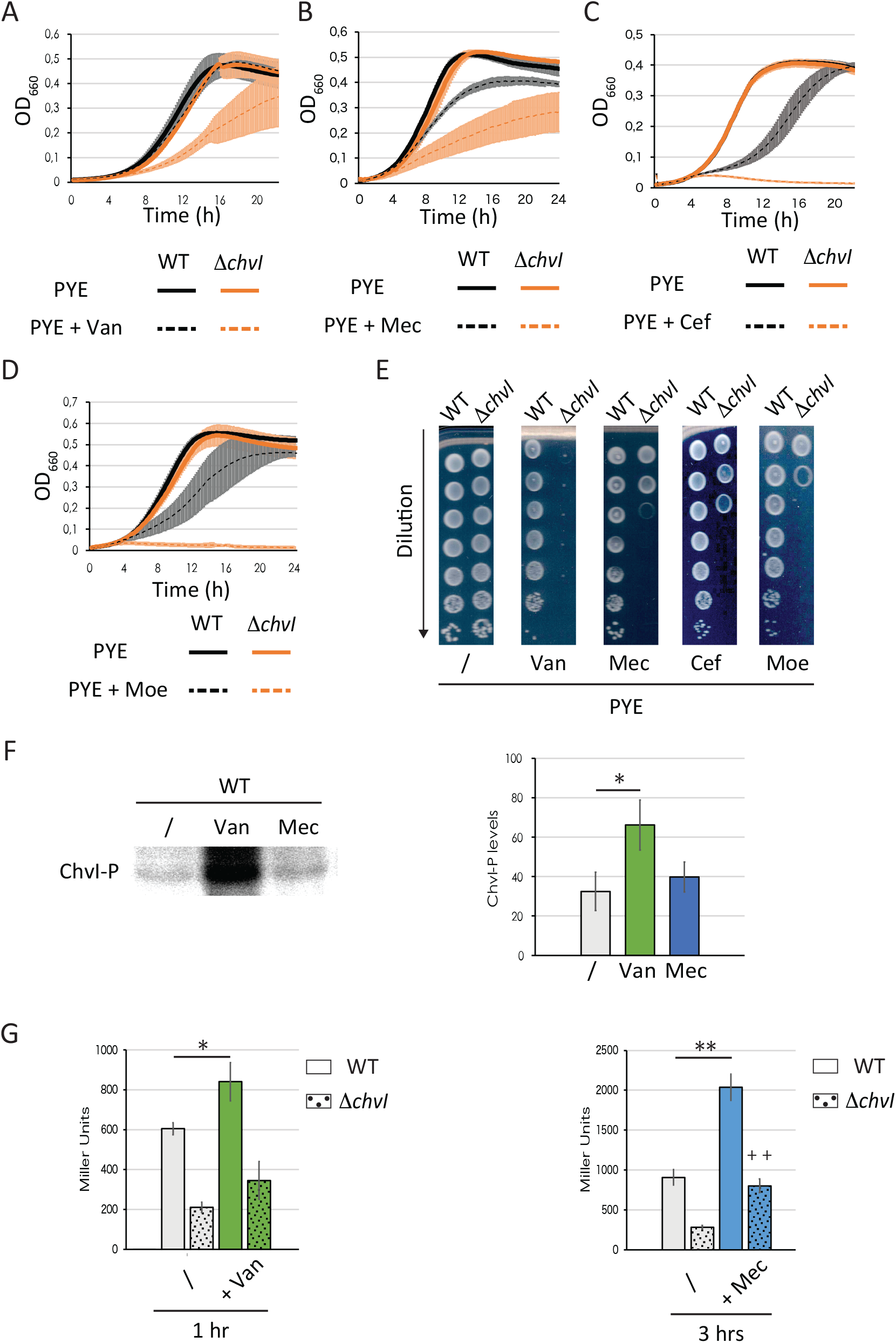
ChvI responds to treatment with antibiotics targeting transpeptidation of peptidoglycan during elongation. (A-D) Growth of WT and *ΔchvI* mutant cells in PYE supplemented with either Vancomycin (Van) (A), mecillinam (Mec) (B), cefsulodin (Cef) (C) or moenomycin (Moe) (D). (E) Viability of WT and *ΔchvI* mutant upon treatment with the antibiotics indicated above. Images are representative of three biological replicates. (F) Quantified *in vivo* phosphorylation levels of ChvI in WT grown in M5GG without antibiotics (/) or with 10 μg/ml vancomycin (Van, green) or 100 μg/ml mecillinam (Mec, blue) for 7 min. (G) Activity of the *chvI* promoter P*_chvI_* (in Miller Units) in WT (solid bars) and *ΔchvI* (dashed bars) cells grown in minimal media M5GG supplemented with antibiotics at the indicated times. The data in figures (A-D) and (F-G) represent the average values of biological replicates (n=3, error bars show standard deviation). * = *p* < 0.05 and ** = *p* < 0.01, and ++ = *p* < 0.01 for comparison with *ΔchvI* cells using single factor ANOVA analysis. WT and *ΔchvI* mutant are represented in (A) to (D) in black and orange, respectively.

To confirm this, similar to cell exposed with osmotic stress, we assessed the phosphorylation of ChvI following exposure to vancomycin or mecillinam antibiotics (**Fig. 5F**). We observed that ChvI phosphorylation increased after vancomycin but not mecillinam treatment. To better understand this result, and considering that the *chvIG-hprK* operon is under positive autoregulation by ChvI∼P, we assessed the activity of the P*_chvI_::lacZ* transcriptional reporter in synthetic media M5GG supplemented with vancomycin or mecillinam (**Fig. 5G**). We observed that activity of the *chvI* promoter significantly increased after exposure to both antibiotics but the response was faster with vancomycin (1 hr) than with mecillinam (3 hrs) treatment. Surprisingly, although the activity of P*_chvI_::lacZ* was lower in *ΔchvI* cells than in WT cells, it remained higher in *ΔchvI* treated cells than in *ΔchvI* untreated cells, even though this difference was significant only for mecillinam (**Fig. 5G**). Together, these data suggest that (i) the response to mecillinam treatment is slower than the one to vancomycin and that (ii) *chvIG-hprK* expression following exposure to antibiotics is induced not only by ChvI itself but also in a ChvI-independent way.

### ChvG relocates from patchy-spotty to midcell upon osmotic shock

Intriguingly, the HK ChvG was previously found in an automated large-scale analysis for protein localisation as displaying a patchy-spotty distribution typically observed with PG-related proteins (Werner *et al.,* 2009). Therefore, we constructed fluorescent protein fusions to analyse ChvG and ChvI localisation patterns on PYE and M2G pads. First, we confirmed that GFP fusions to the N- or C-terminal extremity of ChvG or ChvI were stable and functional, as attested by western blot analyses and growth assays (**Fig. S4AB**). Then, fluorescent microscopy images showed that both ChvG N- and C-terminal fusions to GFP displayed patchy-spotty localisation patterns in cells grown in PYE or M2G cultures (**Fig. 6A**, **Fig. S4C**). We also confirmed the patchy-spotty localisation pattern for the ChvG C-terminal fusion to GFP in high resolution confocal microscopy (**Fig. S4D**). Surprisingly, when cells grown in PYE were washed in M2G and imaged on M2G agar pads, we observed some cells formed foci near mid-cell (**Fig. 6A**). In contrast, cells grown in M2G, washed with PYE and imaged on PYE pad kept their patchy-spotty localisation pattern (**Fig. 6A**). This relocation is reminiscent to what was described for the PG-related proteins RodA, PBP2 and PBP1A, whose localization changed from a typical patchy-spotty distribution to mid-cell when cells were shifted from PYE to M2G agarose pads (Hocking *et al.,* 2012).

**Fig. 6.**
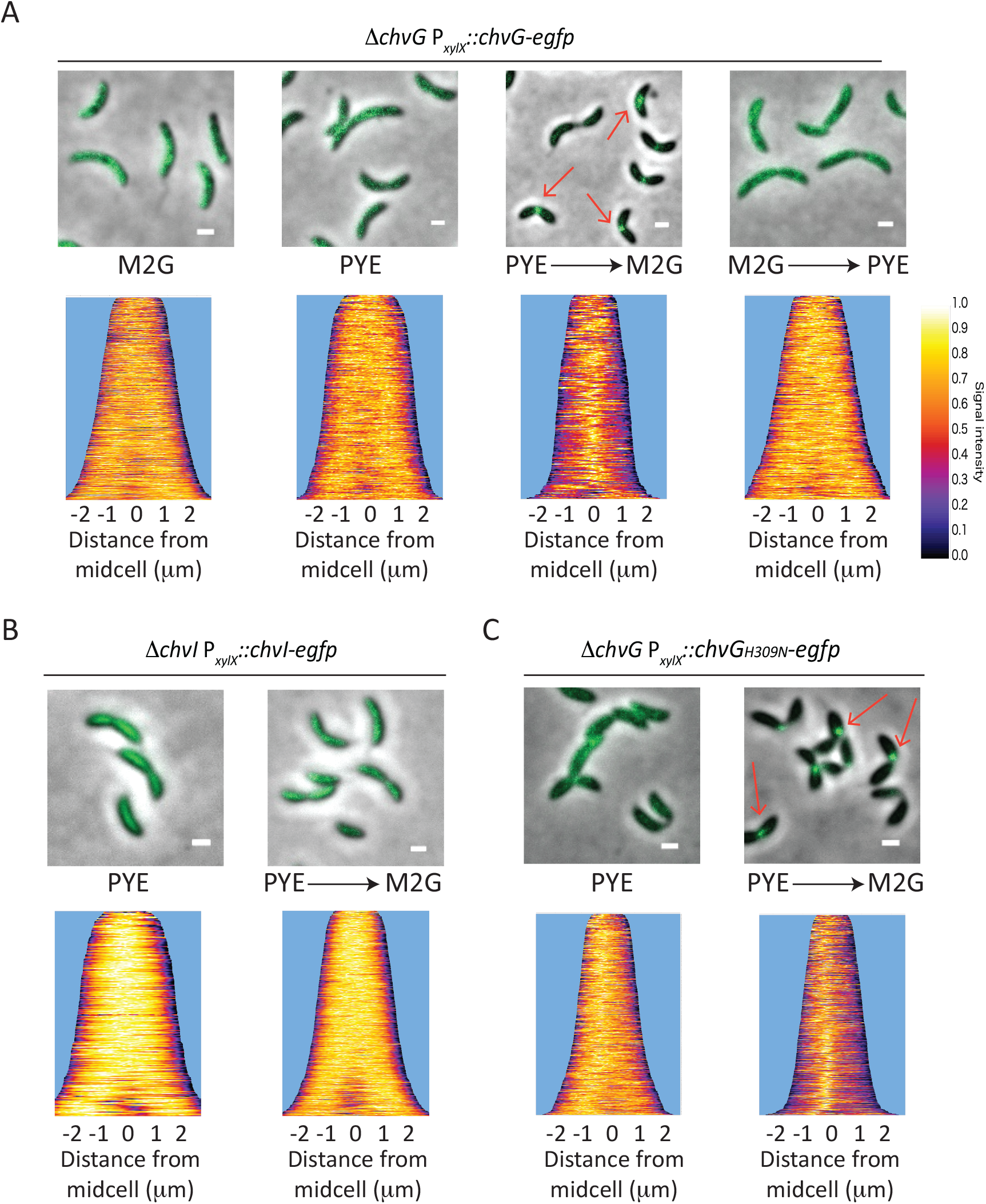
ChvG relocates from a patchy-spotty pattern to mid-cell upon osmotic upshift. (A) Localisation of ChvG-eGFP in cells grown in either M2G or PYE and imaged on M2G or PYE agar pads, respectively; or cells grown in PYE, washed in M2G and imaged on M2G agarose pads (PYE → M2G); or cells grown in M2G, washed in PYE and imaged on PYE agarose pads (M2G → PYE). (B) Localisation of ChvI-eGFP in a *ΔchvI* background and (C) ChvG_H309N_-eGFP in a *ΔchvG* background grown overnight in PYE and imaged on PYE agarose pads or grown in PYE, washed in M2G and imaged on M2G agarose pads (PYE → M2G). Demograph data represent cells sorted from short to long length with no less than 200 cells per sample. Liquid cultures and pads were supplemented with 0.1 % xylose to allow expression of ChvG, ChvI and ChvG_H309N_ fused to eGFP fusions. Scale bar= 1 μm.

In contrast to ChvG, none of the ChvI fusions to GFP (ChvI-GFP and GFP-ChvI) displayed patchy-spotty localization nor foci formation was detected after transition from PYE to M2G pads (**Fig. 6B** and data not shown). Instead, the signal was diffused all over the cell body, indicating that the protein remains diffused in the cytoplasm. Then, we wanted to assess if the catalytic activity had any impact on the protein localisation. For that, we fused GFP to a full-length catalytic mutant of ChvG to (ChvG_H309N_), which like *ΔchvG* was unable to grow on M2G (**Fig. S4E-F**). We observed that, similar to the WT ChvG-GFP, the ChvG_H309N_-GFP showed a patchy-spotty pattern when cells were grown in PYE and relocated at mid-cell when shifted from PYE to M2G (**Fig. 6C**). Thus, our data suggest that ChvG co-localizes with PG synthesis machinery independently of its kinase activity.

### The N-terminal extremity of ChvG determines localisation and relocation

ChvG is a HK anchored in the membrane thanks to 2 transmembrane helices which delimit a periplasmic sensor domain (amino acids 50-221), with a cytoplasmic signal transduction histidine kinase domain (amino acids 242-534) (**Fig. 7A**). To assess which of these ChvG domains are essential for localisation, we fused to mCherry ChvG versions harbouring either the complete (ChvG_1-274_ and ChvG_1-202_) or truncated sensor (ChvG_1-114_) or the catalytic (ChvG_273-534_) domain. First, we showed that both sensor and catalytic domains are essential for *C. crescentus* to propagate in high osmolytes conditions like in M2G since mutants harbouring in frame deletion of each of these domains (ChvG_274-534_ or ChvG_1-274_) were unable to survive in M2G (**Fig. S4G**). Then, we observed that full-length ChvG fused to mCherry displayed similar localisation and relocation patterns (**Fig. 7B-C**) than the ones described for GFP fusions (**Fig. 6A**). Interestingly, the truncated or complete sensor domain alone (ChvG_1-114_, ChvG_1-202_, ChvG_1-274_) fused to mCherry were both localized as patchy-spotty in PYE (**Fig. 7B**). However, only the full-length ChvG-mCherry and ChvG_1-274_-mCherry fusions relocated as foci upon transition to M2G, even though ChvG_1-274_-mCherry foci were not often found at midcell (**Fig. 7C**). Unlike the sensor domain truncates, the catalytic domain (ChvG_273-534_) fused to mCherry displayed neither the patchy-spotty nor the foci pattern of localization and was instead diffusely localized in the cytoplasm (**Fig. 7**B-C). Altogether, our results indicate that the complete periplasmic sensor domain of ChvG edged by transmembrane domains are required for relocation as foci upon osmotic shock whereas the first 114 amino acids are sufficient to determine the patchy-spotty pattern of localization.

**Fig. 7.**
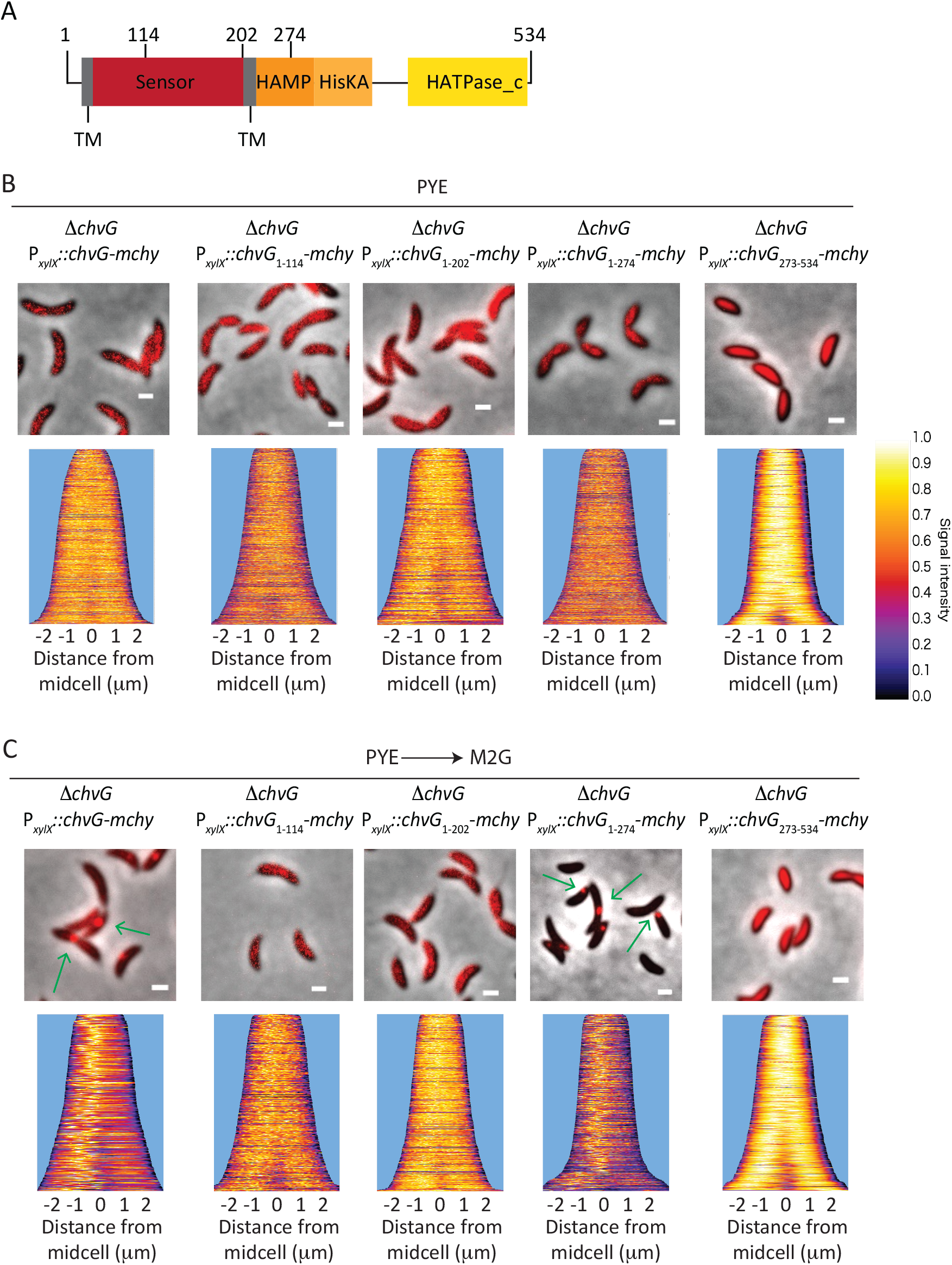
The N-terminal periplasmic sensor domain of ChvG is required for patchy-spotty localisation pattern and mid-cell relocation. (A) Conserved domains in the HK ChvG and position of the amino acids defining the truncated proteins. (B-C) Localisation of ChvG WT and truncated versions in cells grown in PYE and imaged on PYE agarose pads (B) or in cells grown in PYE, washed in M2G and imaged on M2G agarose pads (PYE → M2G) (C). Liquid cultures and pads were supplemented with 0.1 % xylose. Demograph data represent cells sorted from short to long length with no less than 300 cells per sample. Scale bars in microscopy images correspond to 1 μm.

## Discussion

The *α*-proteobacterial TCS ChvGI has conserved cell envelope regulatory functions (Lamontagne *et al*., 2007; Heckel *et* al., 2014; Ratib *et al*., 2018; Vallet *et al*., 2020; Stein *et al*., 2021), but the exact perceived signals that trigger the ChvG-dependent phosphorylation of ChvI remained unknown. We found that hyperosmotic shock and antibiotic treatment harming PG crosslinking stimulates ChvI phosphorylation. Also, we showed that other antibiotics targeting PG synthesis proteins induce *chvIG* expression and impair fitness of the *ΔchvI* mutant. On top of this, we showed that ChvI regulates expression of genes related to PG synthesis, such as *ftsN* and *dipM*, and to osmotic and oxidative stress regulation, such as *nepR*, likely explaining why ChvGI system is required for viability in these stressful conditions. Other targets related to cell envelope architecture, identified by ChIP-seq and RNA-seq experiments, include genes coding for the *β*-barrel assembly machinery (Bam) complex; components of the Tol system; several outer membrane proteins (OMP) such as RasFa, CCNA_03820 and CCNA_01956, and several TBDR proteins; and the cytoplasmic FtsZ-binding protein ZapA. Altogether, our results support a role for ChvGI in *C. crescentus* as a safeguard of cell envelope homeostasis by sensing cell wall-related damages and regulating the expression of genes related to cell division and envelope architecture. In the absence of a functional ChvGI system, *C. crescentus* cannot propagate in minimal media except if the TonB-dependent outer membrane protein ChvT or if NtrX is concomitantly inactivated (Stein *et al.,* 2021). ChvGI is known to indirectly down-regulate ChvT by directly activating the expression of the sRNA ChvR (Fröhlich *et al*., 2018). Our ChIP-seq data showed that ChvI also likely regulates *chvT* expression directly by binding to its promoter region. This dual – transcriptional and post-transcriptional – control indicates the ChvT outer membrane protein is an important player in cell envelope homeostasis. Interestingly, NtrYX has been also described as a cell envelope regulator in *α*-proteobacteria. NtrYX controls succinoglycan and exopolysaccharide (EPS) production, and salt stress response in *S. meliloti* (Wang *et al*., 2013; Calatrava *et al*., 2017). In *Rhodobacter sphaeroides,* NtrYX confers resistance to membrane disruptive agents and regulates the expression of genes coding for PG and EPS synthesis enzymes, lipoproteins and cell division proteins (Lemmer *et al.,* 2017; Lemmer *et al*., 2020). Thus, confirming that *ntrX* inactivation in a *ΔchvI* background partially restored growth in the presence of high concentration of osmolytes further supports the importance of the NtrZXY system, together with ChvGI, in the cell envelope stress response in *α*-proteobacteria.

In *C. crescentus*, the general stress response (GSR) sigma factor SigT is also activated upon osmotic imbalance thanks to a complex network comprising the sRNA (GsrN), the anti-SigT regulator (NepR), the histidine phosphotransferases (LovK and PhyK) and the RR (MrrA, LovR and PhyR) (Alvarez-Martinez *et al*., 2007; Lourenço *et al*., 2011; Foreman *et al*., 2012; Lori *et al*., 2018; Tien *et al*., 2018). In steady-state conditions, NepR impedes SigT-dependent transcription. Upon either oxidative stress or osmotic imbalance, the histidine phosphotransferase MrrA activates PhyK to further phosphorylate PhyR. Once phosphorylated, PhyR∼P interacts with NepR to release SigT from inhibition and allow the expression the SigT-dependent regulon. Interestingly, we found that ChvGI and GSR are interconnected but this connection is counterintuitive. Indeed, we observed (i) that deletion of *sigT* led to higher *chvI* promoter activity and (ii) that ChvI directly represses *phyR* expression, suggesting that ChvI and SigT antagonise each other while being sensitive to hyperosmotic conditions. However, we cannot exclude the possibility that the cell envelope homeostasis is sufficiently disrupted in both single mutants to correspondingly activate the other functional system, *i.e.* ChvGI in *ΔsigT* through P*_chvIG_* and SigT in *ΔchvI* through P*_phyR_*. The antagonistic regulation might allow the response of the systems to different levels of osmotic imbalances. For instance, activation of ChvGI at low hypertonic conditions could down-regulate GSR whereas SigT would be sensitive to higher salts concentrations to which ChvGI is turned down. In agreement with this hypothesis, we found that SigT is essential in more severe osmotic scenarios compared to ChvI.

In addition, constitutive activation of a cell envelope stress response system can be detrimental. Overexpression of a phosphomimetic *chvI* variant has been shown to complement growth, but also to result in dysfunctional filamentous growth in minimal media (Stein *et al.,* 2021). The importance of negative regulation of cell envelope stress response systems has been lucidly demonstrated in *E. coli* with the sigma factor E (*σ*^E^), the inner membrane stress regulator Cpx or the regulator of capsule synthesis Rcs (Missiakas *et al*., 1997; Pogliano *et al*., 1998; De Las Peñas *et al*., 2003; Grabowicz & Silhavy. 2017), Indeed, overactivation of *σ*^E^ causes overexpression of sRNAs inhibiting expression of integral OMPs, therefore weakening the outer membrane integrity (Nicoloff *et al*., 2017). Likewise, constitutive Cpx activation disturbs cell division and morphology causing mis-localisation of FtsZ as well as overexpression of L,D-transpeptidase enzyme LdtD (Pogliano *et al*., 1998; Delhaye *et al.,* 2016). Similarly, overactivation of the RR RcsB results in overexpression of the small outer membrane lipoprotein OsmB, which is toxic through yet unknown mechanisms (Grabowicz & Silhavy. 2017). In any of the previous cases, hyperstimulation can lead to deregulation of cell envelope components.

We showed that ChvGI also regulates sensitivity to the antibiotic mecillinam, cefsulodin and moenomycin which inhibit PG transpeptidation. Previously, Vallet *et al*. (2020) showed a mutant for the *chvIG-hprk* operon failed to grow when exposed to vancomycin, which also impedes PG crosslinking by directly interacting with the D-Ala-D-Ala moiety of the PG precursors. Here, we showed that ChvI was phosphorylated when exposed to vancomycin but not to mecillinam. However, although both antibiotics inhibit the transpeptidation step of PG synthesis, they induced *chvIG* expression in different time windows. In addition, these antibiotics cross the outer membrane with different efficiency. For instance, vancomycin is likely efficiently transported by the TBDR ChvT, thereby reaching high concentration in the periplasm in minutes. It is therefore possible that ChvGI is systematically activated upon inhibition of PG transpeptidation during cell elongation but with different sensitivity depending on the transport rate for each antibiotic.

In contrast to the stressors mentioned above, treatment with cefixime, cefotaxime and sodium deoxycholate did not show differences in cell viability between WT and *ΔchvIG-hprk* cells (Vallet *et al*., 2020). Furthermore, the activation of *chvR* expression upon cefotaxime or sodium deoxycholate exposure was shown to be ChvI-independent. In addition, we showed here that *ΔchvI* cells were not more sensitive to treatment with SDS, polymyxin B or the MreB inhibitor A22. Considering that MreB is a key cytoplasmic component of the elongasome complex, of which PBP2 and other proteins are part, it is therefore likely that ChvGI rather senses specific targets in the periplasm to assess cell envelope stress. We propose that ChvG phosphorylation is stimulated by hyperosmotic conditions or antibiotics targeting PG transpeptidation by specifically sensing the pressure or the integrity of the peptidic crosslinks of PG instead of glycan strands. In such a scenario, the periplasmic sensor domain of ChvG could interact with components of the elongasome or bind peptide bridges of the PG. Notwithstanding that, other stress response systems, yet to be discovered or characterised in the context of the aforementioned stressors, might be involved in responding to the damages that do not activate ChvGI.

We observed that ChvG has a patchy-spotty localisation pattern when grown in complex and synthetic minimal media, while it relocates as foci near mid-cell upon transition from complex to minimal media. The patchy-spotty localisation pattern has been described for multiple PG-related proteins in *C. crescentus*, including MreB, MreC, Pal, PBP1A, PBP2, RodA, TolB (Figge *et* al., 2004; Divakaruni *et al.,* 2005; Werner *et al*., 2009; Hocking *et al.,* 2012; Billini *et al*., 2019). Moreover, RodA, PBP2, PBP1A, which are proteins involved in PG polymerization and crosslinking, have been reported to relocate at mid-cell upon osmotic upshift in a FtsZ-dependent but MreB-independent manner (Hocking *et al.,* 2012). It would be interesting to check whether this ChvG localisation patterns are conserved among *α*-proteobacteria, in particular in Rhizobiales since they do not encode MreB orthologs. In *E. coli*, it has been reported that the PBP2 periplasmic portion physically interacts with those of PBP1A and RodA to mediate PG assembly and to ensure proper cell elongation (Banzhaf *et al*., 2012; van der Ploeg *et al*., 2015). It is yet to determine whether ChvG interacts with PBP2, PBP1a, RodA and/or others with similar localisation patterns. Nonetheless, it is tempting to speculate that ChvG interacts and co-localises with PG-related enzymes as a cell envelope safeguard system in *α*-proteobacteria. A recent study in the Gram-positive bacterium *Bacillus thuringiensis* showed that upon treatment with the antibiotic cefoxitin, the putative PBP protein PbpP derepresses the extracytoplasmic function sigma factor P (*σ*^P^) which increases resistance to *β*-lactams (Nauta *et al*., 2021). Future experiments will aim to determine the ChvG interactome under cell envelope undisturbed and threatening conditions, and dissect its activation mechanism as well as its connection with the general stress response, not only in *C. crescentus* but also in other *α*-proteobacteria.

## Acknowledgments

We are grateful to Clare Kirkpatrick, Sean Crosson and Benjamin Stein for willingness to discuss and share data on their ChvGI-related projects. Also, we would like to thank members of the URBM of the University of Namur for their comments and suggestions on this project, especially Dr. Angéline Reboul for her comments and advices on the microscopy analysis, Jérôme Coppine for constructing some strains and members of the Hallez lab for their comments on the manuscript. This work was supported by the Fonds de la Recherche Scientifique – FNRS (F.R.S. – FNRS) with a Welbio Starting Grant (WELBIO-CR-2019S-05) to R.H. A.Q-Y. was supported by a postdoctoral fellowship from the University of Namur (UNamur). R.H. is a Research Associate of F.R.S. – FNRS.

## Author Contributions

A.Q-Y. and R.H. conceived and designed the experiments. A.Q-Y. performed all the experiments except otherwise stated. A.M. purified the ChvI protein and helped with some plasmid’s constructs. A.Q-Y. and R.H. analyzed the data. A.Q-Y. and R.H. wrote the paper.

