## Supplemental Tables S1-S5 for "The two-component system ChvGI maintains cell envelope homeostasis in *Caulobacter crescentus*"

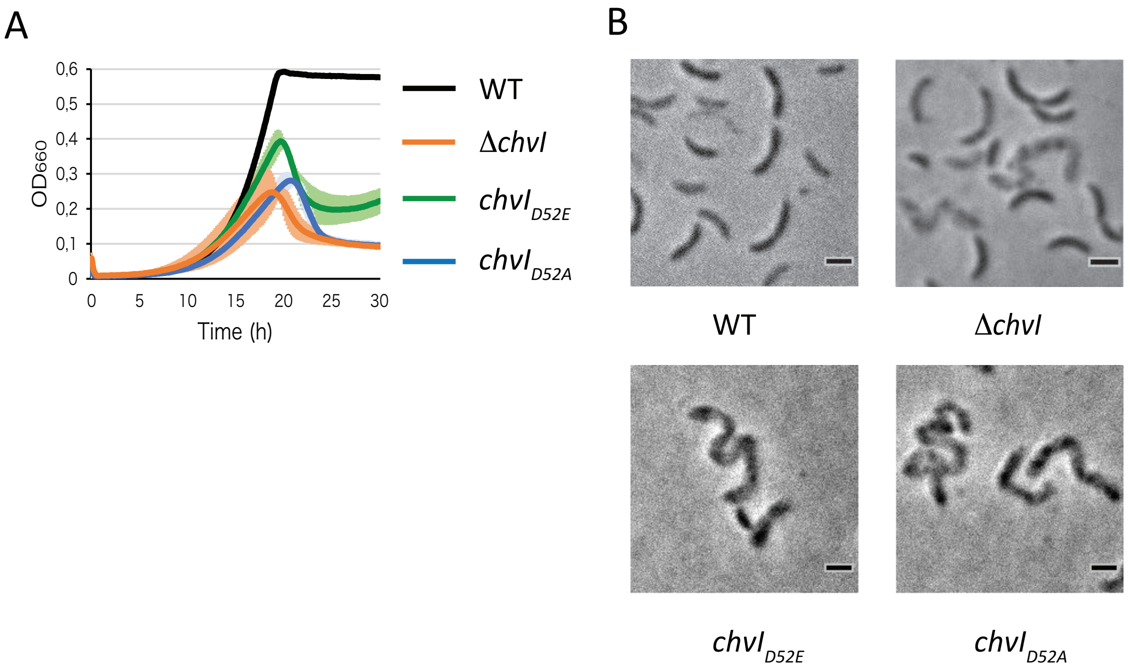


**Figure S1. Growth and morphology of *chvI* mutant strains in synthetic minimal media.** (A) Growth of WT, *chvI*$_{D52E}$ and *chvI*$_{D52A}$ mutant strains in M2G. The data represent the average values of biological replicates (n=3, error bars show standard deviation). (B) Morphology of WT, Δ*chvI*, *chvI*$_{D52E}$ and *chvI*$_{D52A}$ cells after 48 hrs of incubation in M2G. Scale bars in microscopy images correspond to 1 μm.

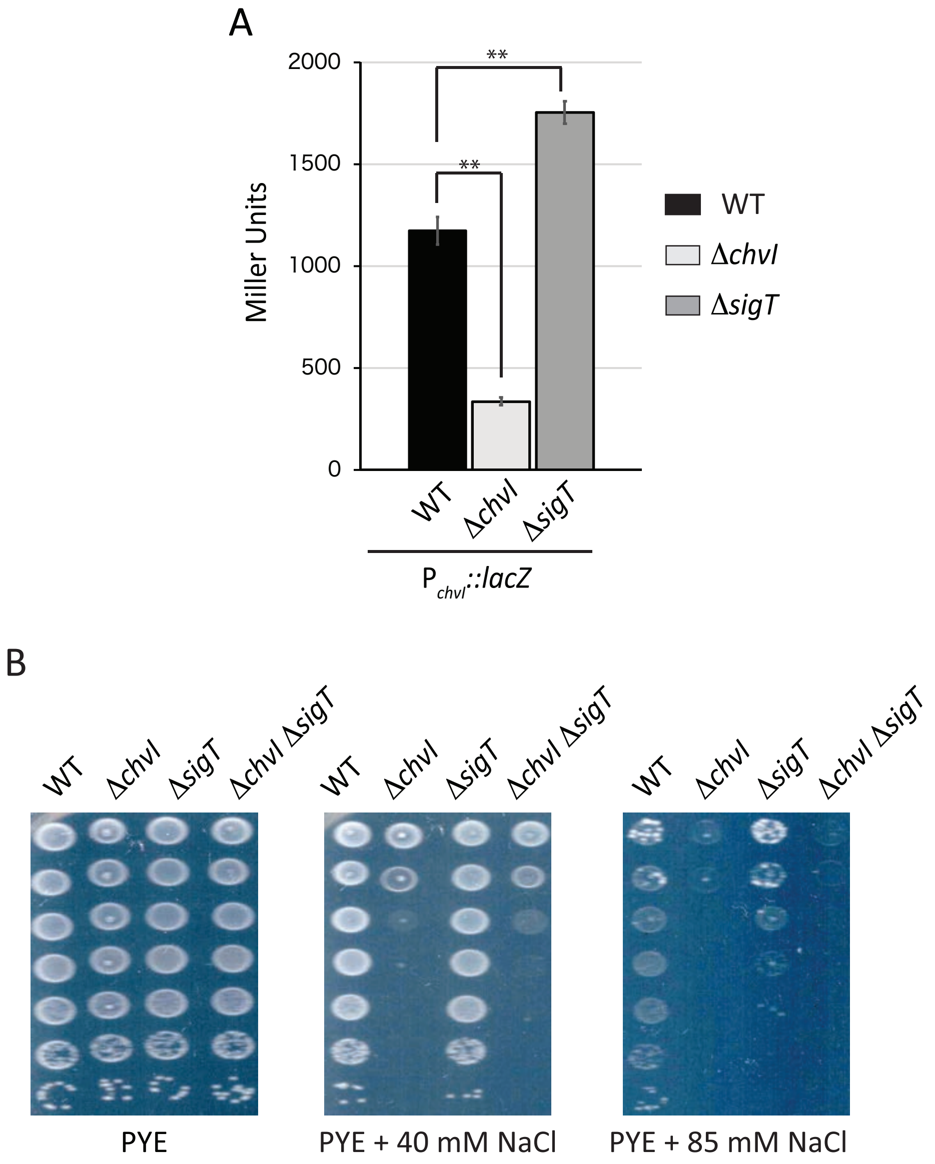


**Figure S2. *chvI* and *sigT* are interconnected.** (A) Activity of the *chvI* promoter $P_{chvI}$ (in Miller Units) in WT (black bars), Δ*chvI* (light grey bars) Δ*sigT* (dark grey bars) cells grown in PYE. The data represent the average values of biological replicates (n=3, error bars show standard deviation). ** = $p < 0.01$ Single factor ANOVA analysis of β-galactosidase activity. (B) Viability of single Δ*chvI* and Δ*sigT* mutants in PYE supplemented with 40 mM and 85 mM NaCl. Images are representative of three biological replicates.

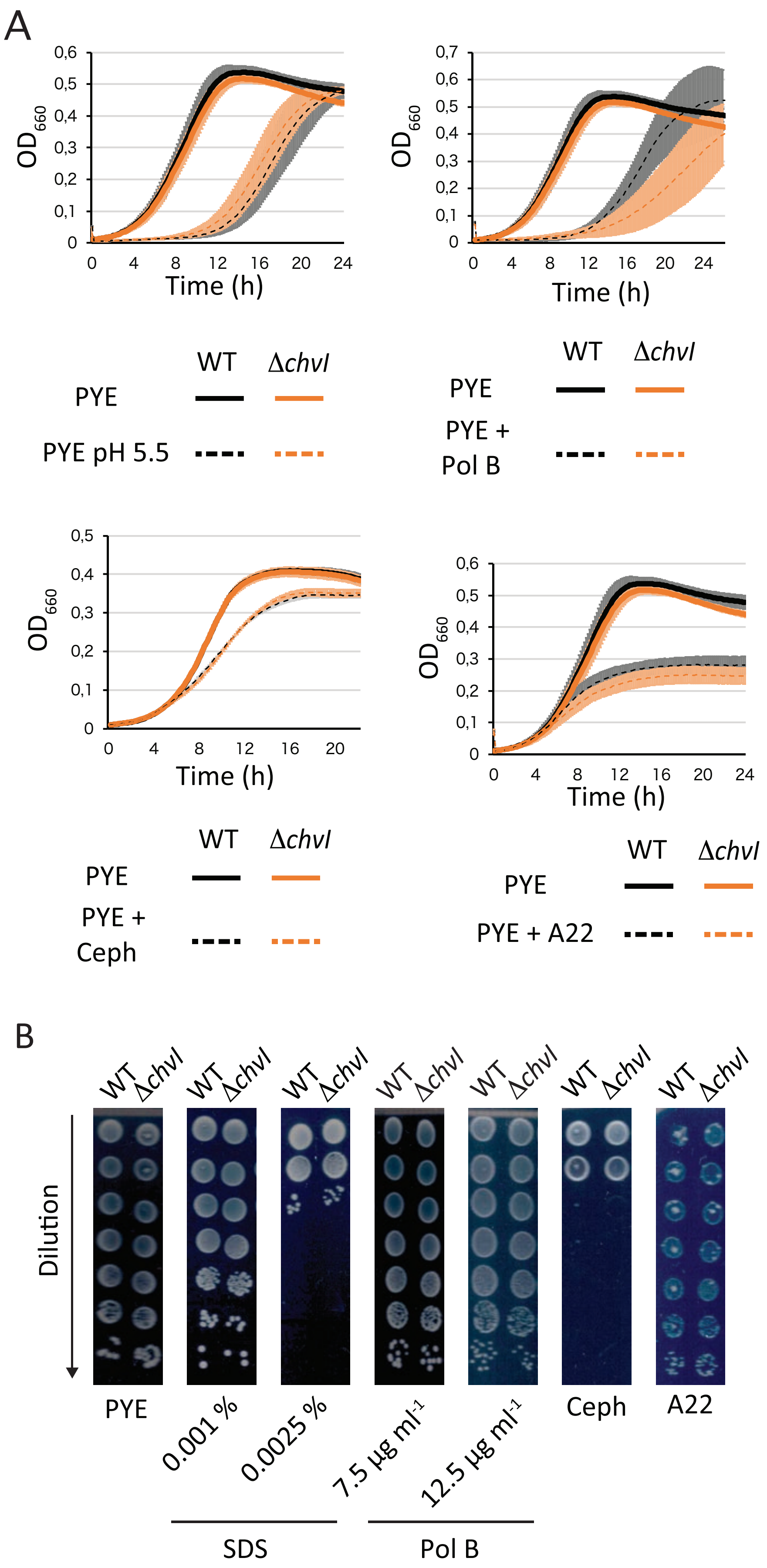

A
OD660
Time (h)
WT
Δchvl
PYE
PYE pH 5.5
PYE + Pol B
PYE + Ceph
PYE + A22
B
Dilution
PYE
0.001 %
0.0025 %
7.5 µg ml-1
12.5 µg ml-1
Ceph
A22
SDS
Pol B

**Figure S3. Δ*chvI* is not sensitive to every cell envelope stress.** (A) Growth of WT (black) and Δ*chvI* (orange) cells in PYE with (dashed lines) or without (solid lines) acidic stress pH 5.5, polymixin B (Pol B) and cephalexin (Ceph). The data represent the average values of biological replicates (n=3, error bars show standard deviation). (B) Viability of Δ*chvI* cells on plates supplemented with 0.001 % or 0.0025% sodium dodecyl sulfate (SDS), Pol B, Ceph or A22. Images are representative of three biological replicates.

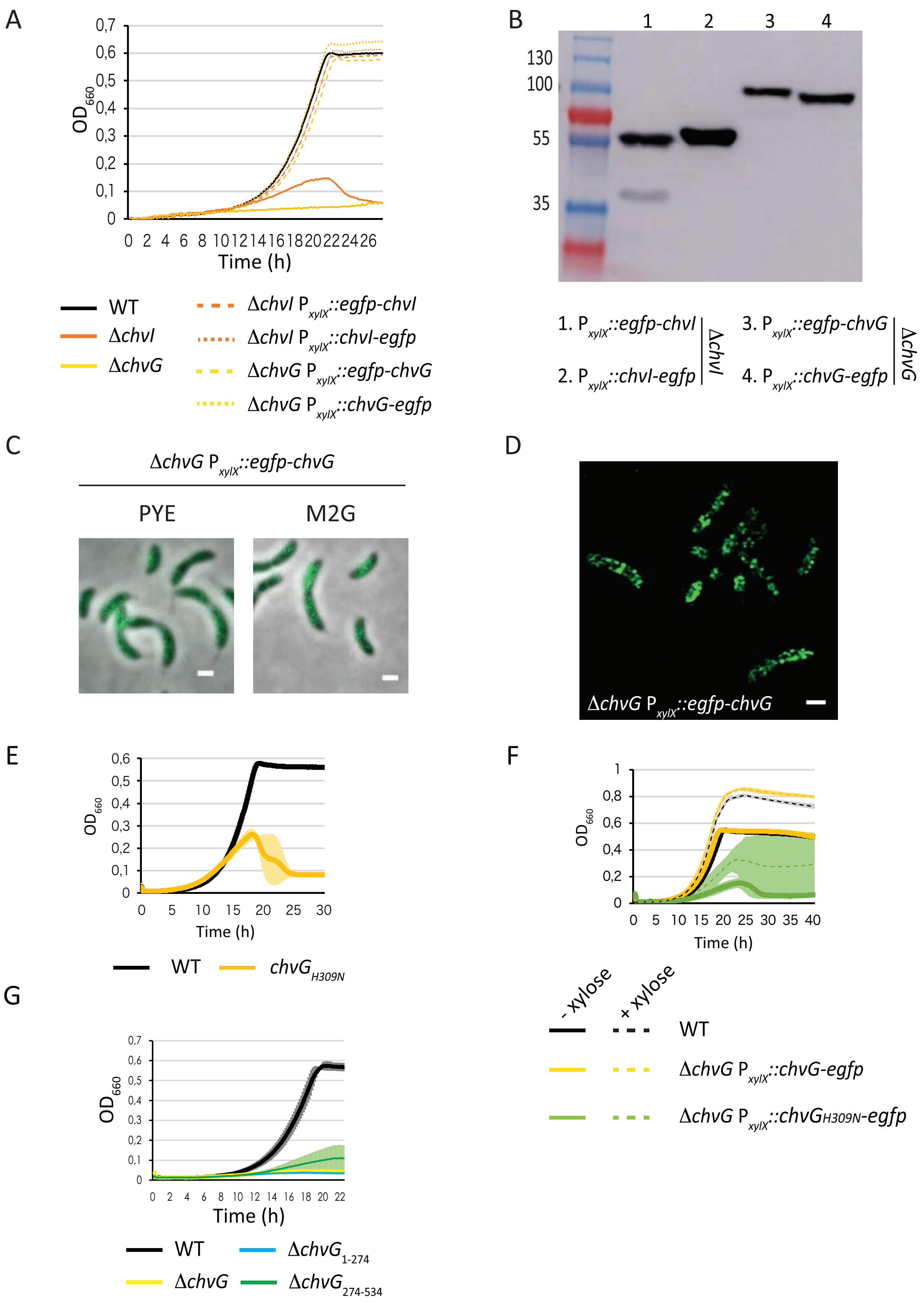
A
0,7
0,6
0,5
0,4
0,3
0,2
0,1
0
OD660
0 2 4 6 8 10 12 14 16 18 20 22 24 26
Time (h)
WT
ΔchvI
ΔchvG
ΔchvI PxylX::egfp-chvI
ΔchvI PxylX::chvI-egfp
ΔchvG PxylX::egfp-chvG
ΔchvG PxylX::chvG-egfp
B
1 2 3 4
130
100
55
35
1. PxylX::egfp-chvI
2. PxylX::chvI-egfp
ΔchvI
3. PxylX::egfp-chvG
4. PxylX::chvG-egfp
ΔchvG
C
ΔchvG PxylX::egfp-chvG
PYE
M2G
D
ΔchvG PxylX::egfp-chvG
E
0,6
0,5
0,4
0,3
0,2
0,1
0
OD660
0 5 10 15 20 25 30
Time (h)
WT
chvGH309N
F
1
0,8
0,6
0,4
0,2
0
OD660
0 5 10 15 20 25 30 35 40
Time (h)
- xylose
+ xylose
WT
ΔchvG PxylX::chvG-egfp
ΔchvG PxylX::chvGH309N-egfp
G
0,7
0,6
0,5
0,4
0,3
0,2
0,1
0
OD660
0 2 4 6 8 10 12 14 16 18 20 22
Time (h)
WT
ΔchvG1-274
ΔchvG
ΔchvG274-534

**Figure S4. ChvG relocates from a patchy-spotty pattern to mid-cell upon osmotic upshift.** (A) Growth of Δ*chvI* and Δ*chvG* mutants complemented with *chvI* and *chvG* N- and C-terminal eGFP fusions. (B) Immunodetection of eGFP fusions with GFP antibodies. The expected molecular weights for eGFP, eGFP-ChvI, ChvI-eGFP, eGFP-ChvG and ChvG-eGFP are 26.94, 54.33, 55.75, 85.88, 86.06 KDa, respectively. (C) Localisation of eGFP-ChvG in a Δ*chvG* background grown overnight in PYE and M2G and imaged in PYE and M2G agarose pads, respectively. (D) Confocal microscopy images of Δ*chvG* cells expressing eGFP-ChvG cells grown in PYE. (E-F) Growth upon endogenous (E) and ectopic expression (F) of $chvG_{H309N}$ in M2G. (G) Impact of deletions in sensor ($\Delta chvG_{274\text{-}534}$) and catalytic ($\Delta chvG_{1\text{-}274}$) domains of ChvG. The data represent the average value of biological replicates (n=3, error bars show standard deviation. Expression of eGFP fusions from $P_{xylX}$ was induced with 0.1 % xylose.
