## Supplemental Figures S1-S4 for "The two-component system ChvGI maintains cell envelope homeostasis in *Caulobacter crescentus*"

### Supplementary tables, methods and references

#### Table S1. Bacterial strains.

| Name | Genetic Information | Reference |
|---|---|---|
| | **Bacteria** | |
| ***Caulobacter crescentus*** | | |
| WT | NA1000, laboratory strain | |
| Δ*chvI* | NA1000 Δ*chvI* | This study |
| Δ*chvG* | NA1000 Δ*chvG* | This study |
| Δ*chvIG* | NA1000 Δ*chvIG* | This study |
| Δ*chvT* | NA1000 Δ*chvT* | This study |
| Δ*chvI* Δ*chvT* | NA1000 Δ*chvI*, Δ*chvT* | This study |
| Δ*ntrX* | NA1000 Δ*ntrX* | This study |
| Δ*chvI* Δ*ntrX* | NA1000 Δ*chvI*, Δ*ntrX* | This study |
| Δ*sigT* | NA1000 Δ*sigT* | This study |
| Δ*xylX* | NA1000 Δ*xylX* | Stephens *et al.,* 2007 |
| $chvI_{D52E}$ | NA1000 $chvI_{D52E}$ | This study |
| $chvI_{D52A}$ | NA1000 $chvI_{D52A}$ | This study |
| $chvG_{H309N}$ | NA1000 $chvG_{H309N}$ | This study |
| Δ*chvI chvI* | NA1000 Δ*chvI* $P_{xylX}$::*chvI*, $Tet^R$ | This study |
| Δ*chvI* $chvI_{D52E}$ | NA1000 Δ*chvI* $P_{xylX}$::$chvI_{D52E}$, $Tet^R$ | This study |
| Δ*chvI* $chvI_{D52A}$ | NA1000 Δ*chvI* $P_{xylX}$::$chvI_{D52A}$, $Tet^R$ | This study |
| Δ*chvI chvI* Δ*xylX* | NA1000 Δ*chvI* $P_{xylX}$::*chvI* Δ*xylX*, $Tet^R$ | This study |
| Δ*chvIG chvI* Δ*xylX* | NA1000 Δ*chvIG* $P_{xylX}$::*chvI* Δ*xylX*, $Tet^R$ | This study |
| WT $P_{dipM}$ | NA1000 pMR15-$P_{dipM}$ | This study |
| Δ*chvI* $P_{dipM}$ | NA1000 Δ*chvI* pMR15-$P_{dipM}$ | This study |
| WT $P_{ftsN}$ | NA1000 pMR15-$P_{ftsN}$ | This study |
| Δ*chvI* $P_{ftsN}$ | NA1000 Δ*chvI* pMR15-$P_{ftsN}$ | This study |
| WT $P_{nepR}$ | NA1000 pMR15-$P_{nepR}$ | This study |
| Δ*chvI* $P_{nepR}$ | NA1000 Δ*chvI* pMR15-$P_{nepR}$ | This study |
| WT $P_{phyR}$ | NA1000 pMR15-$P_{phyR}$ | This study |
| Δ*chvI* $P_{phyR}$ | NA1000 Δ*chvI* pMR15-$P_{phyR}$ | This study |

| | | |
|---|---|---|
| Δ*chvI chvI-egfp* | NA1000 Δ*chvI* $P_{xylX}$::*chvI-egfp*, $Kan^R$ | This study |
| Δ*chvI egfp-chvI* | NA1000 Δ*chvI* $P_{xylX}$::*egfp-chvI*, $Kan^R$ | This study |
| Δ*chvG chvG-egfp* | NA1000 Δ*chvG* $P_{xylX}$::*chvG-egfp*, $Kan^R$ | This study |
| Δ*chvG egfp-chvG* | NA1000 Δ*chvG* $P_{xylX}$::*egfp-chvG*, $Kan^R$ | This study |
| Δ*chvG chvG-mcherry* | NA1000 Δ*chvG* $P_{xylX}$::*chvG-chy*, $Tet^R$ | This study |
| Δ*chvG* $chvG_{1\text{-}114}$*-mcherry* | NA1000 Δ*chvG* $P_{xylX}$:: $chvG_{1\text{-}114}$*-chy*, $Tet^R$ | This study |
| Δ*chvG* $chvG_{1\text{-}274}$*-mcherry* | NA1000 Δ*chvG* $P_{xylX}$:: $chvG_{1\text{-}274}$*-chy*, $Tet^R$ | This study |
| Δ*chvG* $chvG_{273\text{-}534}$*-mcherry* | NA1000 Δ*chvG* $P_{xylX}$:: $chvG_{273\text{-}534}$*-chy*, $Tet^R$ | This study |
| ***Escherichia coli*** | | |
| MT607 | *pro*-82 *thi*-I *hsdR17* (r-m+) *supE44 recA56* | Casadaban & Cohen. 1980 |
| Top10 | *F- mcrA Δ(mrr-hsdRMS-mcrBC) φ80lacZΔM15 ΔlacX74 nupG recA1 araD139 Δ(ara-leu)7697 galE15 galK16 rpsL($Str^R$) endA1* $\lambda^-$ | Life technology |

#### Table S2. Plasmids

| Plasmids | |
|---|---|
| **Name** | **Reference** |
| pNPTS138 | M. R. Alley, Imperial College London (UK), unpublished |
| pNPTS138-Δ*chvI* | This study |
| pNPTS138-Δ*chvG* | This study |
| pNPTS138-Δ*chvIG* | This study |
| pNPTS138-$chvI_{D52E}$ | This study |
| pNPTS138-$chvI_{D52A}$ | This study |
| pNPTS138-$chvG_{H309N}$ | This study |
| pNPTS138-Δ$chvG_{1\text{-}274}$ | This study |
| pNPTS138-Δ$chvG_{274\text{-}534}$ | This study |
| pNPTS138-Δ*chvT* | This study |
| pNPTS138-Δ*ntrX* | This study |

| | |
|---|---|
| pNPTS138-Δ*sigT* | This study |
| pMR15 | Gober & Shapiro, 1992 |
| pMR15-$P_{dipM}$ | This study |
| pMR15-$P_{ftsN}$ | This study |
| pMR15-$P_{nepR}$ | This study |
| pMR15-$P_{phyR}$ | This study |
| pET-28a | Novagen |
| pET-28a-*chvI* | This studychv |
| pXC5 | Thanbichler *et al.,* 2007 |
| pXC5-*chvI* | This study |
| pXC5-$chvI_{D52A}$ | This study |
| pXC5-$chvI_{D52E}$ | This study |
| pXGFPC-2 | Thanbichler *et al.,* 2007 |
| pXGFPC-2 *chvI* | This study |
| pXGFPC-2 *chvG* | This study |
| pXGFPC-2 $chvG_{H309N}$ | This study |
| pXGFPN-2 | Thanbichler *et al.,* 2007 |
| pXGFPN-2 *chvI* | This study |
| pXGFPN-2 *chvG* | This study |
| pXCHYC-5 | Thanbichler *et al.,* 2007 |
| pXCHYC-5 *chvG* | This study |
| pXCHYC-5 $chvG_{1-114}$ | This study |
| pXCHYC-5 $chvG_{1-274}$ | This study |
| pXCHYC-5 $chvG_{273-534}$ | This study |

#### Table S3. Oligos

| ID | Sequence |
|---|---|
| 592 | cttagtcaagcttcttgaagccatgcccgtcac |
| 648 | cgacgaaaccgatcggatcc |
| 649 | cttagtcgaattcgaggacgagacggatagagc |
| 650 | cttagtcgaattcttgggcctcaaccgcaacac |
| 651 | cttagtcaagcttcgacgtaggaattggcgacc |

| | |
|---|---|
| 1013 | cgggaatggcgatcttctgc |
| 1014 | tcgaattcgtcaatgagcgtgatcgcg |
| 1015 | tcaggtcgacccggaattcg |
| 1016 | tcaagcttgctgggcggcgataatctcg |
| 2055 | cctaagtaactaaggatcctagtggtcgccggtcgcttg |
| 2056 | cctaagtaactaagaattcgtccgacctgttccgcgac |
| 2057 | cctaagtaactaagaattcagcgattgagcgccttgcgg |
| 2058 | cctaagtaactaaaagcttcgtcggcatagaccacgc |
| 2155 | ttagttacttaggcatatggctaccgttatcgcgaagcc |
| 2156 | cctaagtaactaagagctcgcgaaagtcacgacgcgatc |
| 2157 | ttagttacttaggcatatggccgcgatcacgctcattga |
| 2158 | cctaagtaactaagagctccgcagctcgttgaggatcaa |
| 2173 | ttagttacttaggggatcccgacctccttcttgggcttgc |
| 2174 | cctaagtaactaagaattcgcgagaccgaagcgacgatg |
| 2177 | aactaaggatcctcagacgtccgagatccctg |
| 2179 | aactaaggtaccagaaccgcaaggacgcgaac |
| 2180 | tcctaagtaactaaaagcttacgagcccgatcatgccttg |
| 2405 | ggatccccgggtacatatgatggccgcgatcacgctcatt |
| 2406 | cctaagtaactaaggtacctcaggcttcgcgataacggtagc |
| 2411 | cctaagtaactaaggtacccaggcttcgcgataacggta |
| 2563 | ttagttacttagggaattcagaaccgcaaggacgcgaac |
| 2568 | ggtagaattctcgcgcgcgtccggcaattcga |
| 2569 | aactaaggtaccttggctaccgttatcgcgaagcc |
| 2570 | aactaaggtaccgccgcgatcacgctcattgacg |
| 2576 | cttcgccgccgacgttgcgaacgagatcaagaatccgct |
| 2577 | agcggattcttgatctcgttcgcaacgtcggcggcgaag |
| 2578 | gtcacggccgaagctagctctgttgttcattccgcgctcg |
| 2579 | tgcaggatatctggatccactgggtctcgacgatctgacgggcgatcg |
| 2580 | ggacctagtgatcctggccgtgaagatgccgcgcatgga |
| 2581 | tccatgcgcggcatcttcacggccaggatcactaggtcc |
| 2582 | ggacctagtgatcctggaagtgaagatgccgcgcatgga |
| 2583 | tccatgcgcggcatcttcacttccaggatcactaggtcc |
| 2584 | gtcacggccgaagctagctagtttctggtgacccgtgc |
| 2585 | tgcaggatatctggatccacaacggtagccaacgccgtacag |

| | |
|---|---|
| 2773 | gtaactaaggtaccgttccgagaggtcgggaagc |
| 2870 | tcacggccgaagctagcgaattcgtggatcaccgaggatcacagagggtc |
| 2871 | gaggaagaaacgacggttgtcctcaaaagtacgcgc |
| 2872 | acttttgaggacaaccgtcgtttcttcctcggct |
| 2873 | ggcccgtgcaattgaagccggctggcgccaaagctgttgaggacgagatcgc |
| 3131 | cttagggaattctccgagaggtcgggaagcgag |
| 3132 | gtaactaacatatgtcggaacggaaggacgaactg |
| 3133 | cttagggaattctgggcgtcgaacagccgcg |
| 3135 | gtaactaaggtaccaggcttcgcgataacggtagc |
| 3136 | gtaactaaggtacctcggaacggaaggacgaactgg |
| 3170 | ttagttacttaggtctagagtgtagagatcggcgggctg |
| 3171 | cctaagtaactaagaattccgggtcgatacgtgtccttcac |
| 3174 | ttagttacttaggtctagacgggtgctggagatcggcac |
| 3175 | cctaagtaactaactcgagagcgtctcctttcccgtcggag |
| 3176 | cctaagtaactaactcgagagcgtctcctttcccgtcggag |
| 3177 | cctaagtaactaactcgagctcagctaggctccaaaactcgg |
| 3182 | ttagttacttaggtctagaggcgtagcggcggatgtaagg |
| 3183 | cctaagtaactaactcgaggtcctcgacgccgaagttcatcgg |
| 3184 | ttagttacttaggtctagattcatccagcgccgcagcgc |
| 3185 | cctaagtaactaactcgaggacccctcccgattctgtgtc |

#### Table S4. ChvI direct targets by ChIP-seq.

| Top hit | Peak Coord | | Reads | Gene(s)/ operon Up-stream | Peaks within gene/ operons | Gene(s)/ operon Down-stream | Description |
|---|---|---|---|---|---|---|---|
| | Start | End | | | | | |
| 1 | 3260512 | 3261530 | 0,087344 | CCNA_03108 | | CCNA_03109 | TonB-dependent outer membrane receptor ChvT; hypothetical protein |
| 2 | 3945553 | 3946727 | 0,076339 | CCNA_03779 | | CCNA_R0088-CCNA_R0092 | hypothetical cytosolic protein; Minimal medium sRNA; Minimal medium sRNA |
| 3 | 2236111 | 2236942 | 0,033142 | CCNA_02086-83 | | | sporulation domain-containing protein FtsN; anhydromuramoyl-peptide exo-beta-N-acetylglucosaminidase; scpAB; scpB |
| 4 | 2874707 | 2875488 | 0,026194 | | | CCNA_02715 | MASE1-family sensor histidine kinase |
| 5 | 2226770 | 2227559 | 0,022965 | CCNA_02075 | | | peptidoglycan-specific endopeptidase, M23 family DipM |
| 6 | 3239945 | 3240736 | 0,021142 | CCNA_03090-89 | | CCNA_03091 | acetyl-coenzyme A carboxylase carboxyl transferase subunit alpha; hypothetical protein; hypothetical protein NstA |
| 7 | 3488054 | 3488951 | 0,018241 | CCNA_03313 | | CCNA_03314 | hypothetical protein; ferredoxin-NADP reductase |
| 8 | 2973941 | 2974697 | 0,017617 | CCNA_02816-15 | CCNA_02817 | CCNA_02818 | hypothetical protein; ice nucleation protein; retrotransposon-related protein; hypothetical protein |
| 9 | 612854 | 613625 | 0,016826 | | | CCNA_00585 | hypothetical protein |
| 10 | 1258370 | 1259098 | 0,016724 | | | CCNA_01155 | TonB-dependent outer membrane receptor |
| 11 | 252038 | 252843 | 0,016579 | CCNA_00236 | | CCNA_00237-239 | hypothetical protein; two-component response regulator chvI; two-component sensor histidine kinase chvG; HPR(SER) kinase/phosphatase HprK |
| 12 | 3713510 | 3714250 | 0,016064 | CCNA_03557 | | | hypothetical protein |

| | | | | | | | |
|---|---|---|---|---|---|---|---|
| 13 | 2901131 | 2901845 | 0,015265 | CCNA_02740-39 | | CCNA_02741 | transposase; transposase; hypothetical protein |
| 14 | 165136 | 165885 | 0,015134 | | CCNA_R0100 | CCNA_00158 | small non-coding RNA chvR; DNA replication and repair protein recF |
| 15 | 1699274 | 1699977 | 0,015006 | CCNA_01583 | CCNA_03954 | | phosphate-binding protein PtsS; hypothetical protein |
| 16 | 792750 | 793440 | 0,014526 | CCNA_00735 | | | hypothetical protein BamF |
| 17 | 1347680 | 1348576 | 0,014025 | CCNA_01221-20 | | | acyl carrier protein; serine palmitoyltransferase |
| 18 | 1162578 | 1163287 | 0,01284 | | | CCNA_01060 | type I protein secretion ATP-binding protein RsaD |
| 19 | 2479442 | 2480102 | 0,011866 | CCNA_02339-38--CCNA_02337-36 | | | methyltransferase; hypothetical protein--nonspecific lipid-transfer protein; lysophospholipase L2 |
| 20 | 3069157 | 3069876 | 0,010922 | CCNA_02910 | CCNA_02911 | CCNA_02912-13 | TonB-dependent receptor; hypothetical protein; prolyl 4-hydroxylase, alpha subunit; hypothetical protein |
| 21 | 1928596 | 1929220 | 0,010865 | | CCNA_01799-01806 (on CCNA_01804) | CCNA_01805-06 | aKG dehydrogenase complex (gluthatione reductase); dihydrolipoamide dehydrogenase; ferritin-like domain protein |
| 22 | 3484061 | 3484706 | 0,009773 | | CCNA_03307 | CCNA_03308 | hypothetical protein; hypothetical protein |
| 23 | 534703 | 535377 | 0,009536 | | | CCNA_00519 | hypothetical protein |
| 24 | 41569 | 42207 | 0,009342 | CCNA_00039-38 | CCNA_00040 | | hypothetical protein; hypothetical protein; hypothetical protein |
| 25 | 1454091 | 1454704 | 0,009226 | | | CCNA_01341 | endopeptidase degP |
| 26 | 564302 | 564985 | 0,009145 | CCNA_R0110 | | CCNA_00546 | small non-coding RNA; hypothetical protein |
| 27 | 3375179 | 3375782 | 0,008372 | CCNA_03212-11 | CCNA_03213 | | hypothetical protein; L-Ala-D/L-Glu racemase; putative polyhydroxyalkanoic acid system protein |
| 28 | 3952328 | 3952937 | 0,008074 | CCNA_03784 | | CCNA_03997 | transporter; amelogenin/CpxP-related protein |
| 29 | 3247461 | 3248133 | 0,00797 | CCNA_03096 | | | TonB-dependent receptor |
| 30 | 3986130 | 3986741 | 0,007779 | | | CCNA_03820-CCNA_R0199 | outer-membrane lipoproteins carrier protein; small non-coding RNA |
| 31 | 2194116 | 2194719 | 0,007758 | | | CCNA_02048 | TonB-dependent receptor |
| 32 | 3877328 | 3877963 | 0,007322 | CCNA_03711-10 | | | ribosome-associated factor Y; nitrogen regulatory EIIA_Ntr protein |

| | | | | | | | |
|---|---|---|---|---|---|---|---|
| 33 | 2141171 | 2141741 | 0,006809 | CCNA_01996-91 | CCNA_01996-91 (on CCNA_01994) (2nd peak) | | undecaprenyl pyrophosphate synthetase; CCNA_01993_mmpA;CCNA_01991_bamB; (1-deoxy-D-xylulose 5-phosphate reductoisomerase) |
| 34 | 455092 | 455711 | 0,006801 | | | CCNA_00451 | TonB-dependent receptor |
| 35 | 1544729 | 1545304 | 0,006777 | CCNA_01427 | CCNA_01428 | CCNA_01429 | lipoprotein, SmpA/OmlA family BamE; hypothetical protein; ubiquinol-cytochrome C reductase chaperone |
| 36 | 2256297 | 2256867 | 0,006607 | CCNA_02106 | CCNA_02107 | | TonB-dependent outer membrane receptor; hypothetical protein |
| 37 | 3000054 | 3000679 | 0,006602 | CCNA_02846 | | | endopeptidase degP |
| 38 | 1050847 | 1051441 | 0,006496 | CCNA_00973 | | CCNA_00974 | hypothetical protein; OAR protein precursor |
| 39 | 2738974 | 2739543 | 0,006018 | CCNA_02588-87 | | CCNA_02589-90 | aminopeptidase; PAS-family sensor histidine kinase; bolA-related transcriptional regulator; hypothetical protein |
| 40 | 1886800 | 1887365 | 0,005996 | CCNA_01759-57 | | | peptidyl-prolyl cis-trans isomerase; 4-hydroxythreonine-4-phosphate dehydrogenase PdxA; dimethyladenosine transferase |
| 41 | 2039583 | 2040156 | 0,005756 | CCNA_01896 | | | hypothetical protein |
| 42 | 2881391 | 2881931 | 0,005756 | | | CCNA_02721 | peptidase, M16 family |
| 43 | 3535655 | 3536200 | 0,005729 | CCNA_03357-56 | | CCNA_03358-59 | hypothetical protein ZauP; cytosolic protein "zapA binds to FtsZ"; glyceraldehyde 3-phosphate dehydrogenase; phosphoglycerate kinase |
| 44 | 242520 | 243188 | 0,005717 | CCNA_00224 | CCNA_00224 (2nd peak) | CCNA_00225 | TonB-dependent receptor; hypothetical protein |
| 45 | 967538 | 968086 | 0,005642 | CCNA_00889 | | | hypothetical protein |
| 46 | 1564844 | 1565427 | 0,005602 | | | CCNA_01451-52 | cold shock protein cspD; hypothetical protein |
| 47 | 1197427 | 1197961 | 0,005537 | CCNA_01090-88 | | | hypothetical protein; hypothetical protein; hypothetical protein |
| 48 | 1122849 | 1123416 | 0,005526 | | CCNA_R0128 | CCNA_01034-35-CCNA_R0129 | small non-coding RNA; TonB-dependent outer membrane receptor; gamma-glutamyltranspeptidase; small non-coding RNA |
| 49 | 2232515 | 2233085 | 0,005522 | CCNA_02082-80 | | | Sec-independent protein translocase protein tatA; Sec-independent protein |

| | | | | | | | |
|---|---|---|---|---|---|---|---|
| | | | | | | | translocase protein tatB; tatC |
| 50 | 695830 | 696415 | 0,005434 | | CCNA_00644-48 (on CCNA_00645) | CCNA_00646-48 | nitrate transport permease protein nrtB; ntrD; nasT; hypothetical protein-nrtABD-nasT (nrtA) |
| 51 | 180921 | 181505 | 0,004855 | | | CCNA_00169 | hypothetical protein |
| 52 | 2945701 | 2946286 | 0,00462 | CCNA_02788 | CCNA_02789 | CCNA_02790-91 | transcriptional regulator, Xre family; transcriptional regulator, Xre family; RNA polymerase ECF-type sigma factor; FecR family protein |
| 53 | 1549888 | 1550402 | 0,004539 | CCNA_01437 | | CCNA_01438 | transglycosylase associated protein; heme O monooxygenase |
| 54 | 1616543 | 1617088 | 0,00452 | CCNA_01505 | | | hypothetical protein |
| 55 | 1216395 | 1216899 | 0,004354 | | | CCNA_01111-12 | DnaA-related protein; hypothetical protein |
| 56 | 765235 | 765739 | 0,004325 | CCNA_00707 | CCNA_00708 | | hypothetical protein; cobaltochelatase cobT subunit) |
| 57 | 3522574 | 3523074 | 0,004268 | CCNA_03341-38 | CCNA_03339 (2nd peak) | | TolQ protein; TolR protein; TolA protein; TolB protein ; TolA protein |
| 58 | 3843579 | 3844106 | 0,004185 | CCNA_03681-79 | CCNA_03680 (2nd peak) | CCNA_03682-84 | ABC transporter ATP-binding protein; aminopeptidase N; PaaI thioesterase family protein; aminopeptidase N; fumarylpyruvate hydrolase; maleylpyruvate isomerase; carbonic anhydrase |
| 59 | 178909 | 179528 | 0,004178 | | | CCNA_00169 | hypothetical protein |
| 60 | 3781267 | 3781819 | 0,004061 | | CCNA_R0079-CCNA_R0093 | | Minimal medium sRNA; Minimal medium sRNA CrfA |
| 61 | 1555254 | 1555735 | 0,00405 | CCNA_01443 | | | hypothetical protein |
| 62 | 3637314 | 3637821 | 0,004001 | CCNA_03909-CCNA_03469 | | CCNA_03471 | conserved hypothetical protein (Tat_SS); arginyl-tRNA synthetase; 4-hydroxy-3-methylbut-2-enyl diphosphate reductase |
| 63 | 2156776 | 2157275 | 0,003909 | CCNA_02010-11 | CCNA_02009 | CCNA_02008-06 | hypothetical protein; methylmalonyl CoA epimerase; hypothetical protein; prolyl-tRNA synthetase; lipoprotein releasing system transmembrane protein lolE; lolD |
| 64 | 691664 | 692203 | 0,003831 | CCNA_00639-37 | CCNA_00640 | | heme:hemopexin-binding protein (OMP pore |

| | | | | | | | |
|---|---|---|---|---|---|---|---|
| | | | | | | | involved in secretion); heme:hemopexin-binding protein; tetratricopeptide repeat family protein; hypothetical protein |
| 65 | 3603188 | 3603663 | 0,003822 | CCNA_03439-37 | | | hypothetical protein; hypothetical protein; hypothetical protein |
| 66 | 2896957 | 2897435 | 0,003762 | CCNA_02734-33 | | | serine acetyltransferase; glutaminyl-peptide cyclotransferase |
| 67 | 500803 | 501284 | 0,003745 | CCNA_00484 | | CCNA_00486 | hypothetical protein; TonB-dependent receptor (far from the peak) |
| 68 | 1732749 | 1733229 | 0,003713 | | | CCNA_01612-17 | rod shape-determining protein mreB; mreCD-PBP2-rodA-acetyltransferase |
| 69 | 454451 | 454985 | 0,003582 | | | CCNA_00451 | TonB-dependent receptor |
| 70 | 4024504 | 4024998 | 0,003514 | | CCNA_03862-63 | | regulatory protein tenI/hypothetical protein |
| 71 | 790240 | 790696 | 0,003505 | | | CCNA_00733-34 | GumN superfamily protein; GumN superfamily protein |
| 72 | 1429727 | 1430196 | 0,00343 | | | CCNA_01303 | hypothetical protein |
| 73 | 2349925 | 2350380 | 0,003395 | CCNA_02200 | CCNA_02201 | CCNA_02202 | cytochrome c-family protein; hypothetical protein; hypothetical protein |
| 74 | 304080 | 304562 | 0,003387 | CCNA_00290 | | CCNA_00291 | autotransporter protein; phosphate regulon sensor protein phoR |
| 75 | 1536254 | 1536721 | 0,003301 | | CCNA_01418 | CCNA_01419-20 | transcriptional regulator; serine hydroxymethyltransferase; putative regulatory protein |
| 76 | 430325 | 430793 | 0,003137 | CCNA_00417-16 | | CCNA_00419 | hypothetical protein; hypothetical protein; fasciclin domain cell surface protein |
| 77 | 3442131 | 3442603 | 0,003112 | | CCNA_03272 | CCNA_03276 | Two-component sensor histidine kinase; glycerophosphoryl diester phosphodiesterase |
| 78 | 2454191 | 2454653 | 0,003072 | CCNA_02308 | | CCNA_02309 | hypothetical protein; EF hand domain protein |
| 79 | 1141919 | 1142377 | 0,003058 | | | CCNA_01047 | TonB-dependent receptor |
| 80 | 2974719 | 2975201 | 0,003056 | CCNA_02817 | CCNA_02818 | | retrotransposon-related protein; hypothetical protein |
| 81 | 2752682 | 2753121 | 0,003055 | | CCNA_02606 | CCNA_R0169 | hybrid sensor histidine kinase/receiver domain protein; small non-coding RNA |

| | | | | | | | |
|---|---|---|---|---|---|---|---|
| 82 | 135210 | 135656 | 0,002953 | | CCNA_00125-24 (on CCNA_00125) | | hypothetical protein; (amino acid efflux permease) |
| 83 | 3624407 | 3624855 | 0,002903 | | | CCNA_03461-60 | hypothetical protein; hypothetical protein |
| 84 | 1147876 | 1148333 | 0,002858 | | | CCNA_01051 | TonB-dependent receptor |
| 85 | 396583 | 397038 | 0,002834 | CCNA_R0106-CCNA_00379-77 | | | small non-coding RNA; thiol:disulfide interchange protein dsbA; dsbA; smc |
| 86 | 994043 | 994475 | 0,002828 | CCNA_00914 | CCNA_00915 | | acetyltransferase; xylene monooxygenase electron transfer component |
| 87 | 3882100 | 3882553 | 0,002768 | CCNA_03717-16 | | | ribonuclease D; hypothetical protein |
| 88 | 2214429 | 2214855 | 0,002759 | CCNA_02063 | CCNA_02064 | | lipoprotein, ComL family BamD; UDP-3-O-(3-hydroxymyristoyl) N-acetylglucosamine deacetylase LpxC |
| 89 | 1478849 | 1479279 | 0,002683 | | CCNA_01361-63 (on CCNA_01363) | | hypothetical protein; PhoPQ |
| 90 | 3703971 | 3704409 | 0,002666 | | | CCNA_03548 | carboxy-terminal processing protease precursor |
| 91 | 178384 | 178813 | 0,002656 | CCNA_00166 | | CCNA_00167 | HvyA; bis(5 -nucleosyl)-tetraphosphatase (symmetrical) |
| 92 | 2993773 | 2994203 | 0,002619 | CCNA_02839 | | CCNA_02840 | glutamate-ammonia-ligase adenylyltransferase; transcriptional regulator mecR-family |
| 93 | 3876566 | 3877035 | 0,002539 | CCNA_03711-10 | | | ribosome-associated factor Y; nitrogen regulatory EIIA_Ntr protein |
| 94 | 2938829 | 2939253 | 0,002516 | | CCNA_02780-81 (on CCNA_02780) | CCNA_02781 | hypothetical protein-(hypothetical protein); hypothetical protein |
| 95 | 4752 | 5166 | 0,002394 | CCNA_R0095 | CCNA_00007 | CCNA_00008 | small non-coding RNA; SSU ribosomal protein S20P; chromosomal replication initiator protein DnaA (2nd peak) |
| 96 | 677252 | 677655 | 0,002384 | | CCNA_00629-34 (on CCNA_00630) | CCNA_00631-34 | chemotaxis locus; cheW; cheYII; cheBII; chemotaxis protein methyltransferase |
| 97 | 3773948 | 3774354 | 0,002316 | | CCNA_03619 (1st peak) | | zinc metalloprotease |
| 98 | 338335 | 338762 | 0,002314 | CCNA_00325-24 | | CCNA_00326 | hypothetical protein; TolR protein; vegetatible incompatibility protein HET-E-1 |
| 99 | 494580 | 495002 | 0,002313 | | CCNA_00478 | | hypothetical protein |
| 100 | 382549 | 382961 | 0,002234 | CCNA_00365 | | CCNA_00366-68 | ornithine decarboxylase; phosphonates transport |

| | | | | | | | |
|---|---|---|---|---|---|---|---|
| | | | | | | | ATP-binding protein phnC; phnDE |
| 101 | 1757586 | 1757974 | 0,002113 | | CCNA_01637-38 (on CCNA_01637) | CCNA_01638 | topoisomerase IV subunit A; beta-lactamase family protein |
| 102 | 1629515 | 1629939 | 0,002081 | | CCNA_01521 | | hypothetical protein |
| 103 | 1291350 | 1291740 | 0,00204 | | | CCNA_01171 | histidine phosphotransferase ShpA |
| 104 | 3355295 | 3355689 | 0,001989 | CCNA_03194-92 | | CCNA_03987-CCNA_03195 | integral membrane protein; lactoylglutathione lyase; transposase; hypothetical protein; RNA polymerase sigma-32 factor RpoH |
| 105 | 224803 | 225195 | 0,001979 | CCNA_00209 | | CCNA_00210 | undecaprenyl-phosphomannose:protein mannosyltransferase; TonB-dependent receptor |
| 106 | 2587945 | 2588320 | 0,001906 | CCNA_02446 | CCNA_02448-46 (on CCNA_02446) | | glutathionylspermidine synthase family; hypothetical protein; putative alpha helical protein; (hypothetical protein) |
| 107 | 3513837 | 3514198 | 0,001735 | CCNA_03333-21 | | | PAS-family sensor histidine kinase; dihydropteroate synthase |
| 108 | 3777206 | 3777566 | 0,001727 | CCNA_03621-20 | | CCNA_03622-23 | transcriptional regulator, AraC family; hypothetical protein; beta-lactamase repressor; transcriptional regulator |
| 109 | 416610 | 416979 | 0,001723 | CCNA_00398 | | CCNA_00399 | hypothetical protein; zinc metallohydrolase, glyoxalase II family |
| 110 | 1850311 | 1850669 | 0,001671 | | | CCNA_01724-25 | hypothetical protein; PQQ enzyme repeat family protein BamA |
| 111 | 1726556 | 1726909 | 0,001659 | CCNA_01606-05 | | CCNA_01607 | Soj/ParA-related ATPase protein; NADPH-dependent FMN reductase family protein; hypothetical protein |
| 112 | 1004971 | 1005341 | 0,00165 | CCNA_00928-27 | | CCNA_00929-32 | transcriptional regulator, GntR family; hypothetical protein; Riboflavin-specific deaminase family protein; riboflavin synthase alpha chain; GTP cyclohydrolase II/3,4-dihydroxy-2-butanone-4-phosphate synthase; ribH |
| 113 | 1781770 | 1782120 | 0,001641 | CCNA_01660 | | CCNA_R0041 | hypothetical protein; tRNA Lys |
| 114 | 1979527 | 1979884 | 0,001633 | | CCNA_01848-47 | CCNA_01846-45 | qoxD; cytochrome c oxidase assembly protein |

| | | | | | | | |
|---|---|---|---|---|---|---|---|
| | | | | | | | Surf1; two-component sensor histidine kinase; two-component response regulator |
| 115 | 1170928 | 1171290 | 0,00162 | | | CCNA_01067 | type I secretion outer membrane protein RsaFa |
| 116 | 2177494 | 2177842 | 0,001617 | CCNA_02033-21 | | CCNA_R0045 | NADH-quinone oxidoreductase chain A; nuo locus; tRNA Asp |
| 117 | 1193070 | 1193424 | 0,001615 | CCNA_01086 | | CCNA_01087 | GTP-binding protein lepA; hypothetical protein EipA |
| 118 | 170561 | 170923 | 0,001582 | | | CCNA_00161 | hypothetical protein |
| 119 | 292222 | 292580 | 0,001549 | | CCNA_00279 | | NAD(P)H dehydrogenase (quinone) |
| 120 | 1301611 | 1301951 | 0,001543 | CCNA_01181 | | CCNA_01182-84 | hypothetical protein; glutathione S-transferase; hypothetical protein; phosphohydrolase (MutT/nudix family protein) |
| 121 | 215293 | 215650 | 0,001539 | CCNA_00200 | | CCNA_00201 | 2OG-Fe(II) oxygenase superfamily protein; outer membrane protein |
| 122 | 2565751 | 2566099 | 0,001485 | CCNA_02421-19 | | | TonB accessory protein exbB; TonB accessory protein exbD; TonB1 protein |
| 123 | 250732 | 251078 | 0,001484 | | CCNA_00236 | CCNA_00235 | hypothetical protein; hypothetical protein |
| 124 | 3294176 | 3294520 | 0,001483 | CCNA_03142 | | CCNA_R0183 | RNA polymerase sigma factor rpoD; small non-coding RNA |
| 125 | 3518400 | 3518747 | 0,001473 | CCNA_03336-35 | | | Tol system periplasmic component YbgF; peptidoglycan-associated lipoprotein pal |
| 126 | 3505822 | 3506177 | 0,001472 | CCNA_03323 | CCNA_03325-24 | CCNA_03326 | phosphoserine aminotransferase; hypothetical protein-hypothetical protein, two-component sensor histidine kinase |
| 127 | 3663926 | 3664267 | 0,001455 | CCNA_03505 | | CCNA_03506 | peroxiredoxin; transcriptional regulator, algH |
| 128 | 532821 | 533156 | 0,001437 | CCNA_00517-16 | | | peptidyl-tRNA hydrolase; integral membrane protein |
| 129 | 2888992 | 2889325 | 0,001428 | CCNA_02725-24 | | CCNA_R0063 | choline dehydrogenase; oxidoreductase; Minimal medium sRNA |
| 130 | 2976656 | 2976999 | 0,001419 | | CCNA_02820 | CCNA_R0174 | hypothetical protein; small non-coding RNA |
| 131 | 361976 | 362315 | 0,001384 | | CCNA_R0104 | | small non-coding RNA |
| 132 | 3774557 | 3774890 | 0,001364 | | CCNA_03619 | | zinc metalloprotease (2nd peak) |

| | | | | | | | |
|---|---|---|---|---|---|---|---|
| 133 | 2808186 | 2808515 | 0,001352 | | CCNA_02654 | | hypothetical protein |
| 134 | 3683077 | 3683410 | 0,001306 | CNA_03525-24 | | CCNA_03526 | hypothetical protein; di-/tripeptide transporter; hypothetical protein |
| 135 | 389985 | 390314 | 0,001278 | CCNA_00374-70 | | | putative ATP synthase protein I; ATP synthase |
| 136 | 1387719 | 1388046 | 0,001268 | CCNA_R0137 | | CCNA_01261-63 | small non-coding RNA; periplasmic multidrug efflux lipoprotein precursor; cation/multidrug efflux pump, AcrB family; pyruvate dehydrogenase E1 component |
| 137 | 445284 | 445619 | 0,001256 | CCNA_00438 | | CCNA_00439-41 | hypothetical protein; methyl-accepting chemotaxis protein McpA; STAS domain protein; cheYI |
| 138 | 260751 | 261081 | 0,001253 | CCNA_00248 | | CCNA_00249 | sensor histidine protein kinase; SCO1/SenC family protein |
| 139 | 488007 | 488331 | 0,001237 | CCNA_00472-70 | | | GDP-mannose 4,6 dehydratase; GDP-L-fucose synthase; O-antigen polymerase |
| 140 | 3744733 | 3745086 | 0,001231 | CCNA_03590-88 | | CCNA_03591 | hypothetical protein; RNA polymerase ECF-type sigma factor sigT; two-component sensor histidine kinase PhyK; hybrid sigma factor/two-component receiver protein phyR |
| 141 | 1610571 | 1610886 | 0,001227 | | CCNA_01498-1500 (upstream of CCNA_01499) | CCNA_01500 | DNA-dependent DNA polymerase III subunit alpha; (acetyl-CoA acetyltransferase); hypothetical protein; hypothetical protein |
| 142 | 3803576 | 3803896 | 0,00122 | CCNA_03644-41 | | | succinate dehydrogenase cytochrome B-556 subunit; succinate dehydrogenase sdhA-D |
| 143 | 1503556 | 1503876 | 0,00121 | CCNA_01386 | | CCNA_01387-88 | hypothetical protein; tRNA (uracil-5-)-methyltransferase; methyltransferase |
| 144 | 3266144 | 3266460 | 0,001185 | CCNA_R0179 | CCNA_03116-17 (on CCNA_03116) | | small non-coding RNA; (hypothetical protein); cytosolic protein BacB |
| 145 | 140117 | 140433 | 0,001181 | | CCNA_00130-29 (on CCNA_00130) | CCNA_00131 | Kup system potassium uptake protein; Rrf2 family protein |
| 146 | 2773217 | 2773527 | 0,001163 | CCNA_03971-CCNA_02623 | | | hypothetical protein; cell division protein FtsZ |

| | | | | | | | |
|---|---|---|---|---|---|---|---|
| 147 | 1850861 | 1851172 | 0,001143 | | CCNA_01724-25 (on CCNA_01724) | | (hypothetical protein)-BamA (2nd peak) |
| 148 | 1774463 | 1774779 | 0,001111 | | | CCNA_01653-54 | peptidyl-prolyl cis-trans isomerase; peptidyl-prolyl cis-trans isomerase |
| 149 | 1061356 | 1061654 | 0,000964 | | | CCNA_00982 | transcriptional regulator |
| 150 | 485653 | 485953 | 0,000907 | | CCNA_00472-70 (on CCNA_00470) | | GDP-mannose 4,6 dehydratase; GDP-L-fucose synthase; (O-antigen polymerase) |
| 151 | 1457875 | 1458161 | 0,000899 | | CCNA_01342-44 (on CCNA_01343) | CCNA_01344 | ATPase, AAA family; (ribosomal large subunit pseudouridine synthase C); hypothetical protein; hypothetical protein |
| 152 | 3754813 | 3755101 | 0,000892 | | | CCNA_03601-02 | hemolysin III-like protein; patatin family phospholipase domain protein |
| 153 | 2140781 | 2141068 | 0,000888 | CCNA_01996-91 | CCNA_01996-91 (on CCNA_01994) (2nd peak) | | undecaprenyl pyrophosphate synthetase; CCNA_01993_mmpA;CCNA_01991_bamB; (1-deoxy-D-xylulose 5-phosphate reductoisomerase) |
| 154 | 314272 | 314558 | 0,000867 | | CCNA_00300 | CCNA_00301-02 | glyoxalase superfamily protein; phosphotransferase family protein; parathion hydrolase |
| 155 | 4002901 | 4003183 | 0,000856 | CCNA_03837 | | CCNA_03838 | hypothetical protein; gluconate 2-dehydrogenase/glyoxylate reductase/hydroxypyruvate reductase |
| 156 | 1840007 | 1840287 | 0,000853 | | | CCNA_01708 | hypothetical protein |
| 157 | 2475301 | 2475580 | 0,000852 | CCNA_02333 | | CCNA_02334 | lipid A biosynthesis lauroyl acyltransferase; transcriptional regulator of stalk biogenesis staR |
| 158 | 851247 | 851526 | 0,000802 | CCNA_00787-CCNA_03938 | | CCNA_00788 | chemotaxis motA protein; hypothetical protein; transporter, major facilitator superfamily |
| 159 | 3837932 | 3838209 | 0,000799 | CCNA_03677 | | CCNA_03678 | CoA-transferase family III protein; hypothetical protein |
| 160 | 116073 | 116346 | 0,000794 | CCNA_00104 | | CCNA_00105 | orotidine 5 -phosphate decarboxylase; MarC family integral membrane protein |
| 161 | 1235496 | 1235769 | 0,000788 | | CCNA_01130-32 (on CCNA_01131) | CCNA_01132 | fliR-flhB-cckA, sensory transduction histidine kinase/receiver protein CckA |

| | | | | | | | |
|---|---|---|---|---|---|---|---|
| 162 | 576012 | 576282 | 0,00077 | | CCNA_00560-64 (on CCNA_00560) | CCNA_00562-64 | phosphohistidine phosphatase SixA; conserved cytosolic protein; dihydroorotate dehydrogenase; SNARE-associated family membrane protein |
| 163 | 2294051 | 2294325 | 0,00077 | CCNA_02136-33 | | | zwf; pgl-edd-glk |
| 164 | 824298 | 824559 | 0,000694 | CCNA_00767-66 | | CCNA_03936-CCNA_00768-69 | hypothetical protein; cobalamin adenosyltransferase family protein; hypothetical protein; hypothetical protein; transcriptional regulator, TetR family |
| 165 | 354535 | 354798 | 0,000681 | | | CCNA_00342-43 | 2-oxoglutarate dehydrogenase E1 component; dihydrolipoamide succinyltransferase component (E2) of 2-oxoglutarate dehydrogenase complex |
| 166 | 2335439 | 2335700 | 0,000664 | | | CCNA_03962 | hypothetical protein |
| 167 | 4003258 | 4003517 | 0,000648 | CCNA_03837 (2nd peak) | | CCNA_03838 | hypothetical protein ; gluconate 2-dehydrogenase/glyoxylate reductase/hydroxypyruvat e reductase |
| 168 | 1184120 | 1184376 | 0,000624 | CCNA_01079 | | CCNA_01080 | hypothetical protein; hypothetical protein |
| 169 | 3397416 | 3397668 | 0,000591 | CCNA_03988 | CCNA_03235-33-CCNA_03988 (on CCNA_03233) | | hypothetical protein |
| 170 | 743621 | 743870 | 0,000567 | CCNA_00688 | CCNA_00688-87 | CCNA_00689 | hypothetical protein; hypothetical protein; PAS-family hybrid histidine kinase/receiver protein (2nd peak) |
| 171 | 2942165 | 2942414 | 0,000567 | | CCNA_02783-87 (on CCNA_02783) | CCNA_02784-87 | hypothetical protein; hypothetical protein; guanylate cyclase; hypothetical protein |
| 172 | 707912 | 708159 | 0,000551 | | | CCNA_R0009 | sRNA |
| 173 | 3208948 | 3209195 | 0,000551 | CCNA_03055 | | CCNA_03054 | transcriptional regulator, AraC family; transcriptional regulator, AraC family |
| 174 | 1812973 | 1813219 | 0,000543 | CCNA_01685 | | CCNA_01686-88 | hemimethylated DNA-binding protein yccV; exopolyphosphatase; 23S rRNA Um2552 2 -O-methyltransferase; potassium channel protein |

| | | | | | | | |
|---|---|---|---|---|---|---|---|
| 175 | 2511935 | 2512177 | 0,000525 | CCNA_02368 | | | alpha-glucosidase |
| 176 | 213720 | 213956 | 0,000463 | | CCNA_00199 | | hypothetical protein |
| 177 | 265733 | 265958 | 0,000373 | CCNA_00252 | | CCNA_00253 | multimodular transpeptidase-transglycosylase PbpX (PBP 1A); aspartate-semialdehyde dehydrogenase |
| 178 | 902058 | 902283 | 0,000365 | | | CCNA_00836 | flagellin |
| 179 | 1860590 | 1860811 | 0,00034 | CCNA_01731-30 | CCNA_01732 | | colicin V production protein; amidophosphoribosyltransferase; DNA repair protein RadA |
| 180 | 673974 | 674198 | 0,000332 | CCNA_00628-27 | | CCNA_00629-34 | methyl-accepting chemotaxis protein; cheAII; cheW; cheYII; cheBII; chemotaxis protein methyltransferase; chemotaxis protein cheY; hypothetical protein |
| 181 | 3253808 | 3254028 | 0,000332 | CCNA_03102-01 | | CCNA_03103-04 | hypothetical protein; integrase/recombinase (XerD/RipX family); shikimate kinase; 3-dehydroquinate synthase |
| 182 | 1205267 | 1205486 | 0,000324 | CCNA_01099 | | | tetracycline resistance protein |
| 183 | 1557589 | 1557807 | 0,000316 | CCNA_01444 | CCNA_01445 | CCNA_01446-47 | poly(3-hydroxyalkanoate) polymerase; hypothetical protein; putative aminotransferase aatC; homoserine dehydrogenase |
| 184 | 451597 | 451817 | 0,000308 | | CCNA_00444 | CCNA_00445-50 | chemotaxix locus |
| 185 | 1397327 | 1397544 | 0,000308 | CCNA_01268 | | CCNA_01269 | cytochrome c; GcrB protein |
| 186 | 1508862 | 1509079 | 0,000308 | | CCNA_01390-91 (on CCNA_01390) | CCNA_01391 | RNA polymerase sigma factor; radical SAM superfamily protein |
| 187 | 3895442 | 3895659 | 0,000308 | CCNA_03726 | | CCNA_03727 | transcriptional regulator; small-conductance mechanosensitive channel mcsS |
| 188 | 840044 | 840259 | 0,000292 | | CCNA_R0014-CCNA_00779 (on CCNA_R0014) | CCNA_00779 | Stat phase sRNA; hypothetical protein |
| 189 | 456564 | 456778 | 0,000284 | | CCNA_00451 | | TonB-dependent receptor |
| 190 | 333063 | 333278 | 0,000219 | CCNA_00317-14 | | CCNA_00318 | GTP-binding protein CgtA; glutamate 5-kinase; 3-polyprenyl-4-hydroxybenzoate decarboxylase; hypothetical protein; hypothetical protein |

**Table S5. Genes identified by RNA-seq down-regulated in the Δ*chvI* mutant upon osmotic stress with 6% sucrose.**

| Top hit | Gene ID | Description | log2 (FC Δ*chvI*/WT ) | P-value | P-adj |
|---|---|---|---|---|---|
| 1 | CCNA_03997 | amelogenin/CpxP-related protein | 8,579868 | 1,10E-239 | 4,43E-236 |
| 2 | CCNA_00237 | two-component response regulator chvI | 7,460508 | 1,34E-191 | 2,72E-188 |
| 3 | CCNA_R0088 | Minimal medium sRNA | 6,136769 | 1,77E-115 | 2,39E-112 |
| 4 | CCNA_01238 | EF-hand domain protein | 5,332779 | 2,91E-107 | 2,95E-104 |
| 5 | CCNA_R0092 | Minimal medium sRNA | 5,227363 | 8,42E-43 | 2,43E-40 |
| 6 | CCNA_00889 | hypothetical protein | 4,883146 | 4,29E-86 | 3,47E-83 |
| 7 | CCNA_R0161 | small non-coding RNA | 4,543405 | 2,92E-29 | 3,81E-27 |
| 8 | CCNA_03987 | hypothetical protein | 4,275498 | 1,49E-46 | 5,03E-44 |
| 9 | CCNA_01660 | hypothetical protein | 4,093043 | 3,23E-48 | 1,19E-45 |
| 10 | CCNA_02817 | retrotransposon-related protein | 4,044014 | 7,82E-51 | 3,16E-48 |
| 11 | CCNA_01087 | cell envelope integrity protein eipA | 4,041426 | 1,91E-53 | 9,67E-51 |
| 12 | CCNA_03308 | hypothetical protein | 3,974113 | 5,99E-30 | 8,66E-28 |
| 13 | CCNA_02309 | EF hand domain protein | 3,947459 | 6,60E-29 | 7,80E-27 |
| 14 | CCNA_03102 | hypothetical protein | 3,784691 | 1,09E-33 | 1,77E-31 |
| 15 | CCNA_03820 | outer-membrane lipoproteins carrier protein | 3,782962 | 8,49E-65 | 5,72E-62 |
| 16 | CCNA_01080 | hypothetical protein | 3,690148 | 1,16E-40 | 2,92E-38 |
| 17 | CCNA_02531 | proline-rich hypothetical protein | 3,486805 | 2,19E-26 | 2,01E-24 |
| 18 | CCNA_03601 | hemolysin III-like protein) | 3,466022 | 2,92E-59 | 1,69E-56 |
| 19 | CCNA_00038 | hypothetical protein | 3,459263 | 3,52E-42 | 9,49E-40 |
| 20 | CCNA_01341 | endopeptidase degP | 3,299589 | 2,05E-52 | 9,23E-50 |
| 21 | CCNA_03602 | patatin family phospholipase domain protein | 3,296568 | 2,80E-38 | 5,96E-36 |
| 22 | CCNA_00687 | hypothetical protein | 3,203373 | 8,72E-39 | 2,08E-36 |
| 23 | CCNA_00039 | hypothetical protein | 3,175555 | 1,20E-23 | 9,91E-22 |
| 24 | CCNA_R0199 | small non-coding RNA | 3,138597 | 1,91E-35 | 3,36E-33 |
| 25 | CCNA_01583 | phosphate-binding protein PtsS | 3,10072 | 7,81E-36 | 1,44E-33 |
| 26 | CCNA_01592 | SN-glycerol-3-phosphate transport ATP-binding protein | 3,084726 | 4,35E-38 | 8,81E-36 |
| 27 | CCNA_02165 | esterase/lipase | 3,081526 | 6,50E-22 | 4,96E-20 |

| | | | | | |
|---|---|---|---|---|---|
| 28 | CCNA_01090 | hypothetical protein | 3,075921 | 1,72E-43 | 5,34E-41 |
| 29 | CCNA_02310 | cellulose 1,4-beta-cellobiosidase | 3,060081 | 7,64E-30 | 1,07E-27 |
| 30 | CCNA_02721 | peptidase, M16 family | 3,050148 | 9,87E-29 | 1,08E-26 |
| 31 | CCNA_03101 | integrase/recombinase (XerD/RipX family) | 3,019754 | 3,27E-25 | 2,88E-23 |
| 32 | CCNA_01067 | type I secretion outer membrane protein RsaFa | 2,98762 | 4,23E-27 | 4,17E-25 |
| 33 | CCNA_01773 | SNARE-associated family membrane protein | 2,986556 | 5,91E-29 | 7,24E-27 |
| 34 | CCNA_02106 | TonB-dependent outer membrane receptor | 2,967977 | 2,22E-28 | 2,30E-26 |
| 35 | CCNA_00888 | hypothetical protein | 2,966904 | 3,27E-20 | 2,41E-18 |
| 36 | CCNA_00686 | peptidase family S41 protein | 2,944474 | 1,82E-26 | 1,75E-24 |
| 37 | CCNA_00484 | hypothetical protein | 2,926154 | 2,15E-26 | 2,01E-24 |
| 38 | CCNA_02308 | hypothetical protein | 2,904541 | 3,41E-29 | 4,31E-27 |
| 39 | CCNA_02846 | endopeptidase degP | 2,873802 | 6,85E-34 | 1,15E-31 |
| 40 | CCNA_01088 | hypothetical protein | 2,86569 | 8,62E-23 | 6,98E-21 |
| 41 | CCNA_02001 | phosphoserine phosphatase | 2,845159 | 6,07E-32 | 9,44E-30 |
| 42 | CCNA_01445 | hypothetical protein | 2,833545 | 1,36E-38 | 3,05E-36 |
| 43 | CCNA_01089 | hypothetical protein | 2,815112 | 9,13E-37 | 1,76E-34 |
| 44 | CCNA_03726 | transcriptional regulator | 2,804138 | 2,20E-28 | 2,30E-26 |
| 45 | CCNA_02219 | hypothetical protein | 2,733332 | 9,54E-26 | 8,58E-24 |
| 46 | CCNA_03338 | TolB protein | 2,699714 | 3,38E-30 | 5,07E-28 |
| 47 | CCNA_00733 | GumN superfamily protein | 2,600268 | 1,73E-29 | 2,33E-27 |
| 48 | CCNA_01446 | putative aminotransferase aatC | 2,593795 | 2,07E-19 | 1,42E-17 |
| 49 | CCNA_02339 | methyltransferase | 2,585514 | 3,45E-28 | 3,49E-26 |
| 50 | CCNA_02889 | peptidyl-prolyl cis-trans isomerase | 2,585088 | 6,91E-29 | 7,80E-27 |
| 51 | CCNA_00546 | hypothetical protein | 2,581795 | 1,17E-13 | 5,42E-12 |
| 52 | CCNA_01505 | hypothetical protein | 2,578745 | 3,66E-11 | 1,42E-09 |
| 53 | CCNA_01386 | hypothetical protein | 2,494249 | 2,47E-24 | 2,08E-22 |
| 54 | CCNA_02865 | phage protein | 2,484474 | 3,01E-14 | 1,52E-12 |
| 55 | CCNA_00735 | hypothetical protein BamF | 2,477619 | 8,96E-18 | 5,58E-16 |
| 56 | CCNA_03909 | conserved hypothetical protein | 2,45932 | 6,94E-29 | 7,80E-27 |
| 57 | CCNA_01653 | peptidyl-prolyl cis-trans isomerase | 2,417407 | 7,40E-16 | 3,99E-14 |
| 58 | CCNA_00123 | 3-hydroxyacyl CoA dehydrogenase | 2,40595 | 1,14E-13 | 5,38E-12 |
| 59 | CCNA_03336 | Tol system periplasmic component YbgF | 2,320795 | 1,74E-24 | 1,50E-22 |
| 60 | CCNA_02201 | hypothetical protein | 2,320121 | 6,67E-12 | 2,75E-10 |

| | | | | | |
|---|---|---|---|---|---|
| 61 | CCNA_03548 | carboxy-terminal processing protease precursor | 2,277761 | 3,32E-16 | 1,84E-14 |
| 62 | CCNA_R0100 | small non-coding RNA chvR | 2,245174 | 1,21E-08 | 3,46E-07 |
| 63 | CCNA_01654 | peptidyl-prolyl cis-trans isomerase | 2,236579 | 3,61E-22 | 2,81E-20 |
| 64 | CCNA_02819 | hypothetical protein | 2,214202 | 8,69E-12 | 3,52E-10 |
| 65 | CCNA_R0106 | small non-coding RNA | 2,189526 | 5,66E-19 | 3,76E-17 |
| 66 | CCNA_01081 | hypothetical protein | 2,177142 | 3,54E-21 | 2,65E-19 |
| 67 | CCNA_01993 | membrane endopeptidase MmpA | 2,173333 | 3,37E-19 | 2,27E-17 |
| 68 | CCNA_01725 | PQQ enzyme repeat family protein BamA | 2,166415 | 1,36E-19 | 9,66E-18 |
| 69 | CCNA_01147 | acetyltransferase | 2,161138 | 5,71E-13 | 2,60E-11 |
| 70 | CCNA_01772 | hypothetical protein | 2,159305 | 1,90E-16 | 1,07E-14 |
| 71 | CCNA_00292 | phosphate transport system permease protein pstC | 2,148154 | 3,56E-22 | 2,81E-20 |
| 72 | CCNA_01344 | hypothetical protein | 2,109334 | 7,12E-20 | 5,14E-18 |
| 73 | CCNA_02202 | hypothetical protein | 2,085797 | 4,21E-14 | 2,08E-12 |
| 74 | CCNA_02338 | hypothetical protein | 2,07658 | 3,46E-16 | 1,89E-14 |
| 75 | CCNA_03212 | hypothetical protein | 2,024365 | 6,94E-12 | 2,83E-10 |
| 76 | CCNA_01427 | lipoprotein, SmpA/OmlA family BamE | 2,013488 | 1,09E-06 | 2,17E-05 |
| 77 | CCNA_03460 | hypothetical protein | 2,006656 | 3,40E-18 | 2,15E-16 |
| 78 | CCNA_03335 | PP-loop family cell cycle control ATPase | 1,996091 | 9,45E-18 | 5,79E-16 |
| 79 | CCNA_01443 | hypothetical protein | 1,974744 | 2,71E-17 | 1,59E-15 |
| 80 | CCNA_01447 | homoserine dehydrogenase | 1,959219 | 2,88E-11 | 1,13E-09 |
| 81 | CCNA_00793 | phosphatidylglycerol glycosyltransferase | 1,957657 | 3,58E-14 | 1,79E-12 |
| 82 | CCNA_01500 | hypothetical protein | 1,953374 | 1,86E-18 | 1,22E-16 |
| 83 | CCNA_03773 | hypothetical protein | 1,952802 | 1,74E-17 | 1,04E-15 |
| 84 | CCNA_00218 | hypothetical protein | 1,94742 | 1,48E-19 | 1,03E-17 |
| 85 | CCNA_00347 | hypothetical protein | 1,889342 | 0,000181 9 | 0,002055 5 |
| 86 | CCNA_00887 | hypothetical protein | 1,873478 | 1,15E-12 | 5,01E-11 |
| 87 | CCNA_01759 | peptidyl-prolyl cis-trans isomerase | 1,862241 | 1,02E-16 | 5,87E-15 |
| 88 | CCNA_01659 | queuine tRNA-ribosyltransferase | 1,859312 | 1,46E-15 | 7,78E-14 |
| 89 | CCNA_01724 | hypothetical protein | 1,854791 | 1,63E-16 | 9,30E-15 |
| 90 | CCNA_01345 | short chain dehydrogenase | 1,825671 | 7,99E-13 | 3,55E-11 |
| 91 | CCNA_01068 | mannosyltransferase | 1,82141 | 2,12E-15 | 1,10E-13 |
| 92 | CCNA_03280 | pyruvate ferredoxin/flavodoxin oxidoreductase family protein | 1,821146 | 1,14E-17 | 6,88E-16 |
| 93 | CCNA_00548 | transposase | 1,813886 | 0,000110 2 | 0,001308 |

| | | | | | |
|---|---|---|---|---|---|
| 94 | CCNA_03083 | hybrid sensor histidine kinase/receiver domain protein | 1,792828 | 2,69E-13 | 1,23E-11 |
| 95 | CCNA_01496 | 2-dehydro-3-deoxyphosphooctonate aldolase | 1,787233 | 1,64E-14 | 8,40E-13 |
| 96 | CCNA_00293 | phosphate transport system permease protein pstA | 1,75902 | 1,94E-15 | 1,02E-13 |
| 97 | CCNA_01636 | alkaline phosphatase | 1,745158 | 1,15E-10 | 4,33E-09 |
| 98 | CCNA_R0074 | small non-coding RNA | 1,742013 | 0,000661 9 | 0,006376 7 |
| 99 | **CCNA_03557** | hypothetical protein | 1,724285 | 2,54E-09 | 8,16E-08 |
| 100 | **CCNA_00398** | hypothetical protein | 1,721562 | 1,71E-12 | 7,28E-11 |
| 101 | **CCNA_02334** | transcriptional regulator of stalk biogenesis staR | 1,718166 | 5,69E-09 | 1,69E-07 |
| 102 | **CCNA_03469** | arginyl-tRNA synthetase | 1,690582 | 7,19E-13 | 3,23E-11 |
| 103 | **CCNA_03461** | hypothetical protein | 1,688066 | 3,43E-06 | 6,14E-05 |
| 104 | **CCNA_00124** | hypothetical protein | 1,685398 | 1,07E-13 | 5,10E-12 |
| 105 | CCNA_03709 | transposase | 1,673242 | 3,44E-06 | 6,14E-05 |
| 106 | CCNA_01385 | predicted rRNA methylase | 1,672307 | 2,05E-09 | 6,70E-08 |
| 107 | CCNA_01907 | ABC transporter ATP-binding protein | 1,658489 | 1,58E-07 | 3,86E-06 |
| 108 | **CCNA_00707** | hypothetical protein | 1,650257 | 2,11E-10 | 7,67E-09 |
| 109 | **CCNA_02063** | lipoprotein, ComL family BamD | 1,649258 | 9,53E-14 | 4,59E-12 |
| 110 | CCNA_02888 | nitrilotriacetate monooxygenase | 1,632957 | 2,29E-08 | 6,27E-07 |
| 111 | **CCNA_00238** | two-component sensor histidine kinase chvG | 1,626522 | 7,18E-14 | 3,50E-12 |
| 112 | **CCNA_02781** | hypothetical protein | 1,625061 | 3,84E-08 | 1,01E-06 |
| 113 | **CCNA_00169** | hypothetical protein | 1,611327 | 3,11E-12 | 1,31E-10 |
| 114 | CCNA_R0121 | small non-coding RNA | 1,603412 | 2,25E-05 | 0,000336 5 |
| 115 | CCNA_03196 | hypothetical protein | 1,558725 | 3,57E-08 | 9,49E-07 |
| 116 | **CCNA_03341** | TolQ protein | 1,544807 | 3,49E-08 | 9,36E-07 |
| 117 | CCNA_00272 | RmuC family protein | 1,53237 | 1,32E-12 | 5,69E-11 |
| 118 | CCNA_03725 | ATP-dependent DNA ligase | 1,522201 | 1,73E-07 | 4,19E-06 |
| 119 | CCNA_R0132 | small non-coding RNA | 1,51777 | 0,000109 2 | 0,001302 9 |
| 120 | CCNA_00784 | heat resistant agglutinin | 1,514111 | 1,98E-05 | 0,000304 6 |
| 121 | **CCNA_01992** | outer membrane protein assembly factor BamB | 1,510435 | 1,07E-12 | 4,73E-11 |
| 122 | **CCNA_03784** | transporter | 1,503962 | 3,22E-05 | 0,000445 8 |
| 123 | CCNA_01501 | protein kinase C-like superfamily protein | 1,497378 | 1,59E-05 | 0,000248 4 |
| 124 | **CCNA_03117** | cytosolic protein BacB | 1,492925 | 1,93E-09 | 6,34E-08 |

| | | | | | |
|---|---|---|---|---|---|
| 125 | CCNA_01994 | 1-deoxy-D-xylulose 5-phosphate reductoisomerase | 1,491637 | 4,05E-06 | 7,06E-05 |
| 126 | CCNA_00547 | PurR-related transcriptional regulator | 1,45568 | 4,07E-09 | 1,26E-07 |
| 127 | CCNA_03506 | transcriptional regulator, algH | 1,45187 | 1,46E-11 | 5,81E-10 |
| 128 | CCNA_02115 | secreted pectate lyase-family protein | 1,442696 | 1,29E-09 | 4,35E-08 |
| 129 | CCNA_00907 | 3-oxoacyl-(acyl-carrier-protein) synthase III | 1,440485 | 3,53E-10 | 1,23E-08 |
| 130 | CCNA_01758 | 4-hydroxythreonine-4-phosphate dehydrogenase PdxA | 1,436887 | 2,18E-10 | 7,89E-09 |
| 131 | CCNA_00116 | phosphoglucosamine mutase | 1,435697 | 2,58E-10 | 9,17E-09 |
| 132 | CCNA_00785 | tRNA (5-aminomethyl-2-thiouridylate) methyltransferase/tRNA (5-carboxymethylaminomethyl-2-thiouridyl | 1,418898 | 4,83E-05 | 0,000638 6 |
| 133 | CCNA_02009 | hypothetical protein | 1,412129 | 8,88E-07 | 1,81E-05 |
| 134 | CCNA_02838 | EF hand domain protein | 1,409921 | 5,98E-07 | 1,29E-05 |
| 135 | CCNA_01497 | ADP-L-glycero-D-manno-heptose-6-epimerase | 1,408935 | 3,85E-09 | 1,20E-07 |
| 136 | CCNA_03710 | nitrogen regulatory EIIA_Ntr protein | 1,407127 | 5,17E-09 | 1,57E-07 |
| 137 | CCNA_01346 | outer membrane protein | 1,404111 | 2,24E-07 | 5,30E-06 |
| 138 | CCNA_03772 | HIT1 protein | 1,399326 | 1,59E-08 | 4,47E-07 |
| 139 | CCNA_03600 | integral membrane protein | 1,399002 | 8,07E-05 | 0,000989 4 |
| 140 | CCNA_03172 | 3-oxoacyl-(acyl-carrier protein) reductase | 1,396147 | 5,36E-07 | 1,17E-05 |
| 141 | CCNA_00645 | nitrate binding protein nrtA | 1,378191 | 1,27E-05 | 0,000201 1 |
| 142 | CCNA_00290 | autotransporter protein | 1,377947 | 2,48E-10 | 8,88E-09 |
| 143 | CCNA_01607 | hypothetical protein | 1,370074 | 5,40E-09 | 1,63E-07 |
| 144 | CCNA_01731 | colicin V production protein | 1,360457 | 4,81E-09 | 1,47E-07 |
| 145 | CCNA_03211 | L-Ala-D/L-Glu racemase | 1,341752 | 3,98E-07 | 8,96E-06 |
| 146 | CCNA_R0128 | small non-coding RNA | 1,341529 | 0,001147 9 | 0,010428 7 |
| 147 | CCNA_02220 | beta-glucosidase | 1,336701 | 6,39E-07 | 1,36E-05 |
| 148 | CCNA_00122 | hypothetical protein | 1,299073 | 5,63E-09 | 1,69E-07 |
| 149 | CCNA_01019 | beta-glucosidase | 1,278072 | 7,01E-08 | 1,83E-06 |
| 150 | CCNA_01613 | rod shape-determining protein mreC | 1,272769 | 3,96E-07 | 8,95E-06 |
| 151 | CCNA_03599 | aspartate-semialdehyde dehydrogenase | 1,248873 | 0,000352 4 | 0,003675 1 |
| 152 | CCNA_03340 | TolR protein | 1,245906 | 3,52E-06 | 6,25E-05 |
| 153 | CCNA_02837 | RNA polymerase ECF-type sigma factor | 1,241975 | 7,74E-05 | 0,000952 7 |
| 154 | CCNA_02820 | hypothetical protein | 1,226276 | 1,63E-06 | 3,13E-05 |
| 155 | CCNA_00378 | thiol:disulfide interchange protein dsbA | 1,222306 | 2,00E-07 | 4,78E-06 |

| | | | | | |
|---|---|---|---|---|---|
| 156 | CCNA_00882 | hypothetical protein | 1,20416 | 0,0002648 | 0,0028645 |
| 157 | CCNA_03995 | hypothetical protein | 1,20192 | 0,0001504 | 0,0017282 |
| 158 | CCNA_01708 | hypothetical protein | 1,191325 | 1,85E-07 | 4,46E-06 |
| 159 | CCNA_00130 | Kup system potassium uptake protein | 1,19066 | 6,78E-05 | 0,0008605 |
| 160 | CCNA_00364 | deoxyhypusine synthase | 1,186074 | 4,65E-05 | 0,0006214 |
| 161 | CCNA_02081 | Sec-independent protein translocase protein tatB | 1,175577 | 3,07E-07 | 7,14E-06 |
| 162 | CCNA_01020 | transcriptional regulator, LacI family | 1,173496 | 4,68E-05 | 0,0006215 |
| 163 | CCNA_01885 | short chain dehydrogenase | 1,17193 | 8,57E-05 | 0,0010477 |
| 164 | CCNA_03092 | cytochrome P450 IVA5 | 1,166815 | 1,43E-07 | 3,52E-06 |
| 165 | CCNA_00121 | hypothetical protein | 1,160166 | 9,85E-07 | 1,98E-05 |
| 166 | CCNA_02816 | hypothetical protein | 1,154049 | 2,24E-05 | 0,0003354 |
| 167 | CCNA_03711 | ribosome-associated factor Y | 1,149614 | 3,29E-07 | 7,56E-06 |
| 168 | CCNA_03882 | phage gp6-like head-tail connector protein | 1,14729 | 0,005454 | 0,0396887 |
| 169 | CCNA_01971 | peptidyl-prolyl cis-trans isomerase | 1,145348 | 1,49E-06 | 2,91E-05 |
| 170 | CCNA_00096 | L-asparaginase | 1,1444 | 1,62E-06 | 3,13E-05 |
| 171 | CCNA_03339 | TolA protein | 1,143314 | 6,38E-06 | 0,0001071 |
| 172 | CCNA_01569 | transcriptional regulatory protein | 1,141523 | 4,87E-05 | 0,0006419 |
| 173 | CCNA_00307 | phospholipid-lipopolysaccharide ABC transporter | 1,137902 | 7,09E-05 | 0,0008936 |
| 174 | CCNA_03318 | cytosolic protein | 1,136987 | 0,0013032 | 0,0115629 |
| 175 | CCNA_R0048 | tRNA Leu | 1,128249 | 0,0001126 | 0,0013324 |
| 176 | CCNA_03209 | acetyl-coenzyme A synthetase | 1,126587 | 0,0018792 | 0,0158735 |
| 177 | CCNA_01503 | hypothetical protein | 1,122059 | 0,0010962 | 0,0100576 |
| 178 | CCNA_00294 | phosphate transport ATP-binding protein pstB | 1,121301 | 5,37E-07 | 1,17E-05 |
| 179 | CCNA_03782 | heme exporter protein A | 1,115029 | 0,0002022 | 0,0022479 |
| 180 | CCNA_01584 | multimodular transpeptidase-transglycosylase Pbp1a (PBP 1A) | 1,112263 | 8,32E-07 | 1,70E-05 |
| 181 | CCNA_01267 | hypothetical protein | 1,107908 | 5,96E-05 | 0,0007699 |
| 182 | CCNA_01268 | cytochrome c | 1,102669 | 2,23E-07 | 5,30E-06 |
| 183 | CCNA_00486 | TonB-dependent receptor | 1,101377 | 4,21E-05 | 0,0005655 |
| 184 | CCNA_03783 | heme exporter protein B | 1,100177 | 0,0004234 | 0,0043369 |

| | | | | | |
|---|---|---|---|---|---|
| 185 | CCNA_02129 | acyl-CoA dehydrogenase, short-chain specific | 1,099674 | 1,40E-06 | 2,75E-05 |
| 186 | CCNA_03487 | hypothetical protein | 1,091469 | 1,11E-05 | 0,000180 2 |
| 187 | CCNA_03357 | hypothetical protein | 1,087396 | 0,001023 6 | 0,009477 5 |
| 188 | CCNA_03175 | cytosolic protein | 1,083812 | 0,000113 4 | 0,001338 |
| 189 | CCNA_03786 | heme chaperone heme-lyase | 1,083588 | 1,96E-06 | 3,70E-05 |
| 190 | CCNA_R0180 | small non-coding RNA | 1,08319 | 5,50E-05 | 0,000715 9 |
| 191 | CCNA_00359 | SH3 domain-containing cell surface protein | 1,079742 | 0,000304 5 | 0,003241 8 |
| 192 | CCNA_02653 | sensory transduction protein kinase | 1,079419 | 9,21E-05 | 0,001116 |
| 193 | CCNA_03864 | cytosolic protein | 1,077941 | 8,31E-06 | 0,000137 7 |
| 194 | CCNA_03621 | transcriptional regulator, AraC family | 1,076176 | 2,29E-05 | 0,000339 9 |
| 195 | CCNA_01991 | hypothetical protein | 1,074656 | 1,90E-06 | 3,61E-05 |
| 196 | CCNA_02064 | UDP-3-O-(3-hydroxymyristoyl) N-acetylglucosamine deacetylase LpxC | 1,069027 | 2,84E-06 | 5,18E-05 |
| 197 | CCNA_03020 | hypothetical protein | 1,054913 | 2,58E-06 | 4,81E-05 |
| 198 | CCNA_01663 | potassium-transporting ATPase A chain | 1,03838 | 0,001118 6 | 0,010193 |
| 199 | CCNA_01732 | DNA repair protein RadA | 1,024789 | 2,41E-05 | 0,000351 7 |
| 200 | CCNA_02075 | peptidoglycan-specific endopeptidase, M23 family DipM | 1,016395 | 2,94E-05 | 0,000417 2 |
| 201 | CCNA_01831 | tryptophan halogenase superfamily protein | 1,011794 | 0,004722 8 | 0,034869 3 |
| 202 | CCNA_00978 | transposase | 1,011037 | 0,007065 8 | 0,048537 2 |
| 203 | CCNA_02082 | Sec-independent protein translocase protein tatA | 0,996894 | 0,000225 2 | 0,002482 3 |
| 204 | CCNA_01082 | hypothetical protein | 0,991064 | 0,001114 3 | 0,010177 4 |
| 205 | CCNA_01709 | hypothetical protein | 0,985944 | 7,12E-05 | 0,000893 9 |
| 206 | CCNA_01442 | N-acetyl-gamma-glutamyl-phosphate reductase | 0,982854 | 0,000385 1 | 0,003965 1 |
| 207 | CCNA_00379 | thiol:disulfide interchange protein dsbA | 0,980262 | 0,000102 4 | 0,001232 5 |
| 208 | CCNA_02794 | asparagine synthetase (glutamine-hydrolyzing) | 0,979714 | 0,000273 1 | 0,002946 1 |
| 209 | CCNA_03671 | predicted phosphohydrolase, Icc family | 0,977623 | 0,002628 | 0,021351 3 |
| 210 | CCNA_01095 | phenylalanyl-tRNA synthetase subunit beta | 0,971444 | 0,002215 3 | 0,018329 2 |
| 211 | CCNA_03791 | transcriptional regulator, MarR family | 0,965848 | 4,00E-05 | 0,000539 4 |
| 212 | CCNA_03091 | hypothetical protein NstA | 0,965182 | 0,000296 4 | 0,003172 8 |

| | | | | | |
|---|---|---|---|---|---|
| 213 | CCNA_01955 | zinc metalloprotease | 0,963472 | 0,0001977 | 0,0022162 |
| 214 | **CCNA_01638** | beta-lactamase family protein | 0,96309 | 6,39E-05 | 0,0008186 |
| 215 | **CCNA_03356** | cytosolic protein "zapA binds to FtsZ" | 0,958486 | 0,0007594 | 0,0072468 |
| 216 | CCNA_00170 | TonB-dependent receptor | 0,956425 | 5,86E-05 | 0,0007605 |
| 217 | CCNA_00196 | 3-isopropylmalate dehydratase large subunit | 0,954461 | 0,0004716 | 0,0047585 |
| 218 | CCNA_02378 | hypothetical protein | 0,953957 | 0,0032419 | 0,0254201 |
| 219 | **CCNA_01612** | rod shape-determining protein mreB | 0,948152 | 2,52E-05 | 0,0003646 |
| 220 | CCNA_00138 | TonB-dependent receptor | 0,946772 | 2,62E-05 | 0,0003774 |
| 221 | CCNA_02955 | putative permease | 0,943332 | 0,0016803 | 0,0143432 |
| 222 | CCNA_03459 | methyl-accepting chemotaxis protein | 0,940613 | 7,14E-05 | 0,0008939 |
| 223 | CCNA_01342 | ATPase, AAA family | 0,939596 | 2,35E-05 | 0,0003472 |
| 224 | CCNA_01504 | ATP-dependent DNA helicase recG | 0,922947 | 2,39E-05 | 0,0003515 |
| 225 | CCNA_00906 | hypothetical protein | 0,922502 | 0,0001983 | 0,0022164 |
| 226 | **CCNA_02008** | prolyl-tRNA synthetase | 0,916529 | 2,48E-05 | 0,0003604 |
| 227 | CCNA_02942 | hypothetical protein | 0,907706 | 5,40E-05 | 0,0007043 |
| 228 | CCNA_00120 | acetyltransferase | 0,902977 | 0,0005857 | 0,0057935 |
| 229 | **CCNA_03838** | gluconate 2-dehydrogenase/glyoxylate reductase/hydroxypyruvate reductase | 0,902758 | 0,0031848 | 0,0250206 |
| 230 | CCNA_03780 | hypothetical protein | 0,8922 | 0,0006172 | 0,0060171 |
| 231 | CCNA_01028 | cytosol aminopeptidase | 0,88544 | 0,0052442 | 0,0383692 |
| 232 | **CCNA_03716** | hypothetical protein | 0,884475 | 0,000217 | 0,0023983 |
| 233 | CCNA_01448 | fructose-1,6-bisphosphatase | 0,884159 | 0,0003022 | 0,0032265 |
| 234 | CCNA_02325 | ornithine carbamoyltransferase | 0,881094 | 0,0021261 | 0,0176278 |
| 235 | CCNA_03488 | hypothetical protein | 0,879008 | 0,0005483 | 0,0054644 |
| 236 | CCNA_01306 | LSU ribosomal protein L3P | 0,874311 | 0,002926 | 0,0234437 |
| 237 | CCNA_01582 | hypothetical protein | 0,872429 | 0,001489 | 0,0129842 |
| 238 | CCNA_00295 | phosphate transport system protein phoU | 0,865636 | 0,0001425 | 0,0016522 |
| 239 | CCNA_02317 | hypothetical protein | 0,860708 | 0,0011712 | 0,0105901 |

| | | | | | |
|---|---|---|---|---|---|
| 240 | CCNA_01664 | potassium-transporting ATPase B chain | 0,858797 | 0,0070552 | 0,0485372 |
| 241 | CCNA_03305 | SSU ribosomal protein S7P | 0,855205 | 0,0001433 | 0,0016564 |
| 242 | CCNA_02326 | acetylornithine aminotransferase/succinyldiaminopimelate aminotransferase | 0,853277 | 0,0018229 | 0,0154299 |
| 243 | **CCNA_03837** | hypothetical protein | 0,847624 | 0,0012026 | 0,0107415 |
| 244 | CCNA_01416 | 3-hydroxyisobutyrate dehydrogenase | 0,8471 | 0,0071303 | 0,048732 |
| 245 | CCNA_R0162 | small non-coding RNA | 0,838332 | 0,0040137 | 0,0305251 |
| 246 | CCNA_01891 | enoyl-CoA hydratase/carnithine racemase | 0,835318 | 0,0016618 | 0,0142355 |
| 247 | **CCNA_00419** | fasciclin domain cell surface protein | 0,834796 | 0,0001816 | 0,0020555 |
| 248 | CCNA_01026 | hypothetical protein | 0,828752 | 0,0025994 | 0,0212043 |
| 249 | CCNA_03174 | permease | 0,827187 | 0,0011038 | 0,0101041 |
| 250 | **CCNA_00517** | peptidyl-tRNA hydrolase | 0,82139 | 0,0035583 | 0,0274224 |
| 251 | **CCNA_01614** | hypothetical protein mreD | 0,800707 | 0,0026613 | 0,0215783 |
| 252 | CCNA_00420 | hypothetical protein | 0,800279 | 0,0015977 | 0,0138425 |
| 253 | CCNA_03304 | protein translation elongation factor G (EF-G) | 0,791329 | 0,0044151 | 0,0332037 |
| 254 | CCNA_00360 | putative hydrolase | 0,791311 | 0,006858 | 0,0475943 |
| 255 | CCNA_01980 | methylisocitrate lyase | 0,788602 | 0,0005956 | 0,0058778 |
| 256 | CCNA_00151 | ribonuclease PH | 0,786424 | 0,000537 | 0,0053743 |
| 257 | **CCNA_03727** | small-conductance mechanosensitive channel mcsS | 0,785585 | 0,0004668 | 0,0047344 |
| 258 | CCNA_00085 | dienelactone hydrolase-related protein | 0,775303 | 0,0019485 | 0,0163564 |
| 259 | CCNA_02480 | benzaldehyde dehydrogenase (NAD+) | 0,771528 | 0,0067963 | 0,0473065 |
| 260 | CCNA_03351 | phosphoribosylamidoimidazole-succinocarboxamide synthase | 0,768125 | 0,0044534 | 0,0333556 |
| 261 | **CCNA_00519** | hypothetical protein | 0,766235 | 0,0007975 | 0,0075746 |
| 262 | CCNA_01863 | outer membrane efflux protein | 0,76103 | 0,0066595 | 0,0466973 |
| 263 | **CCNA_01438** | heme O monooxygenase | 0,757038 | 0,0013238 | 0,0116947 |
| 264 | CCNA_00794 | hypothetical protein | 0,756107 | 0,0036754 | 0,0281111 |
| 265 | CCNA_01330 | DNA-directed RNA polymerase subunit alpha | 0,751066 | 0,0021012 | 0,0175287 |
| 266 | **CCNA_00377** | chromosome partition protein smc | 0,745145 | 0,0009538 | 0,0089334 |

| | | | | | |
|---|---|---|---|---|---|
| 267 | CCNA_02923 | TonB-dependent receptor | 0,740021 | 0,004107 9 | 0,031182 7 |
| 268 | CCNA_03865 | leucyl-tRNA synthetase | 0,739799 | 0,001779 7 | 0,015127 2 |
| 269 | CCNA_02370 | TonB-dependent maltose outer membrane transporter malA | 0,725134 | 0,001512 8 | 0,013162 9 |
| 270 | CCNA_01252 | soluble lytic murein transglycosylase SdpA | 0,72267 | 0,002625 8 | 0,021351 3 |
| 271 | CCNA_03307 | hypothetical protein | 0,721804 | 0,000962 1 | 0,008990 2 |
| 272 | CCNA_R0039 | tRNA Arg | 0,719034 | 0,003488 1 | 0,027035 8 |
| 273 | **CCNA_03195** | RNA polymerase sigma-32 factor | 0,715546 | 0,004432 1 | 0,033269 6 |
| 274 | CCNA_03735 | transketolase | 0,712932 | 0,003036 | 0,024038 7 |
| 275 | CCNA_00306 | hypothetical protein | 0,710158 | 0,006993 7 | 0,048205 3 |
| 276 | CCNA_01546 | DEAD-box RNA helicase-like protein | 0,709828 | 0,005851 3 | 0,042125 |
| 277 | CCNA_01793 | hypothetical protein | 0,705985 | 0,001178 5 | 0,010619 6 |
| 278 | CCNA_00839 | glucan 1,4-beta-glucosidase | 0,702798 | 0,007212 | 0,049206 8 |
| 279 | CCNA_03752 | UTP-glucose-1-phosphate uridylyltransferase | 0,697334 | 0,006274 7 | 0,044774 9 |
| 280 | **CCNA_00585** | hypothetical protein | 0,696581 | 0,002363 | 0,019472 2 |
| 281 | CCNA_00037 | esterase lipase family protein | 0,696386 | 0,002981 3 | 0,023698 5 |
| 282 | CCNA_02583 | phosphoribosylformylglycinamidine synthase I | 0,690972 | 0,003091 8 | 0,024432 6 |
| 283 | **CCNA_02007** | lipoprotein releasing system transmembrane protein lolE | 0,685244 | 0,001719 7 | 0,014647 9 |
| 284 | CCNA_00778 | GTP-binding protein TypA/BipA | 0,683274 | 0,006235 8 | 0,044576 1 |
| 285 | CCNA_01637 | topoisomerase IV subunit A | 0,672635 | 0,002572 7 | 0,021028 5 |
| 286 | **CCNA_03682** | fumarylpyruvate hydrolase | 0,669784 | 0,001811 9 | 0,015369 |
| 287 | CCNA_01956 | outer membrane protein | 0,66209 | 0,002949 7 | 0,023586 3 |
| 288 | CCNA_02232 | TonB-dependent receptor | 0,659847 | 0,005912 1 | 0,042487 |
| 289 | **CCNA_03683** | maleylpyruvate isomerase | 0,658792 | 0,004460 1 | 0,033355 6 |
| 290 | CCNA_00495 | cytosolic protein | 0,658202 | 0,003450 8 | 0,026798 4 |
| 291 | CCNA_01990 | UDP-3-O-(3-hydroxymyristoyl) glucosamine N-acyltransferase | 0,656367 | 0,006782 8 | 0,047306 5 |
| 292 | **CCNA_01060** | type I protein secretion ATP-binding protein RsaD | 0,638711 | 0,003365 | 0,026182 4 |
| 293 | CCNA_03879 | uroporphyrinogen decarboxylase | 0,63689 | 0,004168 1 | 0,031580 7 |

| | | | | | |
|---|---|---|---|---|---|
| 294 | CCNA_01851 | quinol cytochrome oxidase polypeptide II | 0,627876 | 0,0064339 | 0,0455141 |
| 295 | CCNA_02406 | membrane lipoprotein | 0,624075 | 0,0059558 | 0,0427258 |
| 296 | CCNA_00483 | Zn-dependent hydrolase, glyoxalase II family | 0,620069 | 0,0066746 | 0,0467224 |
| 297 | CCNA_03306 | SSU ribosomal protein S12P | 0,614043 | 0,0044959 | 0,0334999 |
| 298 | CCNA_01237 | hypothetical protein | 0,609086 | 0,0068048 | 0,0473065 |
| 299 | CCNA_01650 | copper/zinc superoxide dismutase | 0,605626 | 0,0056222 | 0,040693 |
| 300 | CCNA_01223 | acyl-CoA synthetase | 0,600935 | 0,0067325 | 0,0470458 |
| 301 | CCNA_00029 | lysine exporter protein | 0,596043 | 0,0070988 | 0,0485986 |

Genes highlighted in green were identified in the ChIP-seq regulon in this work, and those in yellow the RNA-seq regulon presented by Stein *et al.* (2021).

#### Table S6. Genes identified by RNA-seq up-regulated in the Δ*chvI* mutant upon osmotic stress with 6% sucrose.

| Top hit | Gene ID | Description | log2 (FC Δ*chvI*/WT ) | P-value | P-adj |
|---|---|---|---|---|---|
| 1 | CCNA_01202 | membrane alanine aminopeptidase | -1,933379 | 2,36E-18 | 1,51E-16 |
| 2 | **CCNA_00438** | hypothetical protein | -1,913719 | 5,69E-06 | 9,67E-05 |
| 3 | **CCNA_01303** | hypothetical protein | -1,882275 | 1,16E-11 | 4,64E-10 |
| 4 | **CCNA_00974** | OAR protein precursor | -1,789948 | 1,36E-08 | 3,85E-07 |
| 5 | CCNA_R0061 | RNase P RNA | -1,754613 | 5,70E-07 | 1,24E-05 |
| 6 | CCNA_01523 | acetyltransferase flmH | -1,690376 | 1,57E-10 | 5,81E-09 |
| 7 | **CCNA_03108** | ChvT TonB-dependent outer membrane receptor | -1,689341 | 8,08E-07 | 1,66E-05 |
| 8 | CCNA_01121 | hypothetical protein | -1,678219 | 4,09E-11 | 1,56E-09 |
| 9 | **CCNA_02200** | cytochrome c-family protein | -1,647808 | 8,55E-08 | 2,19E-06 |
| 10 | **CCNA_00444** | chemotaxis protein methyltransferase | -1,642055 | 1,23E-10 | 4,60E-09 |
| 11 | CCNA_01186 | hypothetical protein | -1,62859 | 0,00026 | 0,0028554 |
| 12 | CCNA_03043 | type IV pilin protein pilA | -1,601108 | 0,0001 | 0,0012221 |
| 13 | CCNA_03395 | two-component receiver domain protein | -1,579211 | 1,27E-05 | 0,0002011 |
| 14 | **CCNA_03325** | hypothetical protein | -1,550968 | 7,06E-07 | 1,47E-05 |
| 15 | CCNA_01476 | CRP-family transcription regulator ftrB | -1,530608 | 2,61E-10 | 9,17E-09 |
| 16 | **CCNA_01269** | GcrB protein | -1,52671 | 2,10E-09 | 6,78E-08 |
| 17 | CCNA_02515 | cytosolic protein | -1,524018 | 6,02E-07 | 1,30E-05 |
| 18 | **CCNA_00417** | hypothetical protein | -1,522626 | 3,87E-09 | 1,20E-07 |

| | | | | | |
|---|---|---|---|---|---|
| 19 | CCNA_00948 | hypothetical protein =SciP | -1,515411 | 3,61E-12 | 1,51E-10 |
| 20 | CCNA_03607 | ribonucleoside-diphosphate reductase subunit alpha | -1,505469 | 3,53E-09 | 1,11E-07 |
| 21 | CCNA_00709 | hypothetical protein | -1,500115 | 2,67E-05 | 0,0003829 |
| 22 | **CCNA_00446** | chemotaxis receiver domain protein cheYII | -1,493908 | 1,22E-09 | 4,15E-08 |
| 23 | CCNA_03896 | conserved hypothetical protein | -1,491758 | 1,39E-09 | 4,63E-08 |
| 24 | CCNA_03136 | flagellar export protein FliJ | -1,489151 | 3,63E-07 | 8,31E-06 |
| 25 | CCNA_01475 | OmpW family outer membrane protein | -1,484898 | 2,35E-08 | 6,38E-07 |
| 26 | CCNA_01901 | lipoprotein | -1,464359 | 1,97E-10 | 7,26E-09 |
| 27 | **CCNA_00439** | methyl-accepting chemotaxis protein McpA | -1,453836 | 3,78E-08 | 9,99E-07 |
| 28 | CCNA_02931 | flgE-related flagellar hook protein | -1,447194 | 4,01E-06 | 7,03E-05 |
| 29 | CCNA_03700 | large-conductance mechanosensitive channel | -1,445665 | 1,59E-06 | 3,09E-05 |
| 30 | **CCNA_03313** | hypothetical protein | -1,445108 | 3,80E-06 | 6,71E-05 |
| 31 | CCNA_03944 | hypothetical protein | -1,441941 | 0,00032 | 0,0033646 |
| 32 | CCNA_01270 | hypothetical protein | -1,438841 | 1,91E-08 | 5,30E-07 |
| 33 | CCNA_03355 | acylamino-acid-releasing enzyme | -1,43795 | 3,74E-11 | 1,44E-09 |
| 34 | CCNA_00350 | hypothetical protein | -1,435351 | 3,40E-06 | 6,14E-05 |
| 35 | **CCNA_00440** | virus protein | -1,423983 | 3,91E-07 | 8,90E-06 |
| 36 | CCNA_02142 | flagellar basal-body rod protein flgF | -1,420865 | 3,23E-07 | 7,47E-06 |
| 37 | CCNA_01526 | flagellar biosynthesis regulatory protein flaF | -1,414115 | 2,00E-05 | 0,0003071 |
| 38 | CCNA_01201 | hypothetical protein | -1,412886 | 1,01E-08 | 2,90E-07 |
| 39 | CCNA_02604 | host cell attachment protein | -1,411122 | 2,66E-06 | 4,94E-05 |
| 40 | CCNA_02257 | fliN family protein | -1,390661 | 0,0003 | 0,003169 |
| 41 | **CCNA_00441** | chemotaxis receiver domain protein cheYI | -1,368874 | 2,95E-09 | 9,41E-08 |
| 42 | **CCNA_00628** | chemotaxis protein cheY | -1,36271 | 1,37E-07 | 3,37E-06 |
| 43 | CCNA_01278 | histidine phosphotransferase domain protein | -1,361569 | 0,00013 | 0,0014602 |
| 44 | CCNA_R0176 | small non-coding RNA | -1,360767 | 0,00618 | 0,044279 |
| 45 | CCNA_01926 | two-component response regulator DgcB | -1,351856 | 1,43E-09 | 4,74E-08 |
| 46 | CCNA_02754 | parathion hydrolase | -1,350412 | 7,05E-10 | 2,44E-08 |
| 47 | CCNA_00835 | flagellin | -1,348627 | 4,28E-07 | 9,56E-06 |
| 48 | CCNA_01157 | protoporphyrinogen oxidase | -1,348367 | 7,48E-10 | 2,56E-08 |
| 49 | CCNA_01528 | flagellin fljK | -1,345288 | 1,07E-06 | 2,14E-05 |
| 50 | CCNA_00447 | chemotaxis protein cheD | -1,33167 | 2,67E-07 | 6,27E-06 |

| | | | | | |
|---|---|---|---|---|---|
| 51 | CCNA_02712 | holdfast attachment protein hfaB | -1,323383 | 3,98E-06 | 7,00E-05 |
| 52 | **CCNA_00166** | HvyA | -1,318296 | 8,98E-08 | 2,28E-06 |
| 53 | CCNA_03246 | hypothetical protein | -1,312915 | 1,11E-06 | 2,19E-05 |
| 54 | CCNA_00943 | flagellar hook-associated protein FlaN | -1,312769 | 6,38E-09 | 1,88E-07 |
| 55 | **CCNA_00236** | hypothetical protein | -1,307781 | 7,38E-06 | 0,0001229 |
| 56 | CCNA_00942 | flagellar hook-associated protein FlgL | -1,304491 | 8,56E-08 | 2,19E-06 |
| 57 | CCNA_00953 | flagellar motor switch protein FliN | -1,304384 | 8,33E-08 | 2,16E-06 |
| 58 | CCNA_00437 | methyl-accepting chemotaxis protein | -1,299406 | 7,59E-07 | 1,57E-05 |
| 59 | CCNA_00443 | chemotaxis protein cheW | -1,295113 | 2,09E-05 | 0,0003179 |
| 60 | CCNA_02667 | flagellar basal-body protein flbY | -1,291132 | 6,53E-07 | 1,38E-05 |
| 61 | CCNA_02711 | holdfast attachment protein hfaA | -1,288624 | 1,25E-05 | 0,0001986 |
| 62 | **CCNA_00442** | chemotaxis histidine kinase protein cheAI | -1,282375 | 1,67E-08 | 4,65E-07 |
| 63 | CCNA_02922 | hypothetical protein | -1,274303 | 0,00013 | 0,0015455 |
| 64 | CCNA_00348 | probable chemoreceptor Y4FA | -1,27202 | 6,79E-09 | 1,99E-07 |
| 65 | CCNA_02845 | two-component response regulator | -1,26644 | 2,01E-08 | 5,53E-07 |
| 66 | CCNA_02199 | methyltransferase | -1,263917 | 2,97E-07 | 6,95E-06 |
| 67 | CCNA_02143 | flagellar basal-body rod protein flgG | -1,263858 | 1,06E-07 | 2,67E-06 |
| 68 | **CCNA_01034** | TonB-dependent outer membrane receptor | -1,258003 | 7,64E-09 | 2,22E-07 |
| 69 | CCNA_00542 | hypothetical protein | -1,257973 | 0,00039 | 0,0040559 |
| 70 | CCNA_00382 | adenine-specific methyltransferase ccrM | -1,250604 | 8,30E-09 | 2,40E-07 |
| 71 | CCNA_02547 | response regulator receiver protein divK | -1,246185 | 2,98E-05 | 0,0004207 |
| 72 | CCNA_03999 | hypothetical protein | -1,241266 | 2,38E-06 | 4,45E-05 |
| 73 | CCNA_03539 | hypothetical protein | -1,23881 | 1,32E-07 | 3,28E-06 |
| 74 | CCNA_01477 | oxygen-independent coproporphyrinogen-III oxidase hemN | -1,2367 | 1,08E-07 | 2,71E-06 |
| 75 | CCNA_01365 | aspartyl protease perP | -1,235193 | 2,98E-05 | 0,0004207 |
| 76 | CCNA_00538 | methyl-accepting chemotaxis protein | -1,234609 | 3,06E-08 | 8,26E-07 |
| 77 | CCNA_00426 | very-short-patch-repair endonuclease | -1,233175 | 0,00062 | 0,0060171 |
| 78 | CCNA_02666 | chemotactic signal-response protein cheL | -1,208915 | 1,21E-05 | 0,0001936 |
| 79 | CCNA_01529 | hypothetical protein | -1,204067 | 1,14E-05 | 0,0001833 |
| 80 | CCNA_00803 | chemotaxis protein cheW | -1,189911 | 6,28E-07 | 1,34E-05 |
| 81 | CCNA_00965 | chaperone protein DnaJ | -1,188986 | 3,07E-05 | 0,0004289 |
| 82 | CCNA_00354 | hypothetical protein | -1,185738 | 0,00043 | 0,0044134 |
| 83 | CCNA_02105 | hypothetical protein | -1,181311 | 0,00095 | 0,0089075 |

| | | | | | |
|---|---|---|---|---|---|
| 84 | CCNA_02145 | flagellar L-ring protein flgH | -1,181267 | 9,66E-07 | 1,95E-05 |
| 85 | CCNA_02844 | antitoxin protein parD-3 | -1,180372 | 2,78E-06 | 5,08E-05 |
| 86 | CCNA_00027 | 2OG-Fe(II) oxygenase | -1,175011 | 1,01E-05 | 0,0001645 |
| 87 | CCNA_02665 | flagellar P-ring protein flgI | -1,165739 | 1,60E-06 | 3,10E-05 |
| 88 | CCNA_02363 | hypothetical protein | -1,160395 | 0,00019 | 0,0021641 |
| 89 | CCNA_01644 | chemotaxis motB protein | -1,158135 | 6,49E-07 | 1,37E-05 |
| 90 | CCNA_02141 | flagellar fliL protein | -1,157392 | 2,22E-05 | 0,0003339 |
| 91 | CCNA_00790 | hypoxia negative feedback regulator FixT | -1,157196 | 0,00047 | 0,0047344 |
| 92 | CCNA_02408 | hypothetical protein | -1,155134 | 0,00032 | 0,0034072 |
| 93 | CCNA_03932 | hypothetical protein | -1,155059 | 5,03E-07 | 1,11E-05 |
| 94 | CCNA_03993 | hypothetical protein | -1,15475 | 0,00389 | 0,0296058 |
| 95 | CCNA_00081 | hypothetical protein | -1,153832 | 2,19E-05 | 0,0003313 |
| 96 | CCNA_01032 | RNA polymerase ECF-type sigma factor | -1,151891 | 0,00032 | 0,003351 |
| 97 | CCNA_01119 | hypothetical protein | -1,150537 | 0,00033 | 0,0034473 |
| 98 | CCNA_02513 | holdfast synthesis gene hfsA, frameshifted variant | -1,145246 | 0,00024 | 0,002637 |
| 99 | CCNA_03762 | hypothetical protein | -1,14113 | 0,0006 | 0,0059093 |
| 100 | **CCNA_00471** | GDP-L-fucose synthase | -1,140668 | 3,43E-06 | 6,14E-05 |
| 101 | CCNA_00094 | probable UDP-N-acetyl-D-mannosaminuronic acid transferase | -1,138585 | 6,60E-07 | 1,38E-05 |
| 102 | CCNA_03691 | KidO | -1,137173 | 6,14E-05 | 0,0007892 |
| 103 | CCNA_00729 | hypothetical protein | -1,135676 | 1,83E-05 | 0,0002844 |
| 104 | **CCNA_00445** | receiver domain-glutamate methylesterase cheBI | -1,1356 | 1,27E-07 | 3,17E-06 |
| 105 | CCNA_03923 | hypothetical protein | -1,134216 | 0,00059 | 0,0057935 |
| 106 | CCNA_02139 | polar flagellum positioning protein pflI | -1,131809 | 5,59E-06 | 9,56E-05 |
| 107 | CCNA_R0117 | small non-coding RNA | -1,127058 | 3,04E-05 | 0,0004263 |
| 108 | **CCNA_00472** | GDP-mannose 4,6 dehydratase | -1,123613 | 4,86E-07 | 1,08E-05 |
| 109 | **CCNA_00416** | hypothetical protein | -1,12286 | 2,69E-06 | 4,96E-05 |
| 110 | **CCNA_00448** | cheU protein | -1,119834 | 4,31E-06 | 7,42E-05 |
| 111 | CCNA_00946 | Basal-body rod modification protein FlgD | -1,115472 | 1,80E-06 | 3,44E-05 |
| 112 | CCNA_00093 | hypothetical protein | -1,113679 | 3,14E-05 | 0,0004364 |
| 113 | CCNA_02196 | hypothetical protein | -1,110725 | 7,34E-05 | 0,0009168 |
| 114 | CCNA_00065 | hypothetical protein | -1,108239 | 0,00012 | 0,0013551 |
| 115 | CCNA_03037 | pilus assembly ATPase CpaF | -1,106773 | 3,23E-05 | 0,0004458 |
| 116 | CCNA_02534 | hypothetical protein | -1,10674 | 0,00298 | 0,0236972 |

| | | | | | |
|---|---|---|---|---|---|
| 117 | CCNA_02364 | methyl-accepting chemotaxis protein | -1,102917 | 7,75E-07 | 1,60E-05 |
| 118 | CCNA_03120 | chemotaxis protein cheW | -1,099605 | 2,02E-05 | 0,000308 |
| 119 | CCNA_01524 | FlbA protein | -1,099099 | 4,29E-06 | 7,42E-05 |
| 120 | CCNA_01532 | regulatory protein flaY | -1,097971 | 1,56E-05 | 0,0002447 |
| 121 | CCNA_01685 | hemimethylated DNA-binding protein yccV | -1,082185 | 0,00093 | 0,0088028 |
| 122 | CCNA_00787 | chemotaxis motA protein | -1,078746 | 4,68E-05 | 0,0006215 |
| 123 | CCNA_02546 | GGDEF/response regulator protein pleD | -1,073107 | 0,00071 | 0,0068204 |
| 124 | CCNA_03915 | hypothetical protein | -1,071251 | 0,00236 | 0,0194722 |
| 125 | CCNA_02722 | hypothetical protein | -1,068107 | 2,29E-05 | 0,0003399 |
| 126 | CCNA_02597 | hypothetical protein | -1,067281 | 8,60E-06 | 0,000142 |
| 127 | CCNA_02621 | CAAX amino terminal protease family | -1,063135 | 9,89E-06 | 0,0001621 |
| 128 | CCNA_01108 | nucleoside-diphosphate-sugar epimerase | -1,058775 | 6,17E-06 | 0,0001041 |
| 129 | CCNA_03039 | pilus assembly protein CpaD | -1,058212 | 3,45E-05 | 0,0004734 |
| 130 | CCNA_02720 | hypothetical protein | -1,057207 | 0,00036 | 0,0037039 |
| 131 | CCNA_02517 | hypothetical protein | -1,056371 | 4,21E-06 | 7,31E-05 |
| 132 | CCNA_00629 | methyl-accepting chemotaxis protein | -1,054025 | 2,12E-05 | 0,0003215 |
| 133 | CCNA_02622 | M61 glycyl aminopeptidase | -1,0535 | 2,70E-06 | 4,96E-05 |
| 134 | CCNA_01424 | hypothetical protein | -1,052211 | 3,34E-06 | 6,06E-05 |
| 135 | CCNA_01163 | ice nucleation protein | -1,048322 | 5,60E-06 | 9,56E-05 |
| 136 | CCNA_03042 | pilus assembly prepilin peptidase CpaA | -1,03898 | 0,00063 | 0,00607 |
| 137 | CCNA_02976 | hypothetical protein | -1,037237 | 6,54E-05 | 0,0008344 |
| 138 | CCNA_02540 | N-acyl-L-amino acid amidohydrolase | -1,036741 | 1,18E-06 | 2,33E-05 |
| 139 | CCNA_03763 | deacetylase | -1,036727 | 7,12E-06 | 0,000119 |
| 140 | CCNA_00834 | flagellin | -1,034313 | 5,22E-05 | 0,0006838 |
| 141 | CCNA_02514 | polysaccharide secretin protein hfsD | -1,031577 | 0,00095 | 0,0089075 |
| 142 | CCNA_03265 | CBS pair-family sensor histidine kinase/receiver domain protein | -1,028312 | 0,00034 | 0,0035669 |
| 143 | CCNA_00982 | transcriptional regulator | -1,024448 | 0,00121 | 0,0107782 |
| 144 | CCNA_01466 | hypothetical protein | -1,024376 | 0,00693 | 0,0478721 |
| 145 | CCNA_03295 | PAS-family sensor histidine kinase | -1,021901 | 2,75E-05 | 0,0003915 |
| 146 | CCNA_00950 | flagellar M-ring protein FliF | -1,020532 | 2,18E-06 | 4,09E-05 |
| 147 | CCNA_03247 | methyl-accepting chemotaxis protein | -1,0199 | 1,12E-05 | 0,0001802 |
| 148 | CCNA_00592 | hypothetical protein | -1,016761 | 0,00053 | 0,005313 |
| 149 | CCNA_03940 | hypothetical protein | -1,014706 | 0,0029 | 0,0233382 |

| | | | | | |
|---|---|---|---|---|---|
| 150 | CCNA_02680 | hypothetical protein | -1,014085 | 5,82E-06 | 9,86E-05 |
| 151 | CCNA_03062 | cell wall hydrolase family protein | -1,006635 | 1,80E-05 | 0,0002808 |
| 152 | CCNA_01952 | N-acetylmuramoyl-L-alanine amidase AmiC | -1,004564 | 0,00043 | 0,0044134 |
| 153 | CCNA_01258 | permease | -1,000021 | 0,00025 | 0,002733 |
| 154 | CCNA_02831 | hypothetical protein | -0,999066 | 2,40E-05 | 0,0003515 |
| 155 | CCNA_02713 | holdfast attachment protein hfaD | -0,998279 | 7,75E-05 | 0,0009527 |
| 156 | CCNA_03660 | Usg protein | -0,995798 | 0,00484 | 0,0356356 |
| 157 | CCNA_03202 | hypothetical protein | -0,993089 | 0,00022 | 0,0023983 |
| 158 | CCNA_02359 | hypothetical protein | -0,991779 | 6,14E-05 | 0,0007892 |
| 159 | CCNA_R0188 | small non-coding RNA | -0,981879 | 0,00643 | 0,0455141 |
| 160 | CCNA_00454 | transcriptional regulator, GntR family | -0,981577 | 3,24E-05 | 0,0004458 |
| 161 | CCNA_03585 | chemotaxis receiver domain protein cheYIV | -0,98132 | 0,00099 | 0,0092568 |
| 162 | CCNA_00951 | flagellar motor switch protein FliG | -0,978554 | 2,49E-05 | 0,0003609 |
| 163 | CCNA_00224 | hypothetical protein | -0,976234 | 4,58E-05 | 0,0006129 |
| 164 | CCNA_01122 | hypothetical protein | -0,975603 | 0,00066 | 0,0063429 |
| 165 | CCNA_03036 | TadB-related pilus assembly protein | -0,975251 | 0,00117 | 0,0105901 |
| 166 | CCNA_03198 | two-component response regulator | -0,973815 | 0,00166 | 0,0142355 |
| 167 | CCNA_02411 | putative lytic transglycosylase pleA | -0,971665 | 0,00115 | 0,0104287 |
| 168 | CCNA_03414 | NAD(P) transhydrogenase alpha subunit | -0,969938 | 8,84E-06 | 0,0001453 |
| 169 | CCNA_00945 | chemotaxis protein MotD | -0,967557 | 0,0001 | 0,0012326 |
| 170 | CCNA_00302 | parathion hydrolase | -0,967207 | 1,03E-05 | 0,0001675 |
| 171 | CCNA_03287 | transcriptional regulatory protein | -0,95314 | 0,00036 | 0,0037595 |
| 172 | CCNA_02625 | cell division protein ftsQ | -0,953117 | 0,00011 | 0,0013044 |
| 173 | CCNA_03839 | acylamino-acid-releasing enzyme | -0,951083 | 0,00012 | 0,0014597 |
| 174 | CCNA_01779 | hypothetical protein | -0,948325 | 0,0006 | 0,0059169 |
| 175 | CCNA_01257 | permease | -0,948023 | 0,00058 | 0,0057709 |
| 176 | CCNA_03424 | AAA-family response regulator tacA | -0,947344 | 1,89E-05 | 0,0002916 |
| 177 | CCNA_03249 | TonB-dependent receptor | -0,946453 | 0,00024 | 0,0026626 |
| 178 | CCNA_02643 | division specific D,D-transpeptidase/cell division protein ftsI | -0,945101 | 9,09E-05 | 0,0011042 |
| 179 | CCNA_00854 | metallo-beta-lactamase protein | -0,944003 | 1,44E-05 | 0,000227 |
| 180 | CCNA_01035 | gamma-glutamyltranspeptidase | -0,936811 | 3,91E-05 | 0,0005293 |
| 181 | CCNA_03137 | endo-1,4-beta-xylanase | -0,936141 | 0,00064 | 0,0062338 |
| 182 | CCNA_01525 | flagellar biosynthesis repressor flbT | -0,932605 | 0,0012 | 0,0107399 |

| | | | | | |
|---|---|---|---|---|---|
| 183 | CCNA_03135 | flagellum-specific ATP synthase fliL | -0,932485 | 4,98E-05 | 0,0006548 |
| 184 | CCNA_01004 | flagellar basal-body rod protein FlgB | -0,932255 | 0,00647 | 0,045581 |
| 185 | CCNA_02949 | hypothetical protein | -0,930419 | 0,00011 | 0,0012951 |
| 186 | CCNA_01185 | hypothetical protein | -0,928673 | 3,47E-05 | 0,0004736 |
| 187 | **CCNA_03089** | hypothetical protein | -0,927527 | 3,75E-05 | 0,0005094 |
| 188 | CCNA_02238 | hypothetical protein | -0,925002 | 0,00565 | 0,0407857 |
| 189 | CCNA_02360 | beta-D-Glcp beta-1,4-glucosyltransferase | -0,924491 | 0,00032 | 0,0033908 |
| 190 | CCNA_01465 | methyl-accepting chemotaxis protein | -0,921667 | 6,90E-05 | 0,000873 |
| 191 | CCNA_02644 | putative cell division protein | -0,919761 | 0,00293 | 0,0234437 |
| 192 | **CCNA_00627** | hypothetical protein | -0,919012 | 0,00439 | 0,0330854 |
| 193 | CCNA_03169 | hypothetical protein | -0,917501 | 0,00028 | 0,0029801 |
| 194 | CCNA_00234 | WecE-family cell wall biogenesis enzyme | -0,916928 | 9,08E-05 | 0,0011042 |
| 195 | CCNA_03686 | alpha2 macroglobulin domain-containing extracellular protein | -0,916167 | 3,02E-05 | 0,0004237 |
| 196 | CCNA_03130 | cell cycle response regulator ctrA | -0,915187 | 0,00101 | 0,0093773 |
| 197 | CCNA_03412 | NAD(P) transhydrogenase subunit beta | -0,913158 | 2,74E-05 | 0,0003915 |
| 198 | CCNA_03410 | peptidoglycan-specific endopeptidase, M23 family LdpE | -0,902956 | 0,00146 | 0,0127708 |
| 199 | CCNA_02719 | hypothetical protein | -0,901484 | 0,00161 | 0,0138884 |
| 200 | CCNA_03754 | glutathione S-transferase family protein FzlA | -0,89973 | 0,00133 | 0,0117216 |
| 201 | CCNA_01006 | flagellar hook-basal body complex protein FliE | -0,897561 | 0,00208 | 0,0173953 |
| 202 | CCNA_03413 | NAD(P) transhydrogenase alpha subunit | -0,89729 | 6,65E-05 | 0,0008463 |
| 203 | CCNA_02642 | UDP-N-acetylmuramoylalanyl-D-glutamate--2, 6-diaminopimelate ligase | -0,896947 | 7,52E-05 | 0,0009332 |
| 204 | CCNA_00875 | Flp/Fap pilin component protein | -0,896139 | 0,00367 | 0,0280942 |
| 205 | CCNA_00967 | transcriptional regulator, TetR family | -0,895939 | 0,00016 | 0,0018596 |
| 206 | CCNA_01248 | transcriptional regulator, TetR family | -0,894381 | 7,66E-05 | 0,0009472 |
| 207 | CCNA_00233 | UDP-N-acetylglucosamine 4,6-dehydratase FlaA1 | -0,892404 | 3,69E-05 | 0,000503 |
| 208 | CCNA_03327 | hypothetical protein | -0,889859 | 0,00447 | 0,0333915 |
| 209 | CCNA_02950 | hypothetical protein | -0,889715 | 0,00038 | 0,0039048 |
| 210 | **CCNA_03326** | two-component sensor histidine kinase | -0,883349 | 0,00364 | 0,0279109 |
| 211 | CCNA_00089 | MHYT/PAS-family GGDEF/EAL protein | -0,869832 | 0,00017 | 0,0019556 |
| 212 | CCNA_00591 | catalase | -0,869626 | 0,00034 | 0,0035669 |
| 213 | **CCNA_02910** | TonB-dependent receptor | -0,866451 | 0,00693 | 0,0478721 |
| 214 | CCNA_02449 | hypothetical protein | -0,861596 | 0,0007 | 0,0067258 |

| | | | | | |
|---|---|---|---|---|---|
| 215 | CCNA_03096 | TonB-dependent receptor | -0,859803 | 7,49E-05 | 0,0009324 |
| 216 | CCNA_00947 | flagellar hook protein FlgE | -0,852767 | 0,00641 | 0,0455141 |
| 217 | CCNA_01220 | serine palmitoyltransferase | -0,850873 | 0,00047 | 0,0047632 |
| 218 | CCNA_00944 | flagellar hook length determination protein | -0,849972 | 0,00017 | 0,0019573 |
| 219 | CCNA_03467 | hypothetical protein | -0,846304 | 0,00012 | 0,0013944 |
| 220 | CCNA_01181 | hypothetical protein | -0,845916 | 0,00165 | 0,014189 |
| 221 | CCNA_00853 | 3-isopropylmalate dehydrogenase | -0,841324 | 0,00047 | 0,0047456 |
| 222 | CCNA_03286 | transporter | -0,840942 | 0,00054 | 0,0053743 |
| 223 | CCNA_01671 | diguanylate receptor protein dgrA | -0,839315 | 0,00062 | 0,0060466 |
| 224 | CCNA_03040 | outer membrane pilus secretion channel cpaC | -0,838347 | 0,00016 | 0,0018647 |
| 225 | CCNA_03933 | hypothetical protein | -0,836738 | 0,00019 | 0,0021641 |
| 226 | CCNA_02645 | S-adenosyl-methyltransferase mraW | -0,836346 | 0,0035 | 0,0270763 |
| 227 | CCNA_00028 | TonB-dependent receptor | -0,831973 | 0,00061 | 0,0060152 |
| 228 | CCNA_01005 | flagellar basal-body rod protein flgC | -0,829987 | 0,0012 | 0,0107399 |
| 229 | CCNA_00952 | flagellar assembly protein FlbE/FliH | -0,823626 | 0,0019 | 0,0160238 |
| 230 | CCNA_00425 | hypothetical protein | -0,822988 | 0,00336 | 0,0261673 |
| 231 | CCNA_02639 | UDP-N-acetylmuramoylalanine--D-glutamate ligase | -0,820277 | 0,00363 | 0,0279109 |
| 232 | CCNA_03531 | carboxypeptidase S1 | -0,820215 | 0,0002 | 0,002232 |
| 233 | CCNA_03090 | acetyl-coenzyme A carboxylase carboxyl transferase subunit alpha | -0,819814 | 0,00015 | 0,0016859 |
| 234 | CCNA_02509 | glycosyltransferase hfsG | -0,81892 | 0,00283 | 0,0228679 |
| 235 | CCNA_02961 | N-acetylneuraminate synthase | -0,818244 | 0,00101 | 0,0093773 |
| 236 | CCNA_02138 | hypothetical protein | -0,814728 | 0,00131 | 0,0116178 |
| 237 | CCNA_02640 | phospho-N-acetylmuramoyl-pentapeptide- transferase | -0,814401 | 0,00037 | 0,0038455 |
| 238 | CCNA_02140 | flagellar motor switch protein fliM | -0,812975 | 0,00126 | 0,0112245 |
| 239 | CCNA_00080 | LexA-related transcriptional repressor | -0,809499 | 0,00026 | 0,0028048 |
| 240 | CCNA_02641 | UDP-N-acetylmuramoyl-tripeptide--D-alanyl-D- alanine ligase | -0,808499 | 0,00026 | 0,0028554 |
| 241 | CCNA_02163 | hypothetical protein | -0,807182 | 0,0012 | 0,0107399 |
| 242 | CCNA_01117 | hypothetical protein | -0,806545 | 0,00104 | 0,0095683 |
| 243 | CCNA_02221 | methionine synthase I metH | -0,799147 | 0,00138 | 0,0121438 |
| 244 | CCNA_03121 | hypothetical protein | -0,798173 | 0,00083 | 0,0078382 |
| 245 | CCNA_02361 | polysaccharide biosynthesis protein celD | -0,79652 | 0,00108 | 0,0099077 |
| 246 | CCNA_02172 | transporter | -0,796063 | 0,00154 | 0,0133396 |

| | | | | | |
|---|---|---|---|---|---|
| 247 | CCNA_03885 | Abi superfamily/CAAX amino terminal protease | -0,794258 | 0,00296 | 0,0236377 |
| 248 | CCNA_02409 | hybrid sensor histidine kinase/receiver protein | -0,791648 | 0,00109 | 0,0100576 |
| 249 | CCNA_02144 | flagella basal body P ring formation protein flgA | -0,79072 | 0,00641 | 0,0455141 |
| 250 | CCNA_00665 | GAF-family sensor histidine kinase | -0,785735 | 0,00664 | 0,0466355 |
| 251 | CCNA_03396 | trypsin-like serine protease, typically periplasmic, contains C-terminal PDZ domain | -0,774429 | 0,00147 | 0,0128453 |
| 252 | CCNA_00137 | hybrid two-component histidine kinase/receiver protein ShkA | -0,774168 | 0,0007 | 0,0067476 |
| 253 | CCNA_00984 | phosphinothricin N-acetyltransferase | -0,774093 | 0,00083 | 0,0078318 |
| 254 | CCNA_01221 | acyl carrier protein | -0,770191 | 0,00355 | 0,0273753 |
| 255 | CCNA_02222 | 5-methyltetrahydrofolate--homocysteine methyltransferase homocysteine-binding subunit | -0,76415 | 0,00078 | 0,0074088 |
| 256 | CCNA_R0066 | 23S RNA | -0,758802 | 0,00212 | 0,0176219 |
| 257 | CCNA_02606 | hybrid sensor histidine kinase/receiver domain protein | -0,751589 | 0,00301 | 0,0238718 |
| 258 | CCNA_00248 | sensor histidine protein kinase | -0,742028 | 0,00646 | 0,045581 |
| 259 | CCNA_02565 | hypothetical protein | -0,732791 | 0,00312 | 0,024568 |
| 260 | CCNA_00626 | methyl-accepting chemotaxis protein | -0,726941 | 0,00333 | 0,0260266 |
| 261 | CCNA_02061 | hypothetical protein | -0,726449 | 0,0024 | 0,0197616 |
| 262 | CCNA_02342 | hypothetical protein | -0,712639 | 0,0014 | 0,0122451 |
| 263 | CCNA_03045 | TadG-related pilus assembly protein | -0,710191 | 0,00289 | 0,0233196 |
| 264 | CCNA_03034 | hypothetical protein | -0,709776 | 0,00212 | 0,0176219 |
| 265 | CCNA_02843 | toxin protein parE-3 | -0,705253 | 0,00486 | 0,0356817 |
| 266 | CCNA_02507 | polyisoprenylphosphate hexose-1-phosphotransferase hfsE | -0,700272 | 0,00479 | 0,0353219 |
| 267 | CCNA_03038 | pilus assembly ATPase cpaE | -0,694468 | 0,00452 | 0,0336347 |
| 268 | CCNA_03713 | RNA polymerase sigma-54 factor rpoN | -0,693479 | 0,00163 | 0,0139929 |
| 269 | CCNA_01967 | hypothetical protein | -0,691658 | 0,00257 | 0,021023 |
| 270 | CCNA_03403 | hypothetical protein | -0,689745 | 0,00434 | 0,0327285 |
| 271 | CCNA_02223 | beta-lactamase, type II | -0,688964 | 0,00193 | 0,0162181 |
| 272 | CCNA_01193 | amylosucrase | -0,683499 | 0,002 | 0,0167317 |
| 273 | CCNA_01165 | hypothetical protein | -0,682255 | 0,00688 | 0,0476575 |
| 274 | CCNA_02402 | methyl-accepting chemotaxis protein | -0,678785 | 0,00161 | 0,0139004 |
| 275 | CCNA_01530 | flagellin | -0,678675 | 0,00315 | 0,0247677 |
| 276 | CCNA_01180 | penicillin acylase | -0,67762 | 0,00273 | 0,0221159 |
| 277 | CCNA_00985 | protease II | -0,671779 | 0,00246 | 0,0202042 |

| | | | | | |
|---|---|---|---|---|---|
| 278 | CCNA_00148 | hypothetical protein | -0,670019 | 0,0038 | 0,0290334 |
| 279 | CCNA_03194 | integral membrane protein | -0,662567 | 0,00491 | 0,0359661 |
| 280 | CCNA_00149 | transcriptional regulator, Cro/CI family | -0,660187 | 0,00461 | 0,0342187 |
| 281 | CCNA_01425 | H+ translocating pyrophosphatase | -0,645281 | 0,00329 | 0,0257277 |
| 282 | CCNA_00836 | flagellin | -0,64323 | 0,00723 | 0,0492411 |
| 283 | CCNA_03803 | cellulose biosynthesis protein CelD | -0,638073 | 0,00419 | 0,031697 |
| 284 | CCNA_01140 | sensory box/GGDEF family protein | -0,637155 | 0,00578 | 0,0416967 |
| 285 | CCNA_00589 | hypothetical protein | -0,630634 | 0,00639 | 0,0455141 |
| 286 | CCNA_02343 | hypothetical protein | -0,629967 | 0,0071 | 0,0485986 |
| 287 | CCNA_03222 | ring hydroxylating dioxygenase, alpha-subunit | -0,628857 | 0,00547 | 0,0397004 |
| 288 | CCNA_03061 | 3-oxoacyl-(acyl-carrier protein) reductase | -0,62103 | 0,00472 | 0,0348693 |
| 289 | CCNA_01625 | aminobenzoyl-glutamate utilization protein B | -0,61917 | 0,0066 | 0,0464727 |
| 290 | CCNA_02944 | hypothetical protein | -0,615087 | 0,00538 | 0,039262 |
| 291 | CCNA_00048 | S-adenosylmethionine synthetase | -0,607808 | 0,00541 | 0,0394684 |
| 292 | CCNA_03590 | hypothetical protein | -0,604046 | 0,00463 | 0,0343299 |
| 293 | CCNA_00279 | NAD(P)H dehydrogenase (quinone) | -0,600041 | 0,00549 | 0,0398095 |

Genes highlighted in green were identified in the ChIP-seq regulon in this work, and those in yellow the RNA-seq regulon presented by Stein *et al.* (2021).

### Supplementary methods

#### Construction of plasmids

pNPTS138-Δ*chvI*
Upstream and downstream regions of *C. crescentus chvI (CCNA_00237)* were amplified from WT gDNA by PCR respectively with primers 1013/1014 (690 bp) and 1015/1016 (710 bp). The PCR were then respectively digested with *Bam* HI/*Eco* RI and *Eco* RI/*Hind* III; and ligated into the pNPTS138 vector cut with *Hind* III and *Bam* HI.

pNPTS138-Δ*chvG*
Upstream and downstream regions of *C. crescentus chvIG (CCNA_00237-CCNA_00238* were amplified from WT gDNA by PCR respectively with primers 2173/2174 and 2563/2180. The PCR were then respectively digested with *Bam* HI/*Eco* RI and *Eco* RI/*Hind* III; and ligated into the pNPTS138 vector cut with *Hind* III and *Bam* HI

pNPTS138-Δ*chvIG*
Upstream and downstream regions of *C. crescentus chvIG (CCNA_00237-CCNA_00238* were amplified from WT gDNA by PCR respectively with primers 2173/2174 and 2179/2180. The PCR were then respectively digested with *Bam* HI/*Eco*

RI and *Eco* RI/*Hind* III; and ligated into the pNPTS138 vector cut with *Hind* III and *Bam* HI.

pNPTS138-Δ*chvG*$_{1\text{-}274}$
Upstream and downstream regions of *C. crescentus chvG*$_{1\text{-}274}$ *(CCNA_00237)* were amplified from WT gDNA by PCR respectively with primers 2173/3135 and 3136/2180. The PCR were then respectively digested with *Bam* HI/*Kpn* I and *Kpn* I/*Hind* III; and ligated into the pNPTS138 vector cut with *Hind* III and *Bam* HI.

pNPTS138-Δ*chvG*$_{274\text{-}534}$
Upstream and downstream regions of *C. crescentus chvG*$_{274\text{-}534}$ *(CCNA_00237)* were amplified from WT gDNA by PCR respectively with primers 2177/2773 and 2179/592. The PCR were then respectively digested with *Bam* HI/*Eco* RI and *Eco* RI/*Hind* III; and ligated into the pNPTS138 vector cut with *Hind* III and *Bam* HI.

pNPTS138-Δ*chvT*
Upstream and downstream regions of *C. crescentus chvT (CC3013*) were amplified from WT gDNA by PCR respectively with primers 2870/2871 and 2872/2873. The PCR products were then assembled with the pHR253 (pNPTS138) vector cut with *Bam* HI and *Hind* III via Gibson assembly.

pNPTS138-Δ*ntrX*
Upstream and downstream regions of *C. crescentus ntrX (CC1743*) were amplified from WT gDNA by PCR respectively with primers 648/649 (620 bp) and 650/651 (580 bp) and cloned into pSK. The pSK-648/649 and pSK-650/651 recombinant plasmids were then digested respectively with *Hind* III/*Eco* RI and *Eco* RI/*Bam* HI; and ligated into the pNPTS138 vector cut with *Hind* III and *Bam* HI.

pNPTS138-Δ*sigT*
Upstream and downstream regions of *C. crescentus CC3475* were amplified from WT gDNA by PCR respectively with primers 2055/2056 and 2057/2058. The PCR were then respectively digested with *Bam* HI/*Eco* RI and *Eco* RI/*Hind* III; and ligated into the pNPTS138 vector cut with *Hind* III and *Bam* HI.

pNPTS138-*chvI*$_{D53A}$
Upstream and downstream regions of *C. crescentus chvI (CC0237* or *CCNA_00237)* were amplified from WT gDNA by PCR respectively with primers 2584/2581 and 2580/2585. A single PCR fragment was generated by overlap extension PCR. Thereafter, the PCR fragment was cloned using blunt ligation with pNPTS138 vector cut previously with EcoRV.

pNPTS138-*chvI*$_{D53E}$
Upstream and downstream regions of *C. crescentus chvI (CC0237* or *CCNA_00237)* were amplified from WT gDNA by PCR respectively with primers 2584/2583 and 2582/2585. A single PCR fragment was generated by overlap extension PCR. Thereafter, the PCR fragment was cloned using blunt ligation with pNPTS138 vector cut previously with EcoRV.

pNPTS138-*chvG*$_{H309N}$

Upstream and downstream regions of *C. crescentus chvI (CC0237* or *CCNA_00237)* were amplified from WT gDNA by PCR respectively with primers 2577/2578 and 2576/2579. A single PCR fragment was generated by overlap extension PCR. Thereafter, the PCR fragment was cloned using blunt ligation with pNPTS138 vector cut previously with EcoRV.

pXC5-*chvI*

*chvI was amplified from WT gDNA by PCR primers* 2405/2406. *The PCR was then digested with Nde I and Kpn I, and ligated into the pXC-5 vector cut with the same restriction enzymes.*

pXC5-$chvI_{D52A}$

*chvI* phosphoablative mutant *was amplified from* the $chvI_{D52A}$ mutant *gDNA by PCR primers* 2405/2406. *The PCR was then digested with Nde I and Kpn I, and ligated into the pXC-5 vector cut with the same restriction enzymes.*

pXC5-$chvI_{D52E}$

*chvI* phosphomimetic mutant *was amplified from* the $chvI_{D52E}$ mutant *gDNA by PCR primers* 2405/2406. *The PCR was then digested with Nde I and Kpn I, and ligated into the pXC-5 vector cut with the same restriction enzymes.*

pXGFPC-2-*chvI*

*chvI* was amplified from WT gDNA by PCR primers 2157/2411. The PCR was then digested with *Nde* I and *Kpn I*, and ligated into the pXGFPC-2 vector cut with the same restriction enzymes.

pXGFPC-2-*chvG*

*chvG* was amplified from WT gDNA by PCR primers 2155/2568. The PCR was then digested with *Nde* I and *Eco* RI, and ligated into the pXGFPC-2 vector cut with the same restriction enzymes.

pXGFPN-2-*chvG*

*chvG* was amplified from WT gDNA by PCR with primers 2569/2156. The PCR were then digested with *Kpn* I and *Sac* I and ligated into the pXGFPN-2 vector cut with the same restriction enzymes.

pXGFPC-2-$chvG_{H309N}$

*chvG* was amplified from $chvG_{H309N}$ gDNA by PCR primers 2155/2568. The PCR was then digested with *Nde* I and *Eco* RI, and ligated into the pXGFPC-2 vector cut with the same restriction enzymes.

pXCHYC-5-*chvG*

*chvG* was amplified from WT gDNA by PCR primers 2155/2568. The PCR was then digested with *Nde* I and *Eco* RI, and ligated into the pXCHYC-5 vector cut with the same restriction enzymes

pXCHYC-5-*chvG*$_{1-274}$
*ChvG*$_{1-274}$ was amplified from WT gDNA by PCR primers 2155/3131. The PCR was then digested with *Nde* I and *Eco* RI, and ligated into the pXCHYC-5 vector cut with the same restriction enzymes.

pXCHYC-5-*chvG*$_{273-534}$
*chvG*$_{273-534}$ was amplified from WT gDNA by PCR primers 3132/2568. The PCR was then digested with *Nde* I and *Eco* RI, and ligated into the pXCHYC-5 vector cut with the same restriction enzymes.

pXCHYC-5-*chvG*$_{1-114}$
*chvG*$_{1-114}$ was amplified from WT gDNA by PCR primers 2155/3133. The PCR was then digested with *Nde* I and *Eco* RI, and ligated into the pXCHYC-5 vector cut with the same restriction enzymes.

pMR15-P*chvI*
P*chvI* was amplified from WT gDNA by PCR with primers 3170/3171. The The PCR product was then digested with *Xba* I/*Xho* I and ligated into the pMR15 vector cut with the same restriction enzymes.

pMR15-P*dipM*
P*dipM* was amplified from WT gDNA by PCR with primers 3174/3175. The The PCR product was then digested with *Xba* I/*Xho* I and ligated into the pMR15 vector cut with the same restriction enzymes.

pMR15-P*ftsN*
P*ftsN* was amplified from WT gDNA by PCR with primers 3176/3177. The The PCR product was then digested with *Xba* I/*Xho* I and ligated into the pMR15 vector cut with the same restriction enzymes

pMR15-P*sigT*
P*sigT* was amplified from WT gDNA by PCR with primers 3182/3183. The PCR product was then digested with *Xba* I/*Xho* I and ligated into the pMR15 vector cut with the same restriction enzymes

pMR15-P*phyR*
P*phyR* was amplified from WT gDNA by PCR with primers 3184/3185. The PCR product was then digested with *Xba* I/*Xho* I and ligated into the pMR15 vector cut with the same restriction enzymes.

pET-28a-*chvI*
*chvI* (*CCNA_00237*)was amplified from WT gDNA by PCR with primers 2157/2158 (~710 bp) , digested respectively with *Nde* I and *Sac* I, ligated into the pET-28a vector cut with the same restriction enzymes.

**Supplementary data references.**

Casadaban, M. J., & Cohen, S. N. (1980). Analysis of gene control signals by DNA fusion and cloning in *Escherichia coli*. *Journal of molecular biology*, *138*(2), 179-207.

Gober, J. W., & Shapiro, L. (1992). A developmentally regulated Caulobacter flagellar promoter is activated by 3'enhancer and IHF binding elements. *Molecular Biology of the Cell*, *3*(8), 913-926.

Stein, B. J., Fiebig, A., & Crosson, S. (2021). The ChvG-ChvI and NtrY-NtrX Two-Component Systems Coordinately Regulate Growth of Caulobacter crescentus. *Journal of Bacteriology*, *203*(17), e00199-21.

Stephens, C., Christen, B., Fuchs, T., Sundaram, V., Watanabe, K., & Jenal, U. (2007). Genetic analysis of a novel pathway for D-xylose metabolism in *Caulobacter crescentus*. *Journal of bacteriology*, *189*(5), 2181-2185.

Thanbichler, M., Iniesta, A. A., & Shapiro, L. (2007). A comprehensive set of plasmids for vanillate-and xylose-inducible gene expression in *Caulobacter crescentus*. *Nucleic acids research*, *35*(20), e137-e137.
